# A protein-DNA surface hydrogel mechanically reinforces the cell nucleus and protects the genome

**DOI:** 10.1101/2025.05.21.655270

**Authors:** Ramesh Adakkattil, Pranay Mandal, Yahor Savich, Valentin Ruffine, Mareike A. Jordan, Henrik Dahl Pinholt, Elisabeth Fischer Friedrich, Frank Jülicher, Stephan W Grill, Alexander von Appen

## Abstract

The nuclear envelope protects the genome from mechanical stress during processes such as migration, division, and compression^1–6^, but how it buffers forces at the scale of DNA remains unclear. Here, we utilize optical tweezers to show that a multivalent protein–DNA co-condensate containing the nuclear envelope protein LEM2^7,8^ and the DNA-binding protein BAF ^9,10^ shield DNA beyond its melting point at 65 pN^11^. Under load, their collective assembly induces an unconventional DNA stiffening effect that provides mechanical reinforcement, dependent on the intrinsically disordered region (IDR) of LEM2. At the nuclear surface, these components form an elastic surface hydrogel in which LEM2 IDR-IDR interactions contract the surface hydrogel relative to its relaxed state, introducing a pre-stress in the lamin network. Inside cells, this surface hydrogel model can recapitulate elastic properties of the nuclear envelope measured via AFM indentation experiments as well as nuclear morphology, using parameters obtained at the molecular scale by use of optical tweezers. Disruption of the surface hydrogel increases DNA damage and micronuclei formation during nuclear deformation. These findings reveal a load-bearing, mesoscale surface hydrogel that reinforces the nucleus and expands the functional repertoire of biomolecular condensates to include DNA protection under mechanical stress.

## Main Text

Mechanical stress can lead to DNA damage, such as double-strand breaks or micronucleation (Extended Data Fig.1a), which are associated with aging-related diseases including cancer, cardiomyopathies, and neurodegeneration^12–18^. The nuclear envelope is a mechanical scaffold that safeguards the genome^5^, but how it protects the genome from mechanical stress-induced DNA damage is not well understood. The nuclear envelope is formed by a highly cross-linked network primarily composed of polymeric lamins, the outer and inner nuclear membranes (INM), and proteins that link the membrane to the encapsulated DNA^5^. Here, we focus our attention on BAF-LEM2, both essential for nuclear envelope reassembly and genome stability^6–9^, that directly connects chromatin to the inner nuclear membrane^19–25^. Understanding how DNA is organized by BAF and the membrane-anchored LEM proteins, such as LEM2, function together can shed light on fundamental principles of nuclear functions and architecture. Notably, LEM2, along with other inner nuclear membrane proteins, contain substantial intrinsically disordered regions (IDRs) that protrude into the nucleoplasm, accounting for an estimated 30% of the total protein mass at the inner nuclear membrane (Fig. 1a, Extended Data Fig.1b, Extended Data Table 1). Proteins with extended IDRs can undergo phase separation to form biomolecular condensates^26,27^ and even hydrogels^28,29^. LEM2 undergoes phase separation^7^ to orchestrate nuclear membrane assembly during mitosis, driven by its amino-terminal IDR (LEM2_NTD_). Recent evidence suggests that protein-DNA co-condensation can generate forces that contribute to biomolecular organization^30–32^. Hence, we here set out to investigate potential protective functions of an extended IDR meshwork built by phase separating proteins at the nuclear surface.

**Table 1:** Copy numbers and average spacing of INM proteins. *Dimers confined only to the nuclear surface (Extended Data Fig. 5i).

| INM Proteins | Copy number | Average spacing (nm) |
| --- | --- | --- |
| BAF dimers* | $3.64 \times 10^5$ | 37 |
| LEM2 | $8.20 \times 10^4$ | 79 |
| Other LEM proteins | $2.14 \times 10^6$ | 15 |
| Lamin A/C | $3.60 \times 10^5$ | 38 |
| Lamin B dimers | $1.71 \times 10^6$ | 17 |

**Fig. 1:**
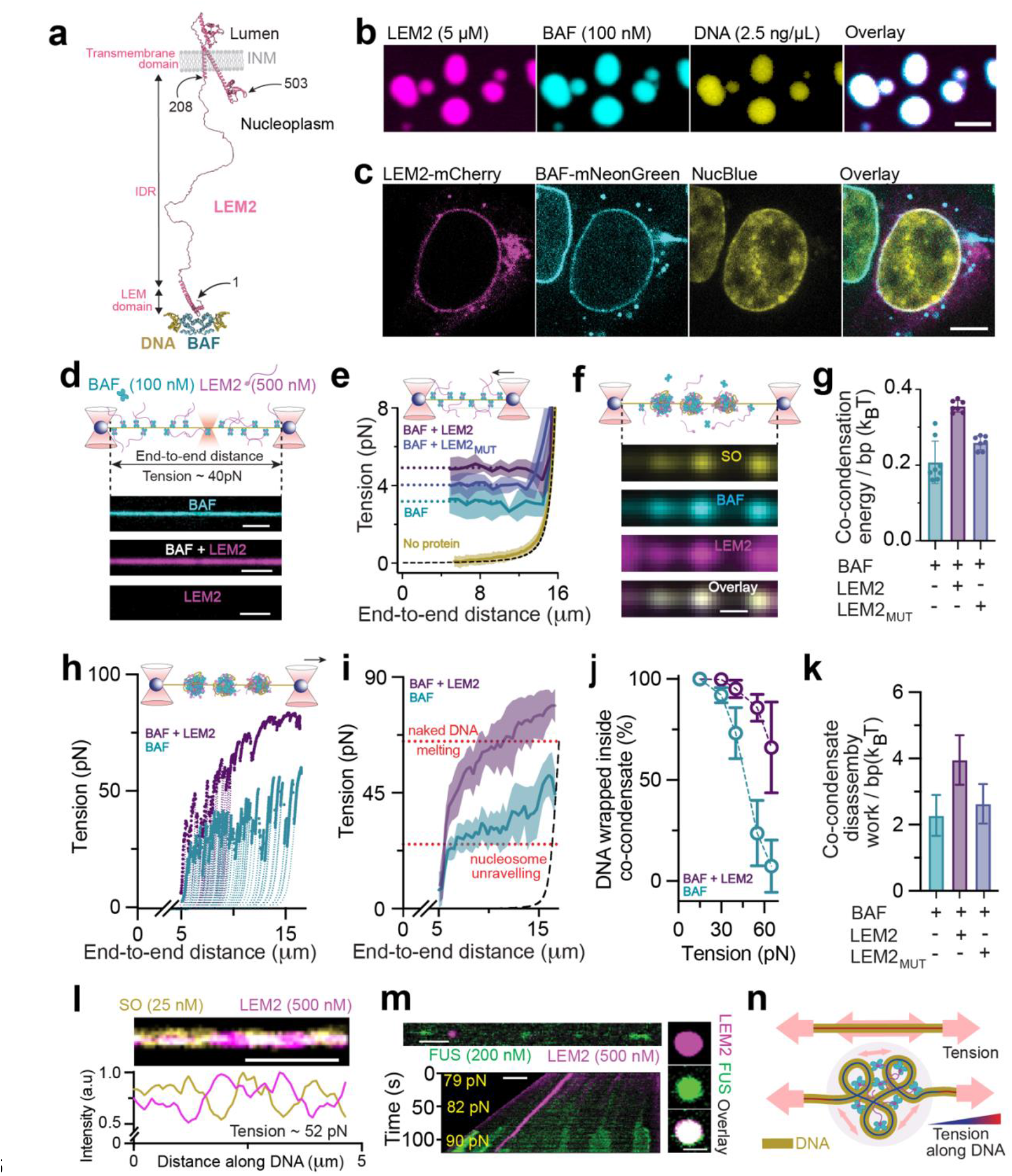
Co-condensation with BAF-LEM2 shields DNA beyond the melting transition. **a**, DNA-BAF-LEM2 interfaces the chromatin surface with the INM. Numbers indicate amino acid positions. **b**, Confocal imaging reveals bulk phase separation of reconstituted DNA-BAF-LEM2 (methods). **c**, Confocal imaging of BAF, LEM2 and DNA in the nucleus shows enrichment at the nuclear envelope (methods). **d**, Top, schematic of the optical tweezers assay where a single λ-DNA molecule is held between two optically trapped beads (large red cones with spheres) via biotin-streptavidin interactions at a tension of ~40 pN. Confocal imaging (small red cone) reveals BAF and LEM2 association to the stretched DNA. Bottom, confocal images of BAF (top), BAF and LEM2 (middle) and LEM2 (bottom) association to stretched λ-DNA. **e**, Average tension vs. end-to-end distance graphs obtained with the optical tweezer by gradually reducing DNA end-to-end distance from 16.7 µm to 5 µm (top inset) reveals a constant tension plateau indicative of protein-DNA co-condensation of DNA with BAF (n = 8), with BAF+LEM2_MUT_ (n = 7), and with BAF+LEM2 (n = 7; methods). Solid colored lines, average tension; colored shaded regions, mean ± standard deviation; dotted lines, extrapolations of tension to zero end-to-end distance; dashed black line, worm-like-chain (WLC) model of λ-DNA elasticity (methods). **f**, Schematic representation (top) and representative example from (**e**) (bottom) of DNA-BAF-LEM2 co-condensates, DNA (yellow) is labelled with Sytox Orange (SO). Scale bar, 0.5 µm. **g**, Co-condensation energy per DNA base pair (bp) obtained by integration of the tension vs. distance curves in (e). Co-condensation energy is significantly higher for DNA-BAF-LEM2 than for DNA-BAF and DNA-BAF-LEM2_MUT_. **h**, Representative tension vs. end-to-end distance graphs (solid dots) obtained by gradually increasing DNA end-to-end distance from 5 µm to ~16.5 µm (top inset) reveals characteristic ‘rip patterns’, with each rip unraveling extended amounts of DNA, during DNA-BAF and DNA-BAF-LEM2 co-condensate tension-induced disassembly. Dotted lines, WLC fits (methods) reveal the contour length of DNA inside co-condensates (see Fig. 1j). **i**, Corresponding average tension vs. end-to-end distance graphs for DNA-BAF (n = 10), and DNA-BAF-LEM2 (n = 10; methods) co-condensate tension-induced disassembly. Solid colored lines, average tension; colored shaded regions, mean ± standard deviation. DNA-BAF co-condensates disassemble at tensions slightly higher than those required to dissociate nucleosomes from DNA, while disassembling DNA-BAF+LEM2 co-condensates requires tensions above the critical melting tension of 65 pN. Dashed black line, WLC model of λ-DNA. **j**, Fraction of DNA remaining co-condensed during tension-induced disassembly as a function of applied tension. Note that 66±22 % of DNA remains co-condensed with BAF+LEM2 at the critical melting tension of 65 pN. **k**, Co-condensate per-base pair disassembly work obtained by integration of the tension vs. distance curves in Fig. 1i (BAF and BAF-LEM2) and Extended data Fig. 3e,f (BAF-LEM2_MUT_; n=5). **l**, Top, confocal image of SO, bright in directions of high DNA tension and dim in regions of low DNA tension, at 52 pN DNA tension reveals that DNA tension is higher outside co-condensates than inside. Bottom, corresponding profiles of normalized intensities showing alternate low and high distributions. **m**, Left, representative image (top) and kymograph (bottom) of DNA-BAF-LEM2 co-condensate disassembly under high tension reveals that the ssDNA binding protein FUS (green) exclusively associates with non-co-condensed regions, revealing that DNA inside co-condensates is protected from melting. Right, FUS partitions into LEM2 condensates in bulk, indicating that FUS is not excluded from DNA-BAF-LEM2 co-condensates. **n**, Sketch of the IDR-dependent mechanism of co-condensate tension buffering. DNA tension (pink arrows) is reduced within co-condensate (bottom) as compared to without (top), since inside tension redistributes through IDR-IDR and other molecular interactions. Arrow thickness indicates relative tension strength, and direction shows the path of redistribution. The color bar represents tension magnitude along the DNA. Scale bar, 5 µm in (**b,c**),(**m**)**-**right and 2 µm in (**d,l**), (**m**)**-**left.

### Protective DNA-protein co-condensation

We first set out to study the effects of BAF and LEM2 on DNA, and investigate the mechanical properties by use of optical tweezers. We established a minimal reconstitution system from purified components (full-length BAF and the N-terminal domain of LEM2 containing the IDR but lacking the transmembrane domain; Fig. 1a, Extended Data Fig.1c-f) and tested if bulk phase separation can be observed, consistent with previous observation^7^ (Extended Data Fig.1k). We found that phase separated LEM2 droplets enrich BAF and DNA *in vitro* (Fig. 1b, Extended Data Fig.1g). Since DNA, BAF and LEM2 enrich at the nuclear periphery in cells (Fig. 1c, Extended Data Fig.1h-j), it is possible that liquid-liquid phase separation and protein-DNA co-condensation contributes to the organization of the nuclear periphery.

We next set out to understand how BAF and LEM2 interact with individual DNA polymers. To this end, we added fluorescently labelled BAF and LEM2 to individual molecules of DNA stretched in the optical tweezer (methods, Extended Data Fig.2a,b) and followed the association of both proteins to DNA (Fig. 1d, Extended Data Fig.2d, Supplementary Video 1,2). We find that BAF homogeneously coats DNA in an apparently sequence-independent manner^33^. While we did not observe binding for LEM2 alone (500 nM; Fig. 1d, Supplementary Video 3), DNA-BAF caused instant LEM2 recruitment into a homogenous film. We conclude that BAF recruits LEM2^7^ to DNA at physiologically relevant concentrations.

Given that LEM2 can undergo phase separation in bulk^7^, we next set out to investigate if BAF-LEM2 can co-condense together with DNA^31^. We first focused on BAF alone, and used the optical tweezer to stretch a single molecule of λ-DNA to 16.7 µm end-to-end-distance (corresponding to ~40 pN of DNA tension, methods). We next allowed BAF to bind, and then gradually reduced DNA end-to-end distance. In all cases we observed that DNA tension reduces in the process, but plateaus at an average of 3.3±0.8 pN (Fig. 1e, Extended Data Fig.2f,g, methods) despite further reduction of the end-to-end distance. The occurrence of a constant tension plateau is indicative of a collective phase transition towards DNA-BAF co-condensate formation, similar to a ‘sticky polymer’ undergoing a globular collapse^31^ (Fig. 1f, Extended Data Fig.2e, Supplementary Video 4). We measured the associated bulk condensation energy to be 0.2±0.1 k_B_T/bp (methods, Fig. 1g, Extended Data Fig.2j,k,n). Adding 500nM LEM2, significantly below the saturation concentration^34^ of LEM2 for bulk phase separation (4.0 µM, Extended Data Fig.1k,l), results in co-condensate formation at significantly increased plateau tension and with significantly increased bulk condensation energy (4.9±0.6 pN and 0.4±.02 k_B_T/bp, respectively; Fig. 1e-g, Extended Data Fig.2h, Extended Data Fig.2l-n, methods, Supplementary Video 5). Instead, adding a variant of LEM2 that is unable to undergo bulk phase separation but still binds BAF (LEM2_MUT_, Extended Data Fig.1f) results in only a marginal increase of both plateau tension and bulk condensation energy (4.1±0.5 pN and 0.3±.02 k_B_T/bp, respectively; Fig. 1e,g). Taken together, DNA cross-bridging by BAF dimers alone can result in protein-DNA co-condensation. LEM2 stabilizes the DNA-BAF interaction, with a significant increase of the per basepair bulk co-condensation energy to a value that is ~20% of typical DNA basepair stacking energies^35^. We conclude that BAF-LEM2 can co-condense together with DNA.

Given that the bulk co-condensation energy is significant, one might expect DNA-BAF-LEM2 co-condensates to be stable and protect DNA from force-induced damage. To evaluate this possibility, we set out to unravel the co-condensates by force while investigating DNA integrity. We first focused on BAF alone. After forming a DNA-BAF co-condensate as described above, we gradually increased the end-to-end distance of DNA and recorded the tension applied (Fig. 1h,i; methods). In all cases we observed that DNA tension increases in the process, to an average disassembly tension of 34±9 pN as the co-condensate unravels through many consecutive ‘ripping events’ each releasing extended amounts of DNA (Fig. 1h; methods, disassembly tension is independent of pulling speeds for speeds between 250 nm/s and 10 nm/s, Extended Data Fig.3a). We note that co-condensate disassembly tension is higher than the tension that needs to be applied to mechanically evict nucleosomes from DNA^36,37^, indicative of strong DNA-BAF interactions. Concurrently, we measured the associated co-condensate disassembly work to be 2.3±0.7 k_B_T/bp (Fig. 1k). Notably, the disassembly work of DNA-BAF-LEM2_MUT_ co-condensates is not significantly increased as compared to DNA-BAF co-condensates (2.6±0.6 k_B_T/bp, Extended Data Fig.3e,f). However, adding the phase-separating variant of LEM2 results in significantly increased DNA-BAF-LEM2 co-condensate disassembly tension and disassembly work (47±8 pN and 3.9±0.8 k_B_T/bp, respectively; Fig. 1i,k), as well as a less pronounced ‘rip pattern’ (Fig. 1h). Remarkably, in all experiments involving BAF-LEM2, tension during unraveling surpasses the critical tension of 65 pN^11^ required to melt DNA. Hence, we find that 66±22% of the DNA remains localized within the co-condensate when the melting tension of 65 pN is crossed for the first time, compared to 8±13% for DNA-BAF co-condensates (Fig. 1j). We conclude that, generally, disassembly tension during co-condensate unraveling is more than ~10x larger than plateau tension during co-condensate formation, and the per-base pair disassembly work is ~10x larger than the per-base pair bulk condensation energy (Fig. 1e,g,i,k). Such hysteresis arises when systems are unable to equilibrate on the experimental timescale (Extended Data Fig.3b-d; methods) particularly when kinetic barriers during formation and dissolution are different, and is often seen in systems that undergo a mesoscale phase transition^38^. Taken together, BAF-LEM2-DNA co-condensates are stable gel-like structures that can retain DNA when subjected to external forces larger than those required to melt DNA.

Is the DNA inside these co-condensates mechanically protected? We hypothesize that, in analogy to hydrogels in material science^39^, cross-bridges and multivalent interactions formed by the IDRs of LEM2 function to redistribute mechanical tension, effectively lowering the tension that is experienced by DNA (Fig. 1n). To test if DNA tension inside DNA-BAF-LEM2 co-condensate is lower than DNA tension outside, we applied the tension-sensitive intercalating DNA dye Sytox Orange^40^ (SO, methods). We find that at high external tension (52 pN), the intensity of SO is significantly higher outside the co-condensates (Fig. 1l, Extended Data Fig.3g,h; methods) than inside, consistent with DNA experiencing lower tension inside co-condensates as compared to outside. Furthermore, addition of the single-stranded DNA (ssDNA) sensor, FUS^31,41^(Extended Data Fig.3i-k), reveals that at tensions up to ~85pN, DNA only melts outside the co-condensate and not inside (Fig. 1m, methods, Supplementary Video 6). We conclude that external tensions that are applied to DNA-BAF-LEM2 co-condensates likely percolate and redistribute throughout this gel-like structure, leading to an overall reduction of DNA tension inside. Therefore, protein-DNA co-condensates formed with multivalent protein-protein interactions can protect DNA from force-induced melting.

### A reinforcing mechanical response

Multiple studies have reported that when cell nuclei are mechanically deformed and undergo nuclear blebbing, rupture, or micronucleation, BAF and LEM2 rapidly associate to DNA that is mechanically stressed, exposed, or even expelled to the cytoplasm, likely to revert the damage^9,12,23,24,42,43^. We therefore wondered if the *de-novo* assembly process of the BAF-LEM2 module on naked, mechanically stressed DNA imparts a more active role in protecting DNA. To mimic such a re-assembly scenario under mechanical stress, we again stretched a single molecule of λ-DNA to 16.7 µm end-to-end-distance (corresponding to ~40 pN of DNA tension), but now monitored changes in DNA tension (by a value ΔT) during module assembly in a constant trap position protocol (methods). Notably, we observed that in the first 100 seconds after addition of BAF-LEM2, the beads in both optical traps are pulled inwards, due to a pulling tension generated by the module assembly process that opposes the external tension applied by the two traps (Fig. 2a). This resulted in a DNA tension increase by ΔT = 7.5±0.9 pN, despite the high initial DNA tension of 40 pN (Fig. 2b, methods). We note that neither the addition of BAF alone, nor adding BAF together with the LEM2_MUT_ resulted in a significant tension increase (Fig. 2b, ΔT = −0.2±0.9 pN for BAF; ΔT = 1.8±0.7 pN for BAF-LEM2_MUT_). Finally, we find that ΔT increases with increasing LEM2 concentration, and that, remarkably, ΔT increases in an apparently linear fashion with initial DNA tension (Fig. 2c). The module thus displays features of a reinforcing response: the higher the mechanical load on DNA, the more this module pulls back while re-assembling.

**Fig. 2:**
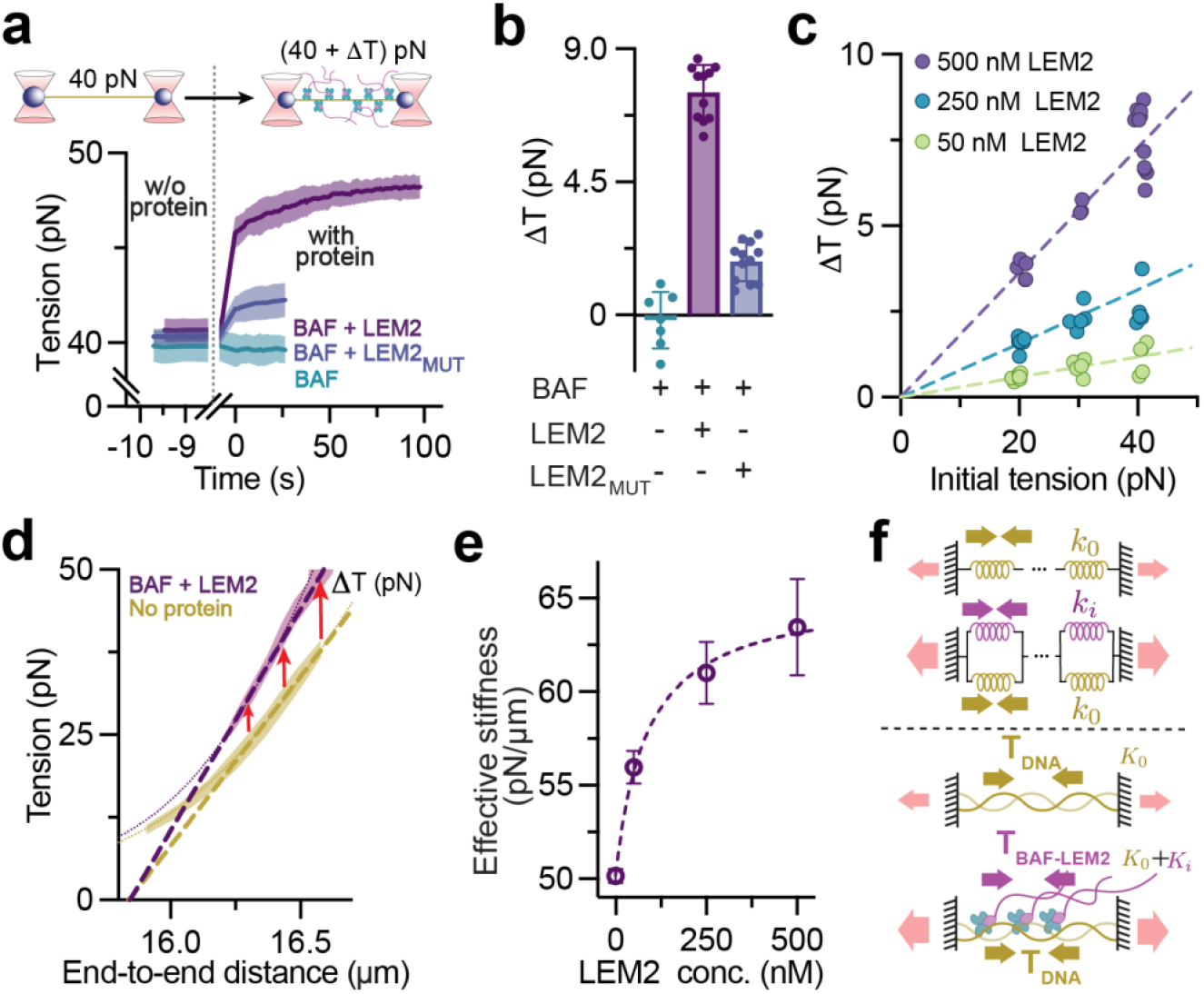
BAF-LEM2 assembly on DNA generates a reinforcing response. **a**, Top, DNA stretched to an initial ~40 pN tension is subjected to a buffer exchange from protein-free (left) to protein-containing (right) at time t = 0s while trap positions are maintained. Bottom, tension increases with time post addition of 100nM BAF + 500nM LEM2 (BAF+LEM2) and 100nM BAF + 500nM LEM2_MUT_ (BAF+LEM2_MUT_) due to module assembly on DNA, but does not increase post addition of 100nM BAF alone. Solid colored lines, mean tension; colored shaded regions, mean ± standard deviation (BAF: n=7, BAF-LEM2: n=11, BAF-LEM2_MUT_: n=12). **b**, Corresponding average tension increases ΔT. **c**, ΔT as a function of initial DNA tension and LEM2 concentration (100nM BAF, n=20 for 500nM LEM2, n=17 for 250nM LEM2, : n=19 for 50nM LEM2). Dashed lines, linear fits. **d**, Direct experimental characterization of BAF-LEM2 module compliance in the enthalpic regime of DNA stretching via average tension vs. DNA end-to-end distance graphs in absence (yellow shaded region, mean ± standard deviation, n=11) and in presence (magenta shaded region, mean ± standard deviation, n=11) of 100nM BAF and 500nM LEM2. Dotted lines, WLC model of λ-DNA. Linear fits (dashed lines) in the enthalpic regime (methods) reveal a stiffness but not a rest-length change between the two conditions (see main text). Red arrows illustrate tension increase ΔT when keeping DNA end-to-end distance fixed. **e**, Effective stiffness as a function of LEM2 concentration (100nM BAF) measured via constant tension experiments (Extended Data 4c, methods). Error bars, mean ± standard deviation; dashed line, fit to the stiffening reinforcement model. **f**, Sketch of the IDR-dependent mechanism of assembly-induced stiffening reinforcement. Top, DNA, represented as a series of springs (stiffness k_o_), is held in place at fixed end-to-end distance. BAF-LEM2 module assembly results in the addition of springs in parallel (stiffness k_i_). Saturated BAF-LEM2 binding increases the effective stiffness of the system from K_0_ (DNA alone) to K_0_ + K_?_. Bottom, Saturated BAF-LEM2 binding and compliant element assembly generates tension T_BAF-LEM2_ (magenta arrows) in addition to DNA tension T_DNA_ (yellow arrows), increasing the total tension experienced by the system (pink arrows).

We next set out to shed light on the mechanism that underlies this reinforcing response. The linear dependence of tension increase ΔT on initial tension suggests that LEM2 and its IDR interactions introduce an additional compliant element. We therefore set out to determine the compliant properties (i.e. effective stiffness and rest length, Extended Data Fig. 4a) of the combined DNA-BAF-LEM2 module in comparison to naked DNA alone. We applied an initial tension between 20 and 40 pN, added BAF and LEM2, and then gradually increased extension while recording tension to measure the effective stiffness of the entire assembly (Fig. 2d, Extended Data Fig.4b). We find a linear relationship in the regime between 20 - 45 pN, with an effective stiffness ~1.3±0.1 fold higher than for naked DNA (assuming saturation, K_0_+K_i_ = 67±3 pN/µm; compared to K_0_ = 51±2 pN/µm, Fig. 2d). Notably, the effective stiffness increases in a saturating manner with increasing LEM2 concentration (Fig. 2e, Extended Data Fig.4f). We obtained an effective rest length by extrapolating the linear fit to vanishing tension (Fig. 2d, Extended Data Fig. 4b,c; methods), and found no significant difference between the two conditions (15.84±0.02 µm without BAF-LEM2 module, compared to 15.84±.01 µm with BAF-LEM2 module). Together, these results suggest a picture where LEM2 IDR-IDR interactions generate compliant elements of identical rest length assembled in parallel to DNA (Fig. 2f). We capture this mechanism in a theoretical description that considers DNA mechanics in the enthalpic regime together with module assembly energetics (stiffening reinforcement model, Supplementary Note 1): DNA is described as a chain of spring elements, and the assembly of BAF-LEM2 modules results in the inclusion of additional spring element in parallel and without a rest length change (Supplementary Note 1, methods, Extended Data Fig. 4c-f). This description accounts for the dependence of stiffness on concentration (Fig. 2e, Extended Data Fig.4e,f) as well as the measured dependencies of ΔT on initial tension and LEM2 concentration. We note that assembling compliant elements in parallel generates a reinforcing response, as can be seen in Fig. 2d when keeping the end-to-end distance fixed: the higher the initial tension, the higher the tension increase upon module assembly (red arrows). This reveals a previously undescribed protein-induced reinforcing response, which, if operating at the surface of the cell nucleus, would significantly impact cellular nuclear mechanics.

### The impact on nuclear surface mechanics

We next set out to explore whether the reinforcing property of LEM2 manifests at the surface of the nucleus in vivo (Fig. 1b,c; Extended Data Fig. 1h-j). To this end, we examined the morphology of the protein–DNA interface at the INM in cryo-electron tomograms of focused ion beam milled cellular thin sections. Across datasets from different cell types^44–46^, we consistently observed an amorphous, ~14 nm thick layer directly beneath the INM (Extended Data Fig. 5a-e). Notably, we find that this layer excludes nucleosomes, revealing a nucleosome exclusion zone at the INM. In agreement with previous studies, this region contains lamins^47–49^ (Extended Data Fig. 5c,d). While DNA cannot be directly visualized in these tomograms, independent work suggests that this layer is enriched in DNA^50^. Given its position and composition, the zone includes BAF and the transmembrane protein LEM2^51–53^. These findings support the existence of a gel-like, chromatin-enveloping layer at the INM that potentially contributes to the mechanical reinforcement of the nuclear surface (Fig. 1b,c; Extended Data Fig. 1h–j).

We next investigated if a) the material properties of this layer at the nuclear surface and thereby b) nuclear morphology are affected when the LEM2-dependent reinforcing response is impaired. We first evaluated if the effective area elastic modulus of the nuclear surface of the cells is changed when LEM2 amounts are reduced by ~74% (from 8.2 × 10^4^ to 2.1 × 10^4^ molecules per cell, Extended Data Fig. 6c-e, methods) using RNAi (siLEM2), by using an AFM in conjunction with confocal imaging (Fig. 3a,b, Extended Data Fig. 10e-g) to quantify the mechanical work (Fig. 3c) associated with an increase of nuclear surface area (Extended Data Fig. 10b, methods). To probe nuclear surface properties, we indented nuclei close to the plasma membrane of latrunculin-A treated cells using an AFM (Extended Data Fig. 10a, methods) cantilever, and find that LEM2 depletion led to a 19.6 ± 9.2% reduction in the area elastic modulus (from 1.84 ± 0.15 pN/nm to 1.48 ± 0.12 pN/nm; Fig. 3d). We next evaluated if nuclear morphology is changed when LEM2 amounts are reduced (Fig.3e, Extended Data Fig. 6f-h). We find that reducing LEM2 amounts by ~74% (methods) leads to a 5.4 ± 2.0% increase of nuclear surface area (from 516±6 µm^2^ to 544±8 µm^2^, Extended Data Fig. 6i,j) and a 28.6 ± 8.0% increase in nuclear asphericity (from 0.0700±0.0020 to 0.0903±0.0054, Fig. 3f bottom; asphericity defined as *A*/*Av*−*1*, where *Av* is the surface area of a sphere with equivalent nuclear volume). We note that we obtain qualitatively similar results when impairing the LEM2-dependent reinforcing response via BAF RNAi (siBAF, Extended Data Fig. 6a,b, Extended Data Fig. 8) with more significant changes to both nuclear asphericity and area elastic modulus as compared to the LEM2 RNAi condition (Extended Data Fig. 8l, Extended Data Fig. 8c). We conclude that reducing either LEM2 or BAF amounts by RNAi results in a significant reduction in the effective area elastic modulus of the nuclear surface as well as a significant increase of nuclear asphericity.

**Fig. 3:**
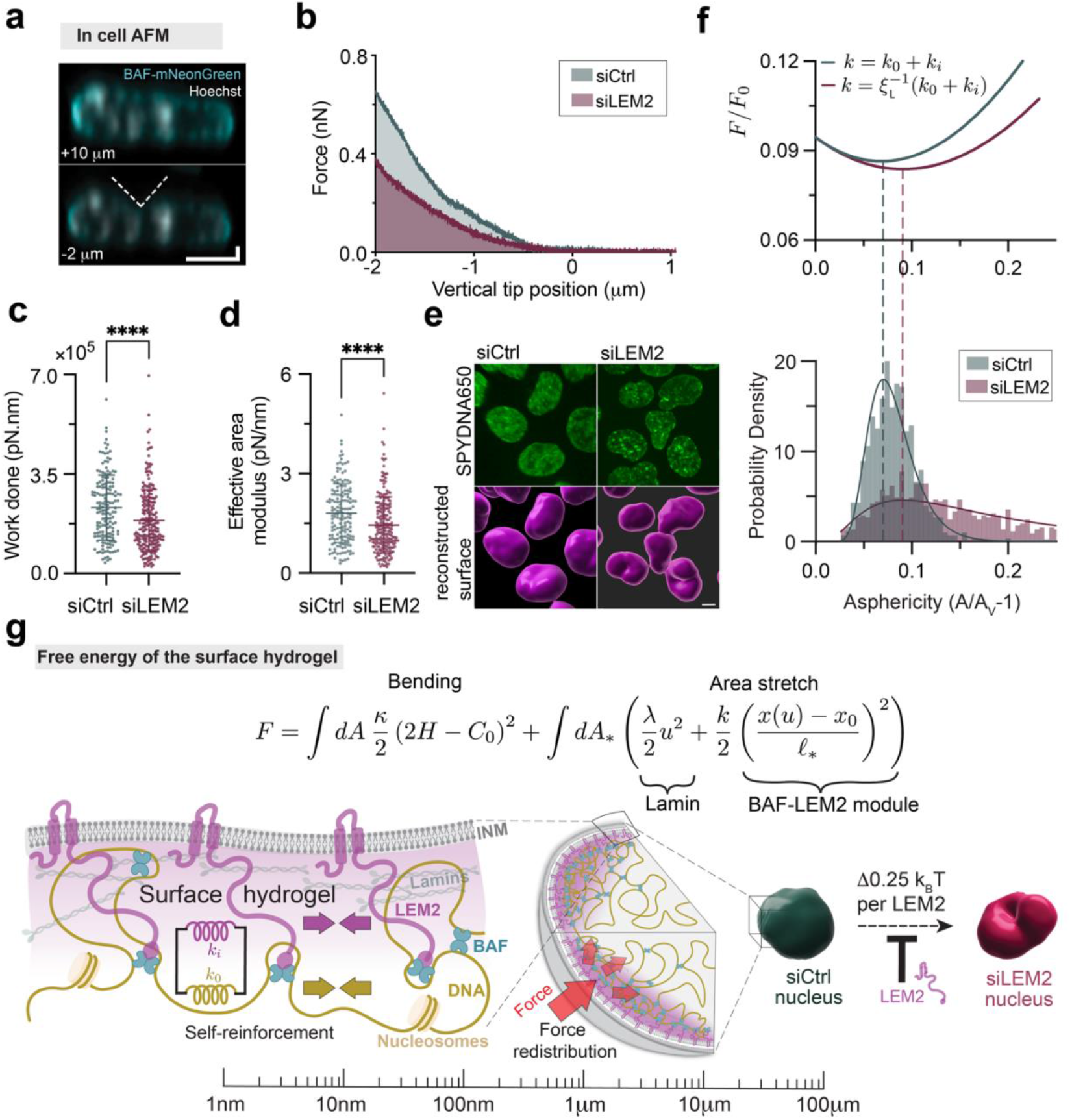
A cross-linked network of LEM2-BAF-DNA forms a surface hydrogel that reinforces the nuclear periphery to regulate its morphology and integrity. **a**, Representative live-cell images of AFM indentations of HCT-116 BAF-mNeonGreen cells; not indented (vertical tip position +10 µm) and indented (vertical tip position −2 µm). Dashed white line, illustration of pyramidal cantilever tip. Scale bar, 5 µm (horizontal and vertical). **b**, Representative AFM force-indentation curves of nuclei of latrunculin A-treated HCT-116 cells, for siCtrl and siLEM2 cells. The cantilever makes contact with the nucleus at a vertical tip position of 0 µm, and we measure the mechanical work (shaded areas under the curves) required to indent the nucleus by 2 µm. Indentation in absence of LEM2 requires less work than in presence. Segmentation of the nuclear surface reveals the nuclear surface area increase (Extended Data Fig. 10b), which together with the mechanical work (**c**) allows for determination of the effective area modulus shown in (**d**, methods). **c**, Mechanical work for a 2 µm indentation of the nucleus of siCtrl and siBAF cells (methods), from three biological replicates (siCtrl: n = 159, siLEM2: n = 225). **d**, Effective area modulus of nuclei of siCtrl and siLEM2 cells (Extended Data Fig. 10d; siCtrl: n = 159, siLEM2: n = 225, methods). **e**, Top, confocal image stacks of DNA (SPYDNA650) of siCtrl (left) and siLEM2 (right) cells. Bottom, 3D segmentations of the nuclear surfaces for determining nuclear volume, shape, and surface area. **f**, Top, plots of the normalized free energy (*F*/*F*_0_) of the REMM model (methods) as a function of nuclear asphericity reveal that the minimum for the siLEM2 condition with *k* = ξ_*L*_ ^−*1*^(*k*_*0*_ + *k*_*i*_) with measured stiffness ratio ξ_*L*_ (Supplementary Note 1) is placed at a larger asphericity than for the siCtrl condition with *k* = *k*_*0*_ + *k*_*i*_. Bottom, histograms of nuclear asphericity (measured as in **e** for siCtrl and siLEM2 cells (n = 603 and n = 901, respectively, pooled across three biological replicates). Colored lines, log-normal fit. With REMM model parameters λ = *0*.*014* pN/nm, *A*_∗_ = *1*.*8 A*_*v*_, *k* = *0*, rest length mismatch *x*_*0*_/*x*_∗_ = *0*.*75* and energy scale *F*_*0*_ = λ*A*_∗_. The histograms peak at the values that minimize the corresponding REMM surface hydrogel free energy *F* (vertical dashed lines). **g**, A nuclear surface hydrogel at the nuclear periphery. Top (theory), surface hydrogel free energy (*F*) is determined by integration over the strain-free area (*A*_∗_) of a bending energy term (left), with bending modulus *k* and average and spontaneous curvatures *H* and *C*_*0*_, a bare area elastic energy term (middle) representing lamins with bare elastic area modulus λ and surface strain *u*, and a DNA-BAF-LEM2 module term (right) with stiffness *k*, stretch relative to rest length *x*(*u*)−*x*_*0*_ with 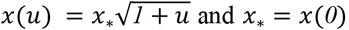 and *x*_∗_ = *x*(*0*), and surface density of DNA-BAF-LEM2 modules in the strain-free state 1/*l*_∗_^*2*^ (Supplementary Note 1). Bottom left (sketch, molecular scale), schematic of the REMM surface hydrogel (purple shade) formed below the INM (gray), comprising LEM2 (magenta), BAF (cyan), DNA (yellow), lamin filaments (green) and nucleosomes, with reinforcing properties as described (elastic properties indicated by coloured springs, tension indicated by colored arrows). Bottom middle (sketch, mesoscale), the surface hydrogel acts as a mechanical buffer, redistributing force along the nuclear periphery to maintain structural integrity under load. Bottom right (sketch, cellular scale), nuclear morphology is regulated by the integrity of the surface hydrogel. LEM2 depletion weakens the cross-linked shell, leading to reduced mechanical resistance and increased nuclear asphericity (red) compared to the more spherical shape in control cells (green).

### A nuclear surface hydrogel

How can a ~14 nm thick surface layer at the nuclear periphery, reinforced by LEM2, influence the mechanical properties of the entire nucleus at the cellular scale? To address this, we developed a mechanical model of the nuclear surface (Extended Data Fig. 5f) as a thin hydrogel layer. This model (Fig. 3g) quantitatively incorporates the DNA-stiffening effects observed in vitro (Fig. 2) along with measurements of nuclear surface mechanics and morphology in cells. We describe the nuclear envelope as a two-dimensional elastic surface hydrogel that encloses the nucleoplasm at constant volume. In the absence of BAF-LEM2, the surface hydrogel is characterized by a bending modulus (*κ*) and a bare area elastic modulus (λ). We capture the effect of the BAF-LEM2 module by introducing a surface density of additional elastic spring elements (Supplementary Note 1, Eq. 19). The bare area elastic modulus λ is largely determined by the lamin network^54^, and the elastic springs have the reinforcing properties described above (Supplementary Note 1, Eq. 36). The effect of an increase of nuclear asphericity upon BAF-LEM2 module impairment via LEM2 or BAF depletion (Supplementary Note 1) can be accounted for if the elastic spring elements associated with the BAF-LEM2 module contract the surface hydrogel relative to its relaxed state due to a rest-length mismatch between the two systems, introducing a pre-stress. When LEM2 or BAF levels are reduced, the elastic spring elements are removed and the surface hydrogel can relax, increasing its surface area and leading to more aspherical shapes (Fig. 3f bottom, Extended Data Fig. 6i,j, Extended Data Fig. 8j-l). We next used the AFM measurements (Fig. 3d, Extended Data Fig. 8c, Extended Data Fig. 10c,d) as well as the reinforcing properties of the BAF-LEM2 module characterized in the optical tweezer to quantitatively account for the observed nuclear shape changes upon either LEM2 depletion (Fig. 3f, top) or BAF depletion (Extended Data Fig. 8l top, Supplementary Note 1), using this REst-length Mismatched Multicomponent (REMM) surface hydrogel model (Supplementary Note 1). We use the rest length mismatch, the relaxed surface area as well as the bare area elastic modulus λ as parameters. Our analysis suggests that λ is ~25 times smaller than the effective area elastic moduli measured with the AFM in control cells. This suggests a soft lamin network providing mainly entropic elasticity^55^. Because the lamin network is soft, changing the stiffness of reinforcing BAF-LEM2 spring elements has a significant effect on the material properties of the REMM surface hydrogel. This model can therefore account for the observed changes in area elastic modulus and nuclear shape. We conclude that the REMM surface hydrogel model, taking into account a stiffening effect based on LEM2 IDR-IDR interactions characterized on the molecular scale, can capture nuclear surface properties at the cellular scale.

We next asked if the LEM2 component of this surface hydrogel imparts specific protective functions. We find that under compression (Extended Data Fig. 9a-b), cells depleted of LEM2 by RNAi accumulate significantly higher levels of DNA double-strand breaks (Extended Data Fig. 9c,d,g-l) and display a larger degree of micronucleation (Extended Data Fig. 9e,f, Extended Data Fig. 10h-k)^13,18^. We conclude that, consistent with our optical tweezer results and our expectations from theory, the LEM2 component, with its IDR’s, provides a specific protective function against mechanically induced damage.

## Discussion

Here, we have used optical tweezers and *in vitro* reconstitution to show that BAF and LEM2 form multivalent protein–DNA co-condensates. We reveal that i) these multivalent assemblies shield DNA beyond its melting threshold at 65 pN of tension, and ii) their assembly on DNA stretched close to melting, generates forces that counteract the stretch. Notably, a surface hydrogel model incorporating the second effect quantitatively recapitulates elastic properties of the nuclear envelope at the cellular scale, using parameters obtained at the molecular scale by use of optical tweezers. Our work paints a picture (Fig. 3g) where the inner nuclear membrane protein LEM2 and the DNA crosslinking factor BAF form a pre-stressed surface hydrogel together with the nuclear lamins that can dynamically resist mechanical stress applied to the nuclear periphery. This is achieved via a molecular reinforcing response based on LEM2’s IDR-IDR interactions, converting up to ~0.25 K_B_T of free energy per LEM2 molecule into mechanical work (methods). Given that there are no ATPases involved (Fig. 2), this free energy should derive from molecular-scale assembly processes but can impact large-scale nuclear organization and nuclear shape, thereby helping to preserve chromatin integrity (methods).

Together, this uncovers a novel design principle in which multivalent protein networks assemble on DNA polymers, generating reinforcement responses that buffer mechanical damage and enhance structural resilience; a strategy reminiscent of load-bearing tough hydrogels in material science^39,56^. Given the interconnected architecture of the inner nuclear membrane, this mechanism likely extends beyond BAF-LEM2 to include other LEM domain proteins^19^, IDR-containing inner nuclear membrane proteins such as lamins or lamin B receptor^54,57,58^, and even chromatin itself^9,51,59^. Lastly, the ability of DNA-protein co-condensation to generate biologically relevant forces suggests a broader dynamic role of condensates in shaping functional chromatin structures during processes such as transcription and replication^60–63^.

## Acknowledgments

We thank Jose Alberto Morin for the initial experiments on this system with the optical tweezer. We thank Tony Hyman, Anatol Fritsch and Juan Iglesias for help with phase separation saturation concentration determination, and Martin Beck and Reiya Taniguchi for sharing cryo electron tomography data. We thank the staff and students of the 2023 MBL Physiology course where important theory, initial experiments but no controls were performed, particularly A. Ortega Granillo, J. Brzostowski, N.King, and D. Fletcher. We thank the following services and facilities at MPI-CBG Dresden for their support: Protein Expression Facility, Biophysics Core facility, Scientific Computing Facility, Genome Engineering Facility and the Light Microscopy Facility.

## Funding

Funding by the Max Planck Society (AvA, SWG, FJ)

European Research Council-Organelloids 101117619 (AvA)

German Research Foundation SPP2191 (AvA)

Heineman Foundation through Minerva (AvA)

Volkswagen Foundation (FJ)

## Supplementary Information

Extended Data Figs. 1-10

Materials and Methods

Supplementary Note 1

Supplementary Note 2

Extended Data Table 1

Supplementary Video 1-6

