## Supplementary Information for "A protein-DNA surface hydrogel mechanically reinforces the cell nucleus and protects the genome"

**Extended Data Figs. 1-10**

**Materials and Methods**

**Supplementary Note 1**

**Supplementary Note 2**

**Extended Data Table 1**

**Supplementary Video 1- 6**

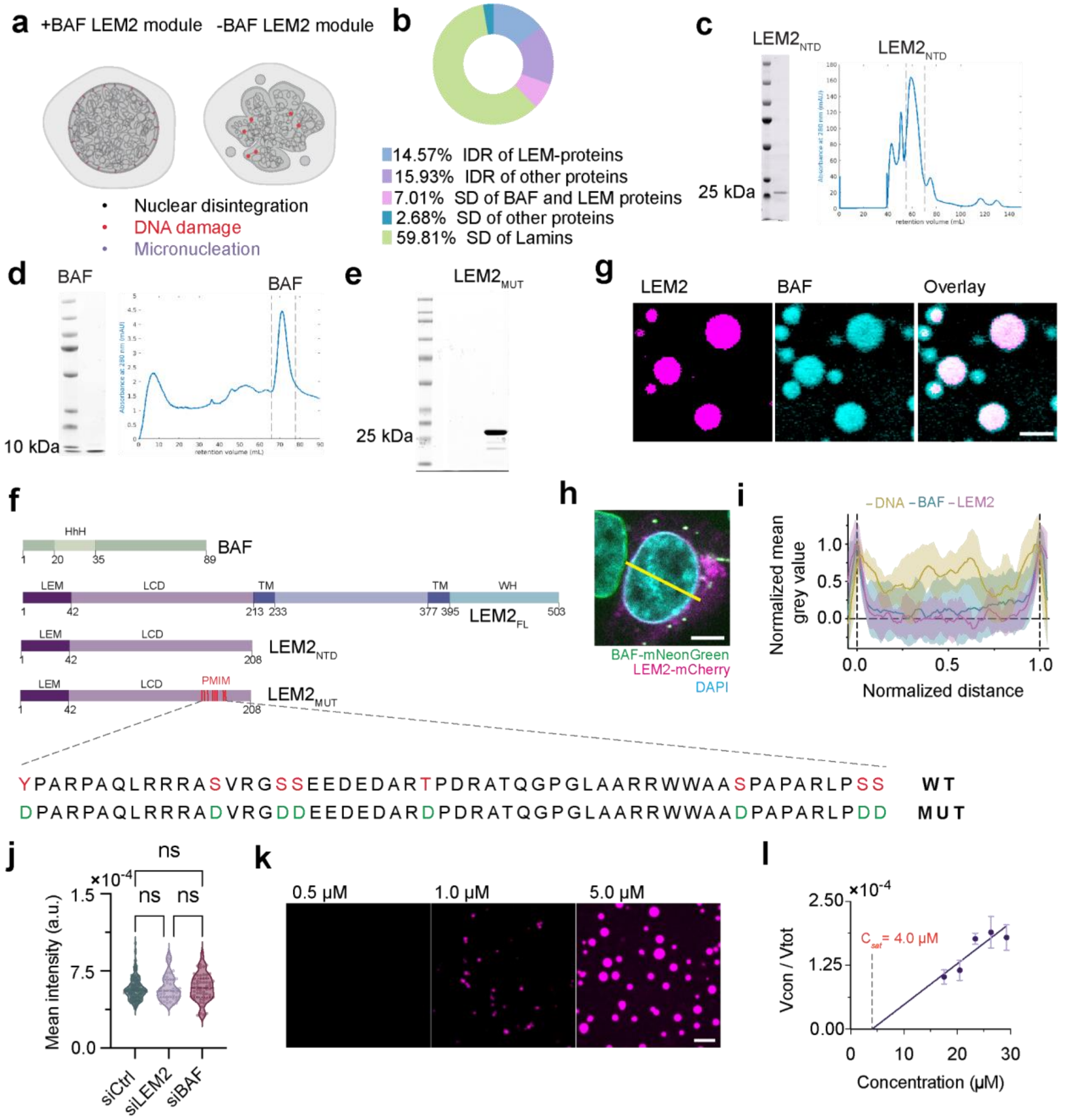


**Extended Data Fig. 1: A bottom-up reconstitution system for *in vitro* assembly of DNA-BAF-LEM2, with corresponding cellular visualization.**

**a,** Schematic representation of nuclear disruption phenotypes when BAF-LEM2 is depleted. **b,** Composition of the nuclear lamina, highlighting structured domains (SD) and intrinsically disordered regions (IDR) of its constituent proteins. SDS PAGE (left) and SEC Elution profile (right) of purified recombinant **c,** LEM2 **d,** BAF. **e,** SDS PAGE of purified recombinant $LEM2_{MUT}$. **f,** Domain chart showing protein domains of various recombinant constructs. Labels indicate domain names, and numbers represent amino acid positions. The red vertical lines indicate phosphomimetic mutations (PMIM; S,Y,T → D) introduced in LEM2 to abolish multivalent interactions. Full length LEM2 ($LEM2_{FL}$) is shown as reference. **g,** Bulk co-condensation assay of LEM2 and BAF showing enrichment of BAF in LEM2 condensates. **h,i,** Normalized averaged line intensity profiles of BAF, LEM2 and DNA across nuclei (left, red line) for different cells plotted as mean ± sd (right). **j,** Quantification of mean pixel intensity of DNA in the nuclear rim across knockdowns (N= 368, 220, 196 for siCtrl, siLME2 and siBAF respectively) indicating that the enrichment of DNA does not change with depletion of BAF or LEM2. Vertical lines in the violins show median and the interquartile ranges. **k,** Confocal images showing bulk phase separation upon concentration titration of LEM2. **l,** Assay of condensate volume fraction to determine $C_{sat}$ for LEM2 (n=33,34,47,43,32 for input protein concentrations 17.58µM, 20.51µM, 23.44µM, 26.37µM, 29.29µM respectively, methods). Bars and whiskers represent mean ± s.d respectively. Scale bar, 5 µm. The statistical test used is Kruskal-Wallis test with multiple comparisons using Dunn's method in **j**. *$p < 0.05$, **$p < 0.005$, ***$p < 0.001$, ****$p < 0.0001$, *ns: non-significant*.

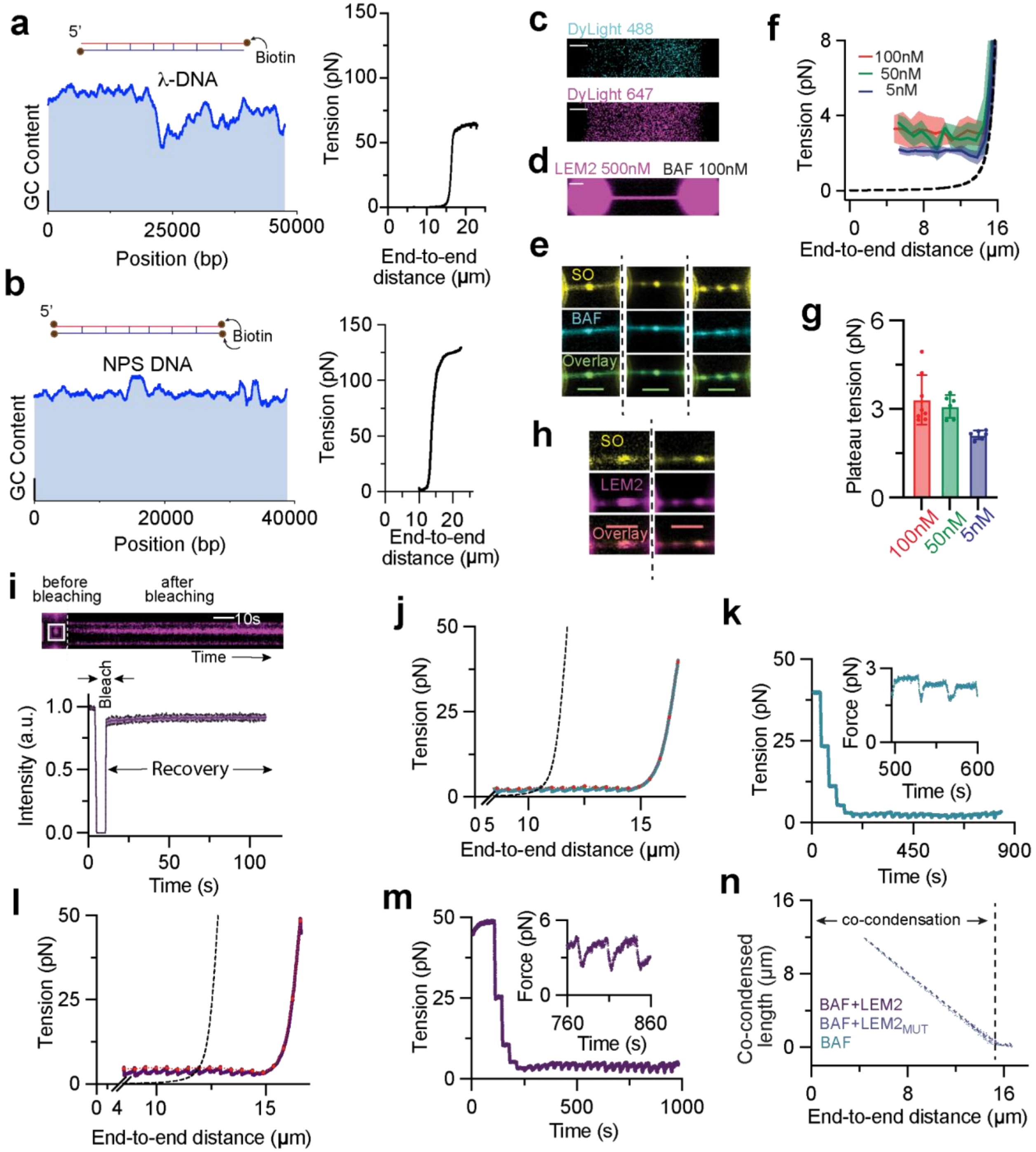


**Extended Data Fig. 2: Reconstitution of DNA-BAF-LEM2 co-condensate in optical tweezers.**

**a,b,** Schematic diagrams and GC content profiles (Scale bar, 10%) of biotinylated (**a**) λ-DNA and (**b**) NPS DNA (left), with corresponding tension-extension curves (right). **c,** Confocal images of control experiment to test binding of the free chemical dye to DNA (tethered between beads), **d,** Confocal image of binding of LEM2 (labeled)+BAF to NPS DNA. **e,** One or more co-

condensates of DNA and BAF in optical tweezer with 100nM BAF (fluorescently labelled). **f,** Average tension-distance profiles of DNA with different concentration of BAF while the end-to-end distance is decreased suggesting that BAF binding to DNA saturates around 100 nM. Black dashed line indicates the eWLC fit. Data is plotted as mean±sd (n=8 for 100nM, n=6 for 50nM, n=6 for 5nM). **g,** The corresponding quantification of plateau tension at 5 µm end-to-end distance from tension-distance profiles in f. **h**, Confocal images showing one or more co-condensates of DNA-BAF-LEM2 in optical tweezer, with 100nM BAF (unlabelled) and 500nM of LEM2 (fluorescently labelled). **i,** FRAP of BAF-LEM2 co-condensate in optical tweezer. Top, pre-bleach confocal slice and post-bleach kymograph of FRAP time course for DNA-BAF-LEM2 co-condensate (inside white ROI). Bottom, average recovery profile of normalized fluorescence intensity (n=7). **j,l,** Tension-distance profiles of DNA while the end-to-end distance is decreased in case of **(j)** BAF and **(l)** BAF-LEM2. Red lines indicate the steady-state values of force for each end-to-end distance. Black dashed lines represent eWLC fit to calculate the amount of co-condensed DNA (methods). **k,m,** Tension-time profiles of DNA while the end-to-end distance is decreased in case of (**k**) BAF and (**m**) BAF-LEM2. Inset shows the equilibration of force with time (methods). **n,** Quantification of total co-condensed DNA length as a function of end-to-end distances for BAF, BAF-LEM2 and BAF-LEM2$_{MUT}$ (BAF : n=8; BAF-LEM2: n = 7; BAF-LEM2$_{MUT}$: n = 7).

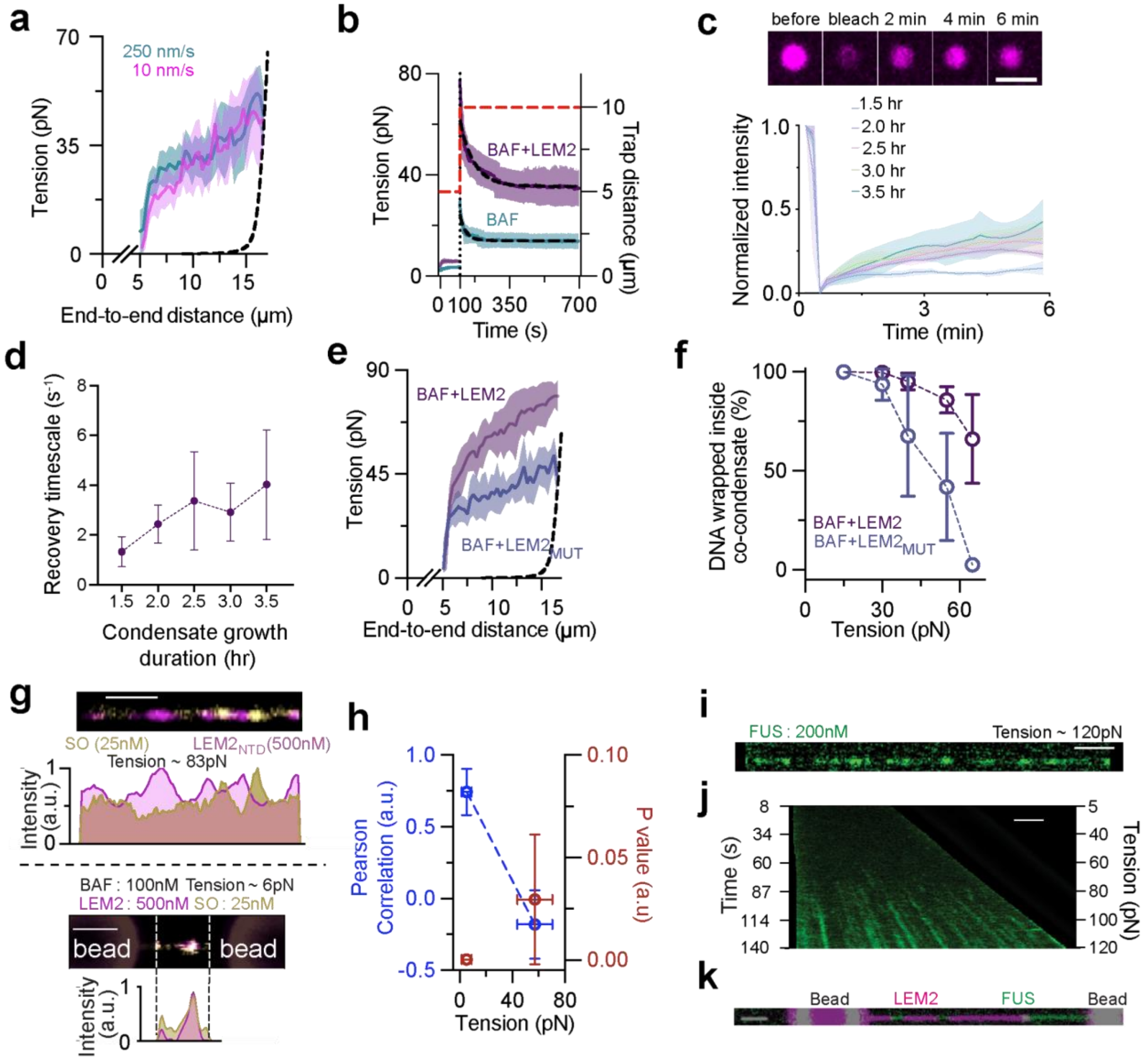


**Extended Data Fig. 3: DNA is protected inside DNA-BAF-LEM2 co-condensate.**

**a,** The tension-distance profiles when the BAF co-condensates are disassembled as the end-to-end distance is increased at different speeds. **b,** Temporal tension relaxation profile following an instantaneous increase in trap-to-trap distance from 5 μm to 10 μm (BAF-LEM2:n=11, BAF:n=9, methods). **c,** FRAP of BAF-LEM2 co-condensate in bulk. Top: Confocal images showing fluorescence recovery after photobleaching (FRAP) of BAF-LEM2 condensates at indicated time points. Bottom, average normalized fluorescence recovery curves at different time points post-condensate formation (1.5 to 3.5 hours). Quantification shows mean ± s.d. of n = 5 replicates except for 2.0 hr, where n = 4. **d,** The timescales of recovery of fluorescence intensity calculated from a simple exponential fit (methods) **e,** Average tension-distance profiles during co-condensate disassembly across multiple experiments for BAF-LEM2MUT (n = 5; BAF+LEM2 is replotted for reference). The dashed black line represents the eWLC fit for naked λ-phage DNA. **f,**

Quantification of DNA wrapped inside the co-condensates as a function of applied tension (BAF+LEM2 is replotted for reference). **g,** Top, confocal image showing the binding pattern of SO on DNA in the presence of BAF-LEM2 co-condensates at high tension, with normalized intensity profiles of SO (yellow) and LEM2 (magenta) along the DNA (methods). Bottom, co-localization of LEM2 and SO signal and corresponding normalised intensity profiles at low tension. Scale bar, 2 µm. **h,** Quantification of the Pearson correlation coefficient of LEM2 and SO signals at low tension (n=6) and high tension (n=8). **i,** Confocal slice showing the binding of FUS-GFP to ssDNA. **j,** Kymograph showing FUS-GFP signal emergence coinciding with melted ssDNA formation during DNA extension. **k,** Overlay of LEM2 (stains dsDNA) and FUS (stains ssDNA) signals on overstretched λ-DNA.

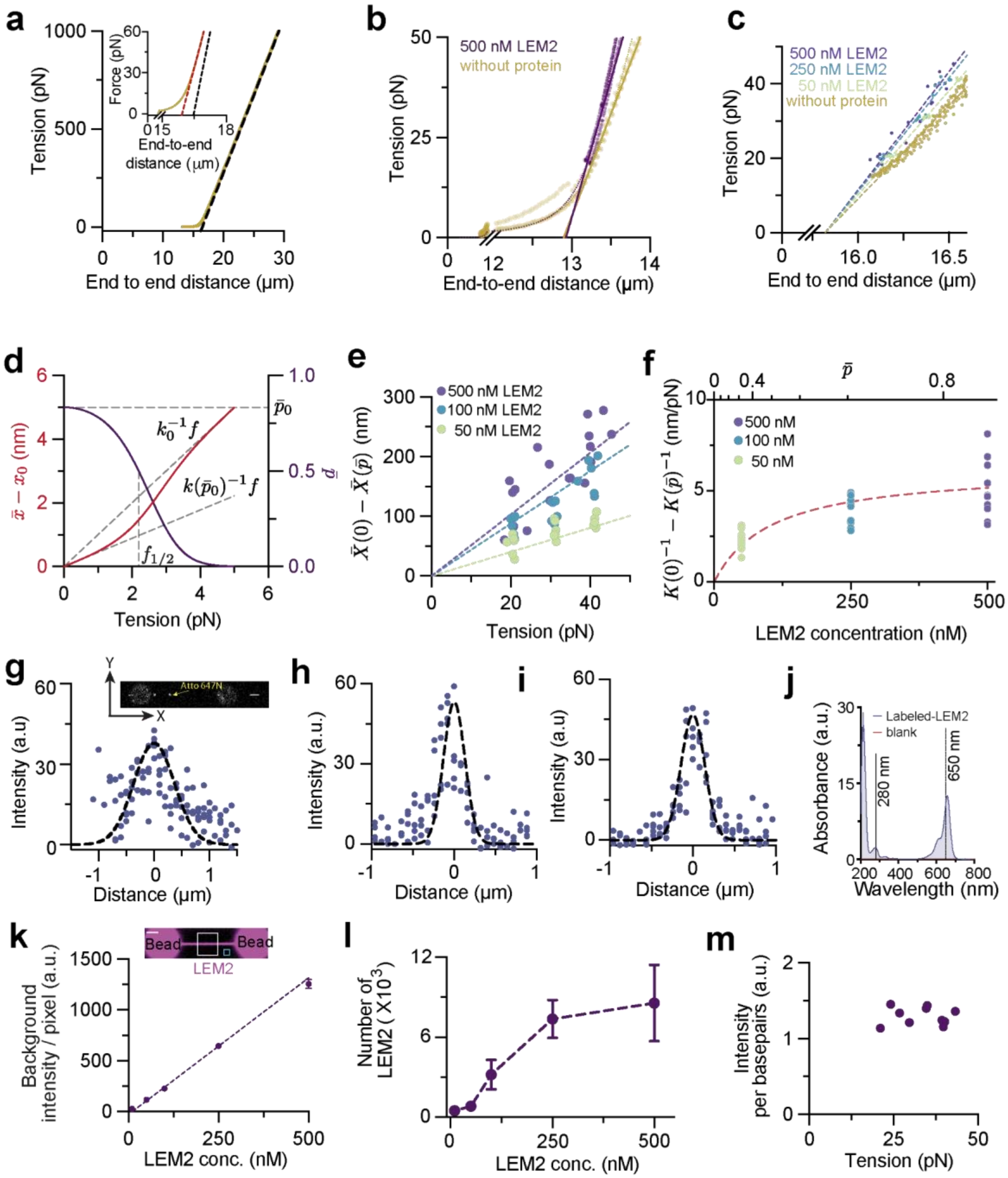


**Extended Data Fig. 4: DNA stiffness is modulated by the BAF-LEM2 module.**

**a,** Asymptote (black dashed line) to the eWLC model (yellow line, for λ-DNA), indicating that the rest length is shorter than the contour length for linear fits (red dashed line, inset) in the 20-40 pN range. **b,** Tension-distance profiles of NPS-DNA with (magenta) and without (yellow) BAF(100nM)-LEM2(500nM) while the end-to-end distance is increased. Dotted lines indicate the eWLC fits, whereas the dashed lines are linear fit to the respective data (methods) within 95% CI.

(naked DNA:n=3, BAF-LEM2:n=3). **c,** Steady state end-to-end distance vs. tension for λ-DNA under a constant-tension protocol, across different LEM2 concentrations (with 100 nM BAF). Linear fits (dashed lines) show progressively higher stiffness with increasing LEM2 concentration (500 nM: n=18; 250 nM: n=18; 50 nM: n=17). **d.** Ensemble average stretch $\bar{x} - x_0$ per binding site (solid line) and average occupation fraction $\bar{p}$ (dotted line) as a function of applied tension on DNA (parameters used : $k_0 = 1$ pN/nm; $k_i = 2$ pN/nm; $c_0 e^{\beta} \varepsilon = 100$ µM, c = 500µM, Supplementary Note 1). Two regimes (at low and high tension) are shown by grey dashed lines with the corresponding stiffnesses $k_0$ and $k(\bar{p}_0)$ indicated. The half-saturation tension $f_{1/2} = \sqrt{2(\mu - \epsilon) k_0 (1 + k_0/k_i)}$ is indicated as a vertical grey dashed line). **e**, Shortening of the end-to-end distance $\bar{X}(0) - \bar{X}(\bar{p})$ under constant-tension condition (from **c**) plotted against applied tension for varying LEM2 concentrations (with 100nM BAF). Dashed lines represent a global fit to the statistical mechanics model (Supplementary Note 1). **f,** Change in DNA compliance $K(0)^{-1} - K(\bar{p})^{-1}$(from panel **e**) shown as a function of LEM2 concentration (with 100nM BAF). The red dashed line represents a global fit to the statistical mechanics model (Supplementary Note 1). The top axis shows the corresponding occupation fraction. **g,** Quantification of the microscope's point spread function (PSF) from the intensity distributions of a single fluorophore. Individual points represent the maximum of pixel values along the x-y plane for each z-slice (methods). The solid line denotes the Gaussian fit. Inset, representative confocal image showing single Atto-647N dyes conjugated to λ-DNA. **(h-i)** Corresponding intensity distributions along (**h**) x and (**i**) y in the focal plane, used to determine the PSF in the x and y directions (n=5). **j,** The UV-Vis spectrum of labeled LEM2-DyLight 650 used to calculate the labeling efficiency (methods). **k,** Calibration of the mean background intensity as a function of input LEM2 concentration. Inset, representative confocal image showing LEM2 (tagged with DyLight-650) bound to DNA in optical tweezers. Cyan and white box indicate the ROIs used for background and DNA bound protein intensity quantifications (methods). **l,** Average number of LEM2 bound in presence of 100 nM BAF as a function of LEM2 concentration (**k,l** LEM2:500 nM n=11; LEM2:250 nM n=10; LEM2:100 nM n=7; LEM2:50 nM n=9; LEM2:10 nM n=12). **m,** Average pixel intensity, as a measure of total number of LEM2 bound to DNA across different forces for 100 nM BAF and 500 nM LEM2 (n=10).

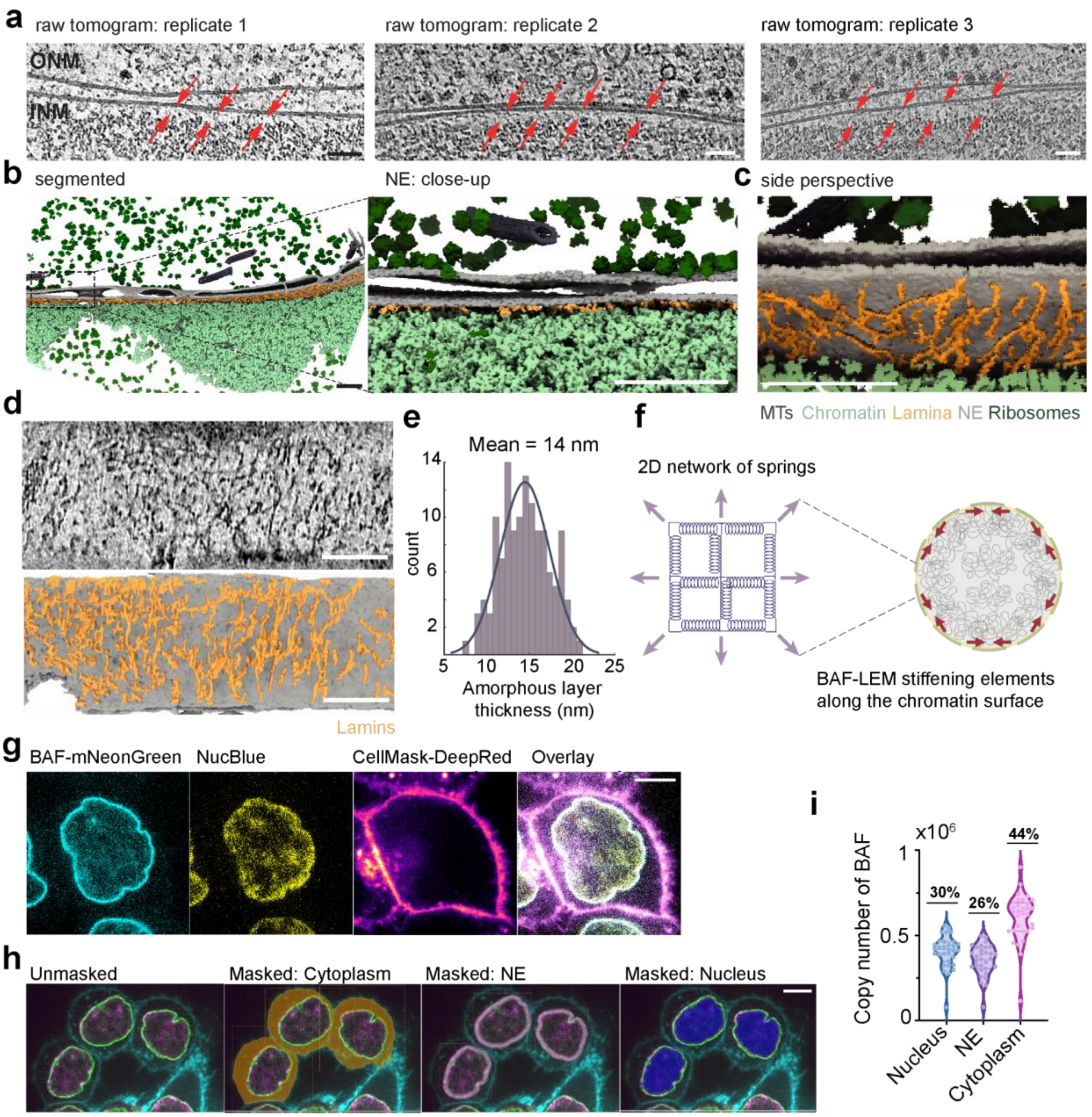


**Extended Data Fig. 5: Visualization of the nuclear lamina hydrogel and estimation of amounts of protein components.**

**a,** Cryo-electron tomography of the nuclear envelope; data obtained from [44]. The three independent tomographic slices reveal a continuous amorphous layer beneath the inner nuclear membrane (marked by red arrows). Scale bar, 50 nm. **b,** Left, 3D segmented model of a tomogram revealing the different molecular components at the nuclear periphery. Right, close-up snapshot of the nuclear envelope showing the exclusion zone flanked by INM and nucleosomes. Scale bar, 100 nm. **c,** Close up to the exclusion zone with nucleosomes models cropped to visualize the embedded lamin network. Scale bar, 100 nm. **d,** A bottom perspective of the nuclear surface showing attachment of the lamin network to the INM, where LEM2 is situated. Chromatin was cropped from the top to reveal the lamin network. Scale bar, 100 nm. **e,** Histogram showing the distribution of hydrogel thickness across tomograms, fitted with a normal distribution (mean thickness = 14 nm, n = 130). **f,** Schematic showing the translation of individual springs to a 2D meshwork (left) representing the network of BAF-LEM2 around the chromatin surface (right). **g,** Confocal images showing the distribution of BAF in different cellular compartments, nucleus, nuclear envelope and the cytoplasm. Scale bar, 5 µm. **h,** Representative images showing the 3D segmentation of BAF pools in the cytoplasm, nuclear envelope and the nucleus. Scale bar, 5 µm. **i**, The average copy numbers of BAF determined from the segmentation in the three cellular pools of BAF: nucleus, NE and cytoplasm (n = 31 cells, methods).

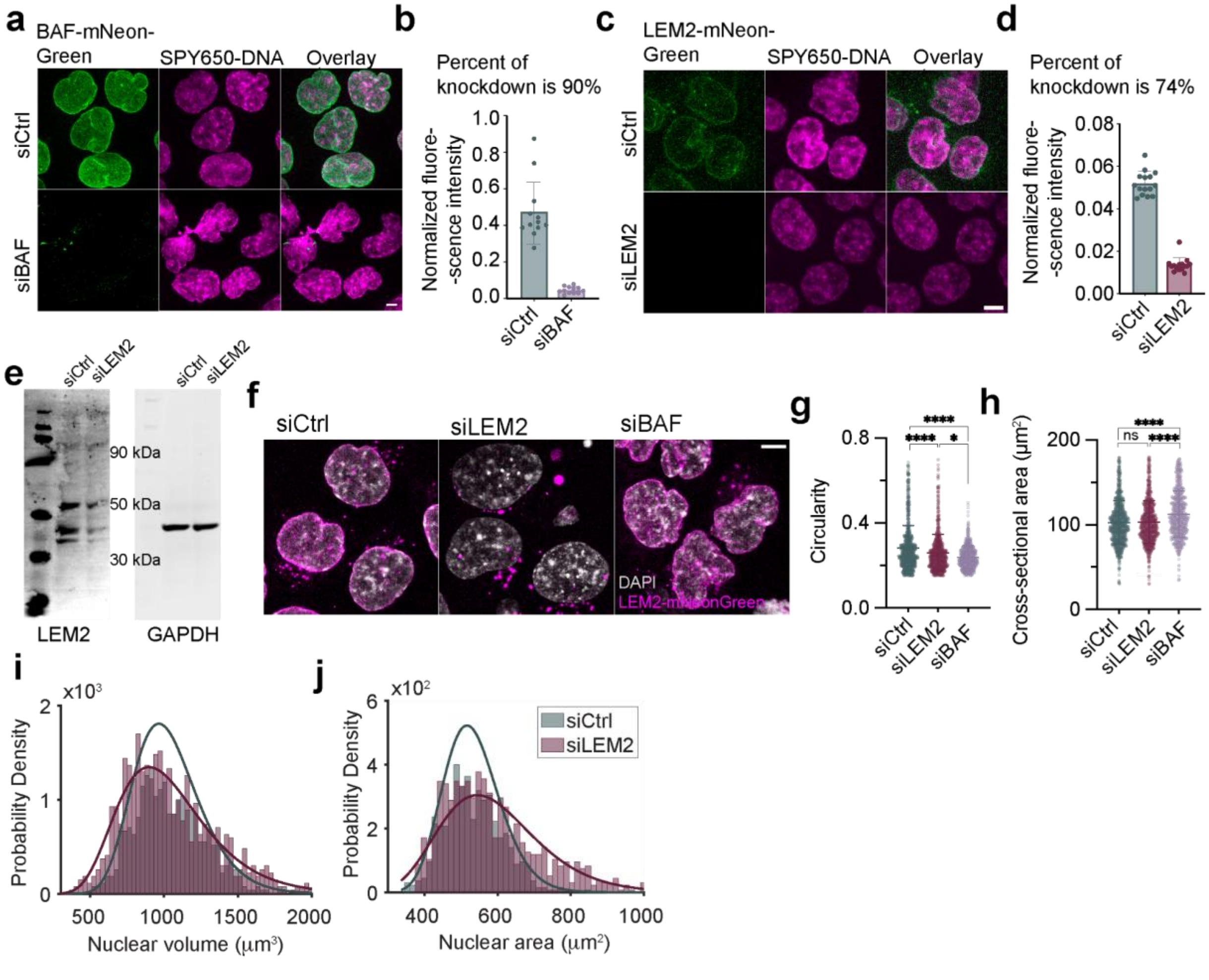


**Extended Data Fig. 6: Nuclear morphology breaks down upon LEM2 depletion.**

**a-d,** Quantification of protein knockdowns. Confocal images showing loss of fluorescence upon (**a**) BAF knockdown, (**c**) LEM2 knockdown and (**b,d**) corresponding quantification of the extent of knockdown from fluorescence loss (siCtrl vs siBAF; n = 12 each, siCtrl vs siLEM2; n = 15 each, each point represents independent fields of view). **e,** Western blot showing depletion of LEM2 in HCT116-Lamin A/C mNeonGreen cells. GAPDH is a loading control. **f,** 2D maximum projection confocal images of nuclei across knockdown showing morphology defects. **g-h,** Cross-sectional area (**g**) and circularity (**h**) of nuclei across knockdowns (siCtrl: n=973, siLEM2: n=790, siBAF: n= 720 cells, data pooled across three biological replicates). **i,** Histograms and corresponding lognormal fits of nuclear volume between siCtrl (965±19 $\mu m^3$) and siLEM2 (899±21 $\mu m^3$). **j,** Histograms and corresponding lognormal fits of the nuclear area between siCtrl (516±6 $\mu m^2$) and siLEM2 (544±8 $\mu m^2$). siCtrl: n = 603, siLEM2: n = 901 cells, pooled across three biological replicates in **i,j.** Bars and whiskers / midline and whiskers in scatter plots represent mean±s.d. respectively. Scale bar, 5 µm. The statistical test used is Kruskal-Wallis test with multiple comparisons using Dunn's method in **g,h**. **p<0.05, **p<0.005, ***p<0.001, ****p<0.0001*, ns: non-significant.

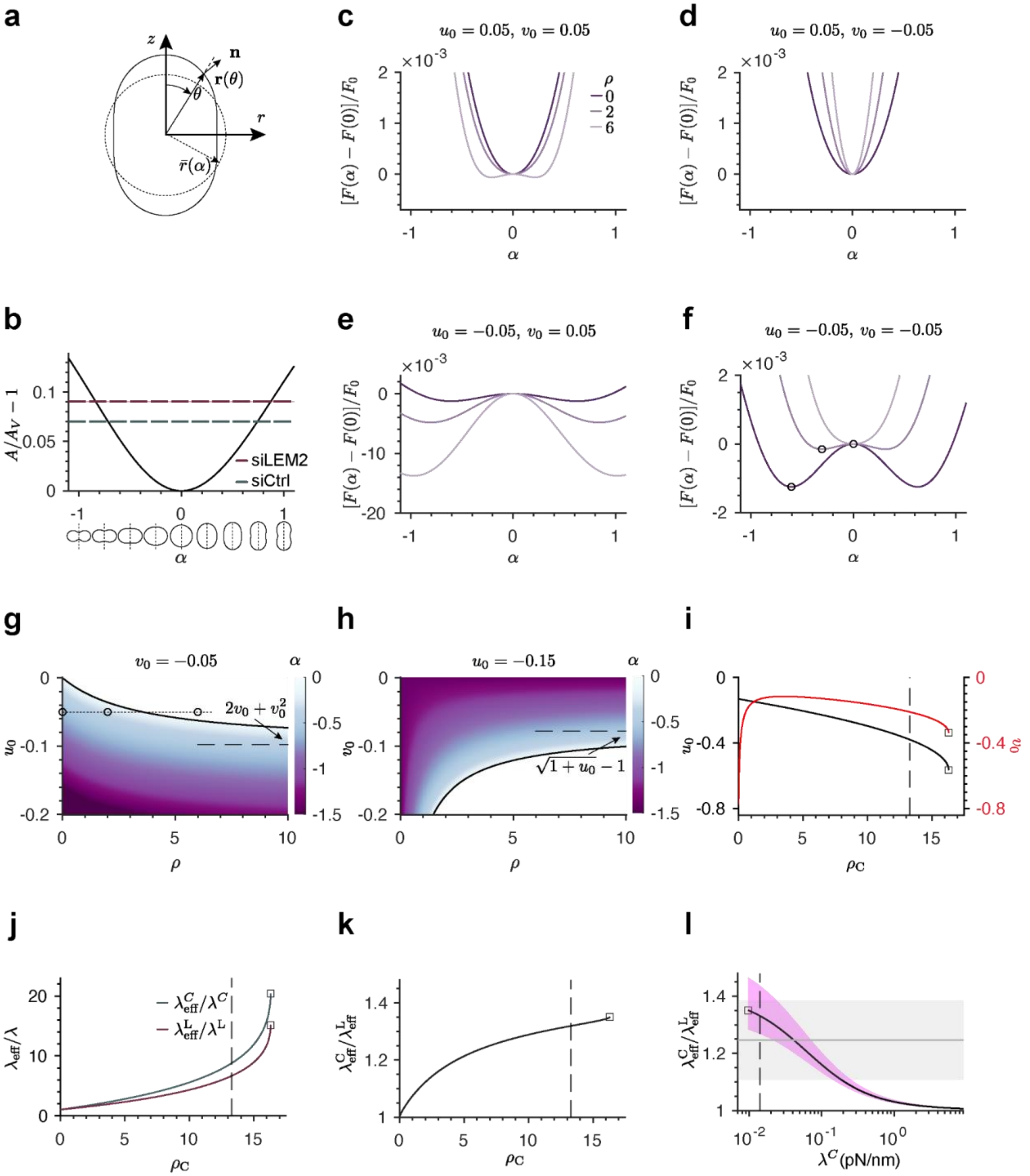


**Extended Data Fig. 7: Elastic surface hydrogel model.**

**a,** Axisymmetric shape parametrization $r(\theta)\ = \bar{r}\ (1 + \alpha\ Y_{20})$ (black solid line), where $\theta$ is the axial angle relative to the $z$-axis, $Y_{20}$ is the spherical harmonic with $l = 2$ and $m = 0$, $\alpha$ is the shape parameter, and the amplitude of $\bar{r}\ (\alpha)$ varies with $\alpha$ to ensure that the enclosed volume $V$ is fixed for all shapes. Here, **n** denotes the unit normal vector to the surface. Shape shown corresponds to $\alpha = 0.5$. **b,** Asphericity $(A/A_V - 1)$ shown as a function of $\alpha$, where $A$ is the surface area of the parametrized shape and $A_V = (6\sqrt{\pi}\, V)^{2/3}$ is the surface area of the sphere enclosing the same volume $V$. Shapes $r(\theta)$ shown for different values of $\alpha$ along the axis. Horizontal dashed lines indicate the asphericity values corresponding to the peaks of the distribution in Fig. 3f. **c–f,** Normalized free energy difference shown as a function of $\alpha$ for dimensionless BAF-LEM2 density $\rho$ = 0, 2, 6 (dark to light color). The parameter values $u_0 = A_V/A_* - 1$ and the rest length mismatch $v_0$ are indicated, where $A_*$ is the strain-free area. Circles in panel **f** indicate the energy minima corresponding to oblate shapes ($\rho = 0{,}2$) and to the sphere ($\rho = 6$). Here, $F_0 = \lambda A_*$, where $\lambda$ is the bare elastic area modulus. **g,h,** Phase diagrams in the $u_0$-$\rho$ plane (left) and $v_0$-$\rho$ (right) plane, where the color indicates the value of $\alpha < 0$, that minimises the free energy. The black critical line (Supplementary Note 1, section II) marks the second-order transition between spheres and oblate shapes. Open circles connected by a dotted line correspond to the open circles in panel **f**. Horizontal dashed lines in **g,h** indicate the asymptotic limit of the critical line for large $\rho$. **i-k,** Parametric plots of parameters $u_0$ and $v_0$ (**i**), normalized effective stiffness $\lambda_{eff}\ /\lambda$ (**j**), and ratio ${\lambda_{eff}}^C/{\lambda_{eff}}^L$ **(k)** corresponding to the two experimental conditions siCtrl (C) and siLEM2 (L). Plots are shown as a function of $\rho_C = \rho$ with $\rho_L = (\lambda^C/\lambda^L)\ {\xi_L}^{-1}\rho$, $\xi_L$ = 1.34 (95% C.I. 1.27–1.46). For these values, minimum energy shapes have the asphericities indicated in panel **b. l,** Ratio of effective area moduli ${\lambda_{eff}}^C/{\lambda_{eff}}^L$ shown as a function of the bare area elastic modulus $\lambda^C$. The measured value (Fig. 3d) is shown as a grey line. Shaded bands mark the 95% CI. In **i-l,** vertical dashed lines mark $\rho_C =$ 13.3 (Supplementary Note 1, Section IIG), open squares denote points where the solution ends.

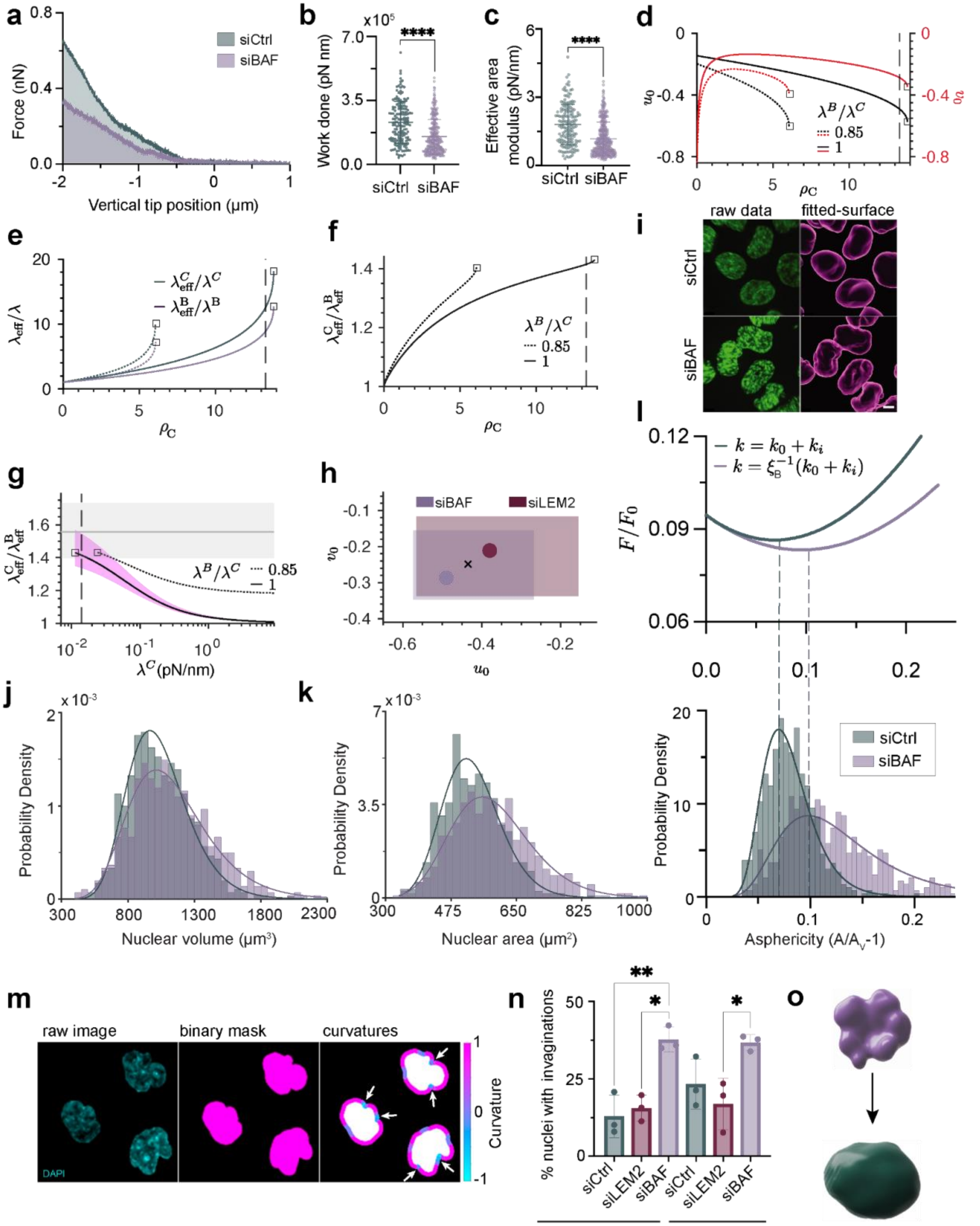


**Extended Data Fig. 8: The surface hydrogel model further accounts for shape and area elastic**

**modulus of BAF depleted nuclei. a,** Representative AFM force-indentation curves of nuclei of latrunculin A-treated HCT-116 cells, for siCtrl and siBAF cells. The cantilever makes contact with the nucleus at a vertical tip position of 0 µm, and we measure the mechanical work (shaded areas under the curves) required to indent the nucleus by 2 µm. Indentation in absence of BAF requires less work than in presence. Segmentation of the nuclear surface reveals the nuclear surface area increase (Extended Data Fig. 10b), which together with the mechanical work (**b**) allows for determination of the effective area modulus shown in (**c**, methods). **b,** Mechanical work for a 2 µm indentation of the nucleus of siCtrl and siBAF cells (methods), from three biological replicates (siCtrl: n = 159, siBAF: n = 319). **c,** Effective area modulus of nuclei of siCtrl and siBAF cells (Extended Data Fig. 10c; siCtrl: n = 159, siBAF: n = 319, methods). **d-f,** Parametric plots of parameters $\boldsymbol{u_0}$ and $\boldsymbol{v_0}$ **(d),** normalized effective stiffness $\boldsymbol{\lambda_{eff}\ /\lambda}$ **(e),** and ratio $\boldsymbol{{\lambda_{eff}}^C/{\lambda_{eff}}^B}$ **(f)** corresponding to the two experimental conditions siCtrl (C) and siBAF (B)**.** Plots are shown as a function of $\boldsymbol{\rho_C = \rho}$ with $\boldsymbol{\rho_B = (\lambda^C/\lambda^B)\ {\xi_B}^{-1}\rho}$, $\boldsymbol{\xi_B = 1.41}$ (95% CI 1.33–1.56), and $\boldsymbol{\lambda^B/\lambda^C}$ as indicated. For these values, minimum energy shapes have the asphericities indicated in panel **b. g,** Ratio of effective area moduli $\boldsymbol{{\lambda_{eff}}^C/{\lambda_{eff}}^B}$ shown as a function of the bare area elastic modulus $\boldsymbol{\lambda^C}$ for two values of $\boldsymbol{\lambda^B/\lambda^C}$, as indicated. The measured value from **c** is shown as a grey line. In **d-g,** vertical dashed lines mark $\boldsymbol{\rho_C = 13.3}$ (Supplementary Note 1, Section IIG), open squares denote points where the solution ends. **h,** Uncertainty in the $\boldsymbol{u_0}$ and $\boldsymbol{v_0}$ parameters is indicated by shaded regions defined by the intersection of the model curve CI band with the experimental uncertainty region (Extended Data Fig. 7l and **(g)**). Parameters corresponding to $\boldsymbol{\rho_C = 13.3}$ are indicated by filled circles. Average of these points ($\boldsymbol{u_0 = -0.44}$, $\boldsymbol{v_0 = -0.25}$) is indicated by black cross. This point is used for Fig. 3f and **l. i,** Top, confocal image stacks of DNA (SPYDNA650) of siCtrl (left) and siLEM2 (right) cells. Bottom, 3D segmentations of the nuclear surfaces for determining nuclear volume, shape, and surface area. **j,** Histograms and corresponding lognormal fits of nuclear volume between siCtrl (965±19 $\mu m^3$) and siBAF (1012±26 $\mu m^3$). **k,** Histograms and corresponding lognormal fits of the nuclear area between siCtrl (516±6 $\mu m^2$) and siBAF (556±9 $\mu m^2$). siCtrl: n = 603, siBAF: n = 591 cells in **j,k. l,** Top, plots of the normalized free energy $(\boldsymbol{F/F_0})$ of the REMM model (methods) as a function of nuclear asphericity reveal that the minimum for the siBAF condition with $\boldsymbol{k = {\xi_B}^{-1}(k_0 + k_i)}$ with measured stiffness ratio $\boldsymbol{\xi_B}$ (see Supplementary Note 1) is placed at a larger asphericity than for the siCtrl condition with $\boldsymbol{k = k_0 + k_i}$. Bottom, histograms of nuclear asphericity measured as in (**i**) for siCtrl and siBAF cells (n = 603 and n = 591, respectively, pooled across three biological replicates). Colored lines, log-normal fit. With REMM model parameters $\boldsymbol{\lambda = 0.014}$ pN/nm, $\boldsymbol{A_* = 1.8\,A_V}$, $\boldsymbol{\kappa = 0}$, rest length mismatch $\boldsymbol{x_0/x_* = 0.74}$ and energy scale $\boldsymbol{F_0 = \lambda A_*}$ . The histograms peak at the values that minimize the corresponding REMM surface hydrogel free energy $\boldsymbol{F}$ (dashed lines). **m**, Representative confocal image of nuclei, corresponding binary mask and calculated contour curvatures used to quantify the invaginations into nucleus. White arrows indicate curvature patterns counted as invaginations (methods). **n**, Percentage of nuclei having invaginations of the nuclear envelope quantified across knockdowns across three biological replicates (uncompressed siCtrl: n=6,10,10; siLEM2: n=9,16,13; siBAF : n =10,14,10, compressed siCtrl: n=17,11,10; siLEM2: n=19,11,11; siBAF: n=18,11,11). **o,** Schematic representing morphology shift from a less-prestrained state (siBAF) to more-prestrained state (siCtrl) for the nucleus upon BAF-LEM2 module activation. Scale bar, 5 µm. The statistical test used is Mann-Whiteney test with no assumptions on standard deviations between the samples in **b,c**. Kruskal-Wallis test with Dunn's correction for multiple comparisons in **n**. *$p<0.05$, **$p<0.005$, ***$p<0.001$, ****$p<0.0001$, ns: non-significant (not shown in **n**).

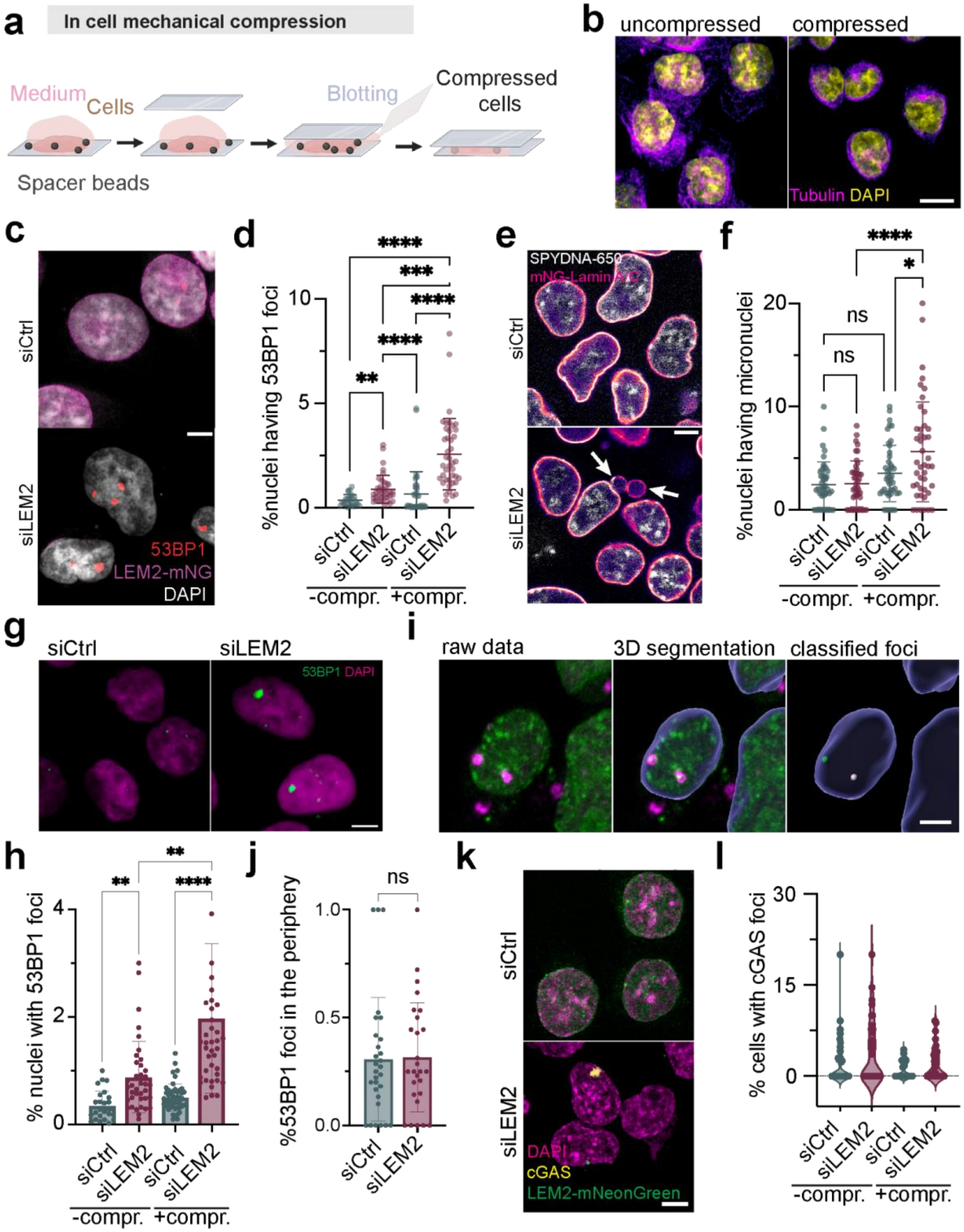


**Extended Data Fig. 9: Characterization of DNA damage and nuclear envelope rupture upon LEM2 depletion.**

**a**, Schematic of the live cell compression protocol. **b,** Maximum projection confocal images showing the loss of spindle architecture upon mechanical stress confirming compression of the nucleus. **c,** Representative confocal images showing DNA damage loci visualized by immunofluorescence of 53BP1 in interphase HCT116 LEM2-mNeonGreen cells following siRNA treatments as indicated and compression to 6 µm. The compression induced mechanical strain results in DNA damage foci in siLEM2 cells. **d,** Percentage of nuclei that form 53BP1 DNA damage foci following siRNA treatments as indicated and remain uncompressed or are compressed to 6 µm (see **c**; three biological replicates, uncompressed: n = 26, 38; compressed: n = 39, 41 fields of view for siCtrl and siLEM2 respectively). DNA damage increases ~2.2 fold in siLEM2 cells with compression compared to without compression. Non-significant comparisons are not shown. **e,** Representative confocal images of live HCT116 LaminA/C-mNeonGreen cells with micronuclei (white arrows) following siRNA treatments and compression to 6 µm. The compression induced mechanical strain results in micronucleation in siLEM2 cells. **f,** Percentage of nuclei that form micronuclei following siRNA treatments as indicated and compression to 6 µm (see **e**, three biological replicates, siCtrl: n = 48; siLEM2: n = 54 fields of view). **g,** Confocal images showing the accumulation of DNA damage foci upon compression of cells to a height of 3 µm. **h,** Quantification of percent of nuclei with DNA damage foci before and after compression from **g** (uncompressed: n=26, 38 and compressed n=48, 42 for siCtrl and siLEM2 respectively. Data is pooled from 3 biological replicates.). **i,** Segmentation of DNA damage foci and classification of foci to bulk (nuclear center up to 1 µm into the NE) or peripheral (1 µm into the nucleus) DNA damage. **j,** Quantification of percentage of DNA damage occurs at the periphery of the nucleus compared to the total damage foci observed (siCtrl: n=29, siLEM2: n=27 each point represents an independent field of view). Note that the distribution of DNA damage across the volume of nucleus does not change significantly between siCtrl and siLEM2. **k,** Maximum projection confocal images of nuclei with nuclear envelope rupture (NER) marked by cGAS foci, across knockdowns with and without compression. **l,** Quantification of percentage of cells with NER across conditions from **k** (uncompressed: n=36,35 and compressed: n=32,32 for siCtrl and siLEM2. Each point represents an independent field of view). Note that all comparisons are non-significant.

Bars and whiskers / midline and whiskers in scatter/violin plots represent mean ± s.d respectively. Scale bar, 5 µm. The statistical test used is Kruskal-Wallis test with multiple comparisons using Dunn's method in **d,f,h** and **l**. Mann-Whiteney test with no assumptions on standard deviations for **j**. * $p < 0.05$, ** $p < 0.005$, *** $p < 0.001$, **** $p < 0.0001$, ns: non-significant. Non-significant comparisons are not shown in **d**,**h** and **l.**

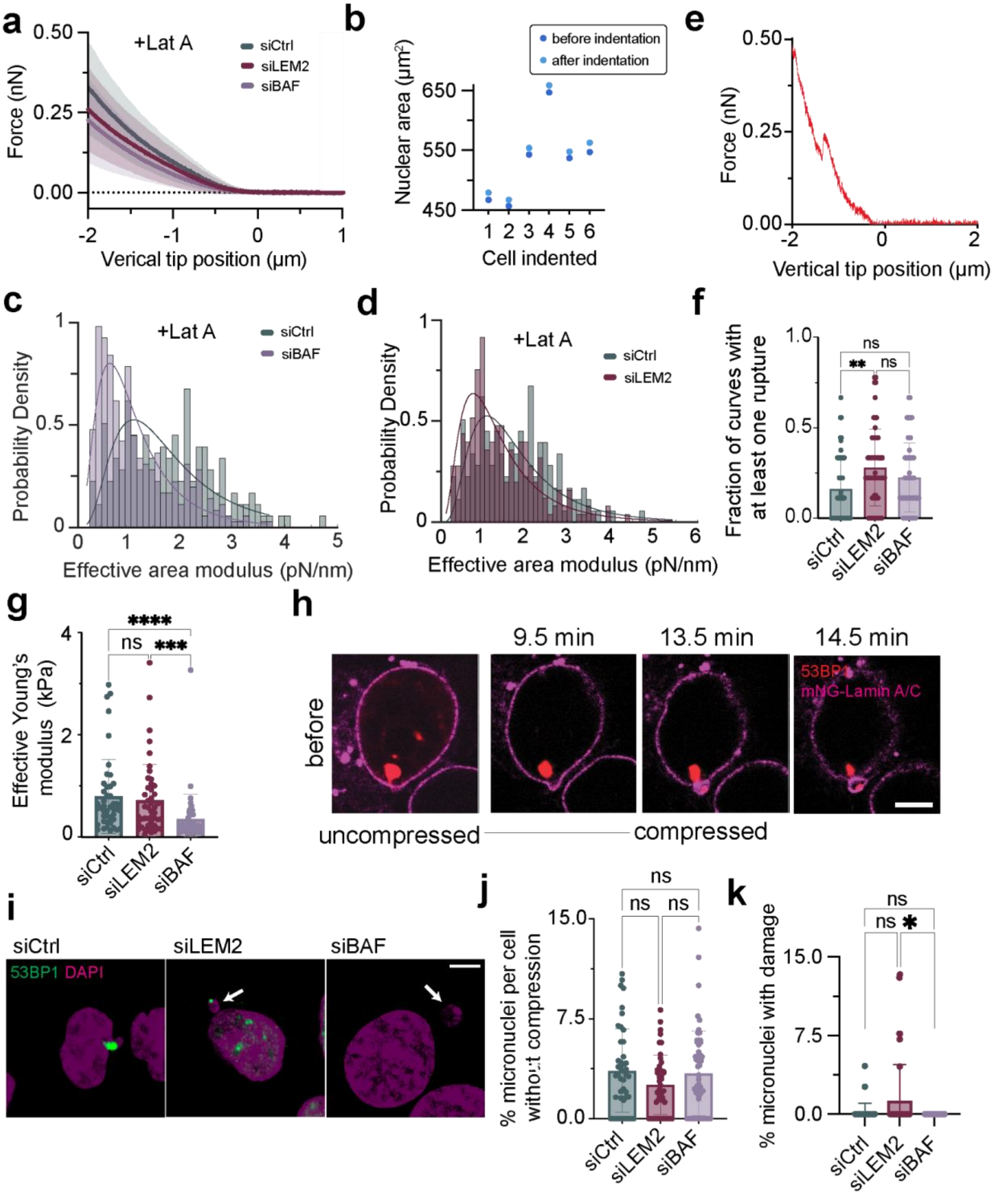
a
+Lat A
siCtrl
siLEM2
siBAF
Force (nN)
0.50
0.25
0.00
-2
-1
0
1
Verical tip position (μm)
b
before indentation
after indentation
Nuclear area (μm²)
650
550
450
1 2 3 4 5 6
Cell indented
e
Force (nN)
0.50
0.25
0.00
-2
0
2
Vertical tip position (μm)
c
+Lat A
siCtrl
siBAF
Probability Density
1
0.5
0
0 1 2 3 4 5
Effective area modulus (pN/nm)
d
+Lat A
siCtrl
siLEM2
Probability Density
1
0.5
0
0 1 2 3 4 5 6
Effective area modulus (pN/nm)
f
Fraction of curves with at least one rupture
1.0
0.5
0.0
ns
**
ns
siCtrl
siLEM2
siBAF
g
Effective Young's modulus (kPa)
4
2
0
****
ns
***
siCtrl
siLEM2
siBAF
h
9.5 min
13.5 min
14.5 min
before
53BP1
mNG-Lamin A/C
uncompressed
compressed
i
siCtrl
siLEM2
siBAF
53BP1 DAPI
j
% micronuclei per cell without compression
15.0
7.5
0.0
ns
ns
ns
siCtrl
siLEM2
siBAF
k
% micronuclei with damage
15.0
7.5
0.0
ns
ns
*
siCtrl
siLEM2
siBAF

**Extended Data Fig. 10: Mechanical weakening of the nuclei upon BAF or LEM2 module depletion.**

**a,** Average force-indentation traces of nuclei across knockdowns in the presence of latrunculin A (siCtrl: n = 159, siLEM2: n = 225, siBAF: n = 319, data pooled across three biological replicates). **b,** The area change observed upon intending the nuclei with an AFM cantilever (until 2 µm) for 6 replicates. **c,** Histograms and corresponding lognormal fits of effective area modulus calculated from individual AFM curves in the presence of latrunculin A for siCtrl and siBAF (siCtrl: n = 159, siBAF: n = 319, data pooled across three biological replicates, methods). **d,** Histograms and corresponding lognormal fits of effective area modulus calculated from individual AFM curves in the presence of latrunculin A for siCtrl and siLEM2 (siCtrl: n = 159, siLEM2: n = 225, data pooled across three biological replicates). **e,** Representative force-indentation trace showing a force dent corresponding to a NE rupture event. **f,** Quantification of the NE rupture events across knockdowns (n = siCtrl:55, siLEM2:57, siBAF:55 cells, methods). **g,** Young's modulus values for nuclei calculated from **a** (n = siCtrl:45, siLEM2:46, siBAF:45 cells, methods). **h**, Confocal time course of DNA damage occurring live upon mechanical compression resulting in the expulsion of a micronucleus in a control cell. **i**, Quantification of percentage of cells with micronuclei before compression (siCtrl: n=44, siLEM2: n=44, siBAF: n=55, data pooled across three biological replicates with each point representing an independent field of view). **j**, Confocal images slices showing formation of micronuclei with DNA damage foci contained inside (arrows point to damage foci) after compression. **k**, Quantification of percentage of nuclei having a micronuclei containing DNA damage (siCtrl: n=37, siLEM2: n=41, siBAF: n=35, data pooled across three biological replicates with each point representing an independent field of view).Bars and whiskers represent mean ± s.d respectively. Scale bar, 5 µm. The statistical test used is Kruskal-Wallis test with multiple comparisons using Dunn's method in **j,k,f,g**. * $p < 0.05$, ** $p < 0.005$, *** $p < 0.001$, **** $p < 0.0001$, ns: non-significant.

**Materials & Methods**

No statistical methods were used to predetermine sample size. The experiments were not randomized and, except where stated, investigators were not blinded to allocation during experiments and outcome assessment.

#### Section 1: Protein biochemistry and structure

##### A. Structure of LEM2 and NE protein distribution with AlphaFold

The LEM2 structure (Fig. 1a) was modeled using the published BAF-LEM assembly [52,64,65] and AlphaFold [66] predictions for remaining LEM2 domains (UniProt ID: Q8NC56) using ChimeraX [67].

To calculate the total protein mass at the nuclear envelope and their domain distribution, the repertoire of protein members of the inner nuclear membrane were first listed. Using OpenCell [68], UniProt and AlphaFold, the copy numbers and domain assignments of the proteins were curated. In brief, for each protein, the corresponding AlphaFold structure was examined and structured and disordered domains were annotated manually. The data was further analysed to categorize residues belonging to structured or disordered domains of BAF-LEM proteins, Lamins and the rest (Extended Data Table 1). The tabulated results were then plotted as a pie chart in Extended Data Fig. 1b.

##### B. Purification of LEM2 proteins

###### 1. LEM2

LEM2 was purified according to the published protocol[7] adapted to in-house equipment. All steps were carried out at room temperature unless otherwise noted. The plasmid encoding LEM2 (a gift from the Frost lab) was inserted into BL21 T7 Express pRare bacterial cells (produced in house) and expanded into 3 L of culture in TB with Kanamycin. Harvested cells were resuspended in lysis buffer (30 mM HEPES pH 7.4, 500 mM KCl, 5% glycerol, 20 mM imidazole, pH 8.0, Benzonase (250 U/mL stock,1:10000 dilution in final volume), 1 mM MgCl2, PMSF (100 mM stock,1:100 dilution in final volume) and cOmplete EDTA-free protease inhibitor cocktail (Roche, #38249996990, 1 tablet/50 mL), and lysed by three rounds of high-pressure homogenization. Lysate was then clarified by centrifugation (30,000g, 30 min), loaded on a pre-equilibrated PureCube 100 Compact Cartridge Ni-INDIGO column (Cube Biotech, #75302, 5 mL column volume, 5 mL/min loading rate, room temperature), and washed extensively with lysis buffer. Protein was eluted with lysis buffer supplemented with 300 mM imidazole, pH 8.0. The imidazole eluate was diluted using ice-cold shock buffer (40 mM HEPES pH 7.4, 5% glycerol) to bring down the salt concentration to 50 mM to induce the phase separation of $His_6$-SUMO-

LEM2. After incubation in ice for 20 minutes, the droplets were pelleted by centrifugation (10,000g, 10 min, 4 ºC). The pellet was resuspended in a 5 mL high-salt buffer (40 mM HEPES pH 7.4, 500 mM KCl, 5% glycerol) to dissolve the droplets. To this mixture, His6-Ulp1 (3.2 mg/mL stock, 1:500 equivalents) was added (4 ºC, overnight) to induce the cleavage of His6-SUMO tag. Spin-concentrated cleaved protein was further purified by size exclusion chromatography using the HiLoad Superdex 200 pg 16/600 column in high salt buffer. The LEM2-containing elution fractions were pooled, concentrated, and snap frozen in liquid $N_2$ as single-use aliquots.

###### 2. LEM2$_{MUT}$

LEM2$_{MUT}$ (Extended Data Fig. 1f) was purified following the LEM2 protocol, with the following modifications: the lysis buffer contained 350 mM KCl instead of 500 mM, the storage buffer had 110 mM KCl instead of 350 mM, and the preparatory phase separation step was omitted.

##### C. Purification of BAF

Full-length human BAF (Uniprot ID O75531) was purified following published protocols [9,65] with modifications to in-house equipment. $His_6$-SUMO-BAF was cleaved using $His_6$-Ulp1 protease (30 min, room temperature) to generate traceless BAF. The protein was then purified via a Superdex 200 pg 16/600 gel filtration column, and fractions containing the BAF dimer were pooled, concentrated, flash-frozen in liquid nitrogen, and stored as single-use aliquots at -80 °C.

##### D. Protein labelling for fluorescence imaging

LEM2 was fluorescently labeled using DyLight 650 NHS Ester (Thermo Fisher, #62265). For imaging, 1% labeled protein was mixed with the unlabeled fraction. BAF was labeled with DyLight 488 NHS Ester (Thermo Fisher, #46402), with a 5% labeled fraction supplemented for imaging experiments. The labeling efficiency was calculated from the UV-Vis spectrum of the labeled protein according to the manufacturer's instructions (Extended Data Fig. 4j).

#### Section 2: Preparation of DNAs used in optical tweezer

##### A. λ-DNA

Commercially available biotinylated bacteriophage λ-DNA was bought from Lumicks (#SKU: 00001) which has 6 biotins on the 3' end of one strand and 4 biotins on the 3' end of the other strand. In case of protein counting calibration measurement, described later, biotinylated λ-DNA with 2x ATTO 647N was used, also bought from Lumicks (#SKU: 00020).

**B. NPS DNA**

Two repeats of a 172 bp sequence, each containing a 147 bp nucleosome positioning site (NPS) (ACAGGATGTATATATCTGACACGTGCCTGGAGACTAGGGAGTAATCCCCTTG GCGGTTAAAACGCGGGGGACAGCGCGTACGTGCGTTTAAGCGGTGCTAGAGC TGTCTACGACCAATTGAGCGGCCTCGGCACCGGGATTCTCCAG and a 25 bp spacer (GGCGGCCACGAGATCCAATACATGC) was cloned into the pET-28a vector and amplified to 12 repeats using a previously described method [69]. The 12 NPS-spacer sequences, along with an additional 25 bp spacer and 57 bp flanking sequences, were excised from pET-28a and inserted into the 18,397 bp plasmid TH1704 via *XhoI/NcoI* restriction sites, generating the 19,887 bp plasmid TH1704-12NPS. This plasmid was linearized alongside the 23,286 bp fosmid TH1703 using *BamHI* and *HindIII*, followed by ligation to form the 39,782 bp fosmid TH1703/04-12NPS, with the 12 NPS repeats centrally positioned within a *NotI*-linearized plasmid. For optical tweezers experiments, *NotI* overhangs were ligated to biotinylated handles following [69,70] . A 1,870 bp DNA segment from pET-28a, containing a *PspOMI* site near the center (1 bp offset), was PCR-amplified (primers: CGAGATCTCGATCCTCTACG, GCTAACCAGTAAGGCAACC) in the presence of 10% Biotin-16-dUTP. Digestion with *PspOMI* produced *NotI*-compatible overhangs, generating randomly biotinylated 932 bp and 934 bp handles, which were ligated to the ends of the *NotI*-linearized 39,782 bp fosmid TH1703/04-12NPS. GC content of the DNAs were calculated as a moving average of counts of G and C with a window size 1000 base pairs in Extended Data Fig. 2a-b.

**Section 3: Bulk Assays**

**A. Fluorescence microscopy of phase separation and co-condensation**

Fluorescently labelled LEM2, BAF and DNA oligos were assayed for phase separation (Extended Data Fig. 1k-l) and co-condensation (Fig. 1b, with FUS in Fig. 1m, Extended Data Fig. 1g) by spinning disc confocal microscopy, performed in glass-bottom ULA coated low-binding 384-well plates (Revvity #6057302). Reaction was assembled directly in the 384 well plates with a total system volume of 20 µL. The phase separation of LEM2 was triggered by diluting the protein in a working buffer (30 mM Hepes pH 7.4, 150 mM KCl) to bring down the salt concentration from 500 mM KCl in the storage buffer to 150 mM KCl.  A concentration titration was performed to screen for phase separation across concentrations (Extended Data Fig. 1l). The formed LEM2 droplets were allowed to grow and settle for 30 min before the start of confocal imaging. The droplets were imaged with an Andor Eclipse Ti inverted spinning-disc microscope with an Andor iXon 897 electronmultiplying charge-coupled device camera and Yokogawa CSU-X1 disc (10000 rpm). The droplets were imaged using either a UPLSAPO ×100/1.45 numerical aperture (NA) oil-immersion objective or ×60/1.20 NA water-immersion objective (Nikon).

For co-condensation assays, different components were assembled together. LEM2 was diluted to 5 μM final concentration and 150 mM salt concentration to allow droplet formation alongside BAF and DNA-oligos. After 30 min of incubation at room temperature, the co-condensates were imaged to examine enrichment or exclusion of individual components. For multichannel imaging, $LEM_{NTD}$ was labeled with DyLight 650, BAF was labeled with DyLight 488 and DNA-oligo was labeled with Atto 565. FUS was tagged with eGFP. Excitation wavelengths were: 488 nm for DyLight 488; 561 nm for Atto565; 640 nm for DyLight 650.

##### B. Concentration titration for $C_{sat}$

For concentration titration (Extended Data Fig. 1k-l) of LEM2 in order to measure $C_{sat}$ , LEM2 dilutions of different concentrations were assembled in a water-in-oil emulsion, imaged and analysed as described in the in-house developed inPhase method for identifying phase diagrams [34].

##### C. FRAP of condensates to measure recovery timescales

LEM2 droplets with 1% labeled fraction were assembled in glass-bottom ULA-coated 384-well plates (PerkinElmer) and allowed to grow for 0.5 hr, 1 hr, 2 hr, 2.5 hr, and 3 hr. FRAP and droplet dynamics were imaged (Extended Data Fig. 3c) using an Andor Eclipse Ti inverted spinning-disc microscope with an Andor iXon 897 EMCCD camera, Yokogawa CSU-X1 scan head (10,000 rpm), and FRAPPA unit. A UPLSAPO ×100/1.45 NA oil-immersion objective (Nikon) was used, with 650 nm excitation for DyLight 650. Photobleaching was done with a 638-nm laser at 70% power for 20 ms per bleaching event. Images were captured pre-bleach (2 frames, 10 ms interval) and post-bleach (91 frames, 10 sec interval). The bleached spot and background were identified manually, and mean intensities were recorded using FIJI. Further data analysis was carried out using a custom MATLAB script to account for natural photobleaching, data normalization to pre-bleach condensate intensity and to determine the recovery timescales from a simple exponential fit for each condensate of different age. Replicates were aligned to the time of bleaching, averaged, and plotted mean ± s.d in Extended Data Fig. 3c,d.

#### Section 4: Optical tweezers

##### A. Protein Binding Kinetics on stretched DNA:

Optical tweezer experiments were performed using a C-Trap system (LUMICKS) with a microfluidic flow cell containing separate laminar flow channels. Two streptavidin-coated polystyrene beads (~4.34 μm, Spherotech #SVP-40-5) were trapped and moved into a channel with biotinylated DNA (λ-phage or custom made NPS DNA; 10 ng/ml; Extended

Data Fig. 2a,b) in a working buffer (20 mM HEPES, 150 mM KCl, pH 7.5). Single DNA tethering was confirmed by monitoring inter-bead distance and tension during modulation of the "steerable trap," ensuring the resulting tension-distance curve matched the eWLC model. Following tether confirmation, either a constant trap position or constant tension protocol was used based on experimental requirements.

1. **Constant trap position protocol:**

   In this case, beads tethered with DNA were first moved into a working buffer channel, where optical trap positions were fixed to apply a tension of ~40 pN. Without altering trap positions, the bead-DNA assembly was transferred to a channel containing various protein combinations in the same buffer. Depending on the proteins, incubation lasted several tens of seconds to allow binding (Fig. 1d). Real-time binding was monitored using integrated confocal imaging. As a control, the labeling dye alone showed no DNA association (Extended Data Fig. 2c). Tension was continuously recorded to assess whether protein binding altered DNA mechanics. Trap positions remained fixed throughout and maintained to a position that resulted in ~40 pN tension on naked DNA. Binding was observed for ~100 s (BAF-LEM2) or ~40 s (BAF, BAF-LEM2$_{MUT}$). Tension increase ($\Delta$T, Fig. 2b) was calculated as the difference between the mean tension during the final ~5 s in the protein and buffer channels.

   To assess the impact of tension and protein concentrations on binding kinetics (Fig. 2c), the same protocol was repeated with trap separations adjusted to apply ~30 pN or ~20 pN at different LEM2 concentration (with BAF:100nM).

2. **Constant tension protocol :**

   To investigate how protein binding affects DNA stiffness, tethered DNA was first transferred to a working buffer channel, where a constant tension was applied using a PID-enabled force-feedback system integrated with an optical tweezer setup. Maintaining this constant tension, the bead-DNA assembly was then moved to a channel containing various protein combinations and the binding was monitored via confocal imaging. The binding of the BAF-LEM2 module to DNA increased its stiffness, as indicated by a reduction in end-to-end distance (Extended Data Fig. 4c,e).

#### B. Observation of DNA-BAF-LEM2 co-condensates:

After proteins bound to stretched DNA, co-condensates formed as slack was introduced by stepwise reduction of the DNA end-to-end distance to ~5 µm. At each step, the steerable trap was moved ~800 nm toward the fixed trap, allowing the system to reach steady state while recording tension and end-to-end distance (Extended Data Fig. 2j for BAF; Extended Data Fig. 2l for BAF-LEM2). Tension required finite time to equilibrate (Extended Data

Fig. 2k, m), with equilibration time depending on protein combination and concentration. Steady state was typically reached within ~50 s, during which trap positions remained fixed. One mean steady-state tension-distance curve (red curve in Extended Data Fig. 2j,l) was generated by averaging tension and end-to-end distance over the final 5 s of each step. The average steady-state tension as a function of end-to-end distance for each protein condition (Fig. 1e, Extended Data Fig. 2f) was obtained by averaging multiple steady-state traces, binned by end-to-end distance (bin size ~650 nm). Notably, quantification of average tension in the co-condensate regime across different BAF concentrations shows that DNA binding by BAF saturates at around 100 nM (Extended Data Fig. 2f,g).

##### C. Co-condensed energy calculation :

The co-condensation energy per DNA base pair was calculated by first determining the total length of condensed DNA. At each end-to-end distance ($d_i$) and corresponding tension ($T_i$), the contour length of "non-condensed" DNA ($L_{nc,i}$) was computed using the eWLC model (black dashed line, Extended Data Fig. 2j, 2l):

$$L_{nc,i} = d_i/(1 - \frac{1}{2}\sqrt{\frac{k_B T}{T_i P}} + \frac{T_i}{S})$$

where $P$ is the persistence length (~50 nm) and $S$ is the stretch modulus (~1300 pN for BAF; ~2000 pN for BAF-LEM2). The "co-condensed length" was then $L_{c,i} = L_0 - L_{nc,i}$, with $L_0$ ~16.5 µm for λ-DNA (Extended Data Fig. 2n). The co-condensation energy per base pair was obtained by calculating the area between the tension-distance curves of naked and protein-bound DNA, normalized by ($L_{c,i}$/0.34), using 1bp=0.34nm.

##### D. FRAP of co-condensates in optical tweezer:

Fluorescence recovery after photobleaching (FRAP) was performed by first acquiring a ~5 s time series of confocal images of the co-condensate using low laser excitation (10%). A region of interest (ROI) that included one or more condensates was then photobleached using high laser excitation (90%) for 1s. Recovery was monitored by capturing a long time series (~100 s) at low laser excitation (10%) to track fluorescence recovery (Extended Data Fig. 2i).

##### E. DNA-co-condensate disassembly experiment :

After co-condensates were formed due to slack, the DNA was gradually stretched by moving one trapped bead with the steerable trap while recording tension. Confocal imaging was used throughout to capture condensate dynamics under applied tension. Pulling was performed at two speeds (10 nm/s and 250 nm/s), with no significant difference in dissembling behavior (Extended Data Fig. 3a). The average tension-distance curve (Fig. 1i) was generated by binning data by end-to-end distance (~250 nm bins) and calculating the mean tension for each bin.

To assess the temporal stability of co-condensates (Extended Data Fig. 3b), the trap-to-trap distance was abruptly increased from 5 µm to 10 µm and held for ~10 min. This induced a rapid tension spike (~80 pN for BAF-LEM2; ~40 pN for BAF), followed by a gradual decline to a stable value above the co-condensate plateau. This slow decrease suggests a delayed disassembly of DNA from the condensates, indicative of a long relaxation timescale and a potential hardening effect.

##### F. Co-condensates disassembly work calculation :

The amount of DNA within co-condensates ($L_{c,i}$) during unraveling (Fig. 1j) was determined using previously described eWLC-based calculations from the tension ($F_i$) and corresponding end-to-end distance ($d_i$) in the mean tension-distance traces (Fig. 1i). For BAF-LEM2, $F_i$ was set to 65 pN to determine $d_i$. This $d_i$ was then used to find the corresponding tension in the BAF condition and hence corresponding co-condensed length. The total co-condensed DNA length ($L_T$) at 5 µm end-to-end distance was set as the reference point (100% wrapped). The amount of DNA unraveled at each step was computed as $L_T-(L_0-L_{c,i})$. The disassembly work per base pair was calculated by integrating the area between the tension-distance curves of protein-bound and naked DNA, then normalizing by the amount of DNA base pairs unraveled till 65pN.

##### G. DNA stretching experiment without co-condensates :

Proteins were first allowed to bind to DNA held in an extended state for ~100 s under tension ≥20 pN using constant tension mode, as previously described - a condition that prevented co-condensate formation. Following binding, the tethered DNA was continuously stretched at 250 nm/s by moving the steerable trap, and the extension continued until the tether ruptured (Fig. 2d, Extended Data Fig. 4b).

##### H. Calculation of DNA rest length :

Rest length was estimated from an asymptotic fit to the eWLC model, ($T = S(d-L_0)/L_0$), which fits well at high tensions (T~1000 pN) and yields a rest length equal to the contour length (~16.5 µm for λ-DNA, Extended Data Fig. 4a). Due to experimental limitations, a linear fit between 25-50 pN to the eWLC model (P = 50 nm, S = 1300 pN) was used as an approximation, yielding a lower rest length via the x-intercept calculation. This was consistent with rest length estimates obtained from (i) linear fitting of final end-to-end distances in constant tension mode, yielding ~15.81 ±0.08 µm (Extended Data Fig. 4c), and (ii) linear fitting of the average traces from DNA stretching experiments without co-condensates, yielding ~15.84±0.02 μm (Fig. 2d). DNA stretching experiments with NPS DNA (Extended Data Fig. 4b) showed the same result: the rest length was ~12.90±0.07 μm for naked DNA and ~12.93±0.04 μm with the BAF-LEM2 module.

##### I. Calculation of DNA stiffness :

DNA stiffness at a given BAF-LEM2 concentration was determined using two complementary methods: (i) from the slope of a linear fit to DNA stretching traces in the absence of co-condensates (Fig. 2d, protocol described above), and (ii) from the slope of a linear fit to final end-to-end distances measured at varying tensions using the constant tension method (Extended Data Fig. 4c, protocol described above). Fig. 2e shows the effective DNA stiffness obtained via the second method, plotted as a function of BAF-LEM2 concentration and fitted using a three-parameter model (Supplementary Note 1, Eq. S18). The resulting fit yielded the following parameter estimates with confidence intervals : $k_0/N$ = 50.21 pN [49.05, 51.37], $k_i/k_0$ = 0.30 [0.26, 0.35], $\exp(\beta\epsilon)c_0/c_L$ = 66.87 [32.69, 101.06].

A control experiment confirmed that stiffness change or tension rise was not due to varying LEM2 binding to DNA at different tensions, as average bound fluorophore intensity (tagged to LEM2) remained constant across tensions (Extended Data Fig. 4m).

##### J. Analysis of LEM2/SO intensity distribution along DNA length:

To correct for background protein (LEM2 or SO) signal, a small ROI excluding DNA and beads was selected from the confocal image (cyan box, Extended Data Fig. 4k). The mean intensity ($b^k$) due to protein of choice (k = LEM2 or SO) of this ROI was subtracted from each pixel intensity, $I_{i,j}^k$, of the image, where i and j denote row and column indices of the pixel considered. The protein intensity along DNA was calculated as a function of column index j :

$$I_j^k = \sum_{i=1}^{N_1} (I_{i,j}^k - b^k)$$

with $N_1$ representing the number of rows in the considered ROI of the image. Finally the protein intensity was normalised by the maximum intensity as $I_{norm,j}^k = I_j^k \,/\, max\,(I_j^k)$ (Fig. 1l, Extended Data Fig. 3g). Pearson correlation coefficient between the LEM2 and SO signal was calculated as

$$Pearson\ correlation\ coefficient\ = \frac{\sum_{j=1}^{N_2} (I_{norm,j}^{LEM2} - \langle I_{norm,j}^{LEM2}\rangle)(I_{norm,j}^{SO} - \langle I_{norm,j}^{SO}\rangle)}{\sqrt{\sum_{j=1}^{N_2} (I_{norm,j}^{LEM2} - \langle I_{norm,j}^{LEM2}\rangle)^2}\sqrt{\sum_{j=1}^{N_2} (I_{norm,j}^{SO} - \langle I_{norm,j}^{SO}\rangle)^2}}$$

where $N_2$ represents the number of columns in ROI considered in the image (Extended Data Fig. 3h).

**K. DNA stretching experiment with co-condensates in presence SO :**

Co-condensates were initially formed by incubating pre-slacked DNA with BAF (100nM) and LEM2 (500nM), along with SO (25nM). Following the previously described protocol, the steerable optical trap was used to stretch the DNA-co-condensate assembly while simultaneously recording the applied tension and imaging the unraveling process. The intensity profiles of LEM2 and SO along the DNA were then quantified using the method described above (Extended Data Fig. 3g). We observe an anti-correlation between LEM2 co-condensates and the SO signal, indicating that DNA tension is high whenever the DNA is outside the co-condensate, and lower inside (Extended Data Fig. 3h). This result is not due to SO exclusion from co-condensates, which is supported by two control experiments revealing a positive correlation between LEM2 co-condensates and SO signal at low tensions that lie within the constant tension plateau (Extended Data Fig. 3g, bottom).

**L. DNA overstretching experiment with in presence FUS :**

To confirm that FUS binds exclusively to single-stranded DNA (ssDNA), two control experiments were conducted :

1. **NPS-DNA stretching in the presence of FUS:**

   An NPS-DNA construct stretched with 200 nM GFP-FUS (in working buffer, described previously) showed binding only above ~80 pN (Extended Data Fig. 3i,j), to the melted ssDNA. No binding occurred below 65 pN, consistent with previous reports where biotinylated, bead-tethered strands prevented peeling under high tension (Extended Data Fig. 2b).

2. **λ-DNA Overstretching and Exposure to BAF + LEM2 + FUS:**

   λ-DNA was overstretched and partially melted at ~65 pN due to the presence of nicks. Upon exposure to a mixture of BAF-LEM2 and FUS, complementary binding of FUS to ssDNA and BAF-LEM2 to dsDNA was observed (Extended Data Fig. 3k). No binding of FUS to dsDNA was detected under the working buffer conditions.

**M. Protein count on DNA :**

To avoid nonspecific signal from proteins bound to the beads, the number of proteins bound to DNA was quantified within a ROI (Extended Data Fig. 4k) carefully selected to exclude the beads. The total number of DNA-bound proteins was then extrapolated from this ROI.

To estimate the number of proteins bound to stretched DNA in the optical tweezer setup, the confocal volume, $V_C$, of the imaging system was determined by imaging λ-DNA labeled with ATTO-647N (Lumicks, #SKU: 00020) across different focal planes (z-stacks). Extended Data Fig. 4g shows the total photon count from an individual fluorophore as a function of z-position, with $z=0$ corresponding to the plane of maximum intensity. The data were fitted with a Gaussian to determine the full width at half maximum (FWHM), giving the confocal depth (~Δz). Photon count distributions along the x and y directions at z = 0 (Extended Data Fig. 4h,i) were fitted with Gaussians to extract the FWHM, representing the point spread function in x and y (Δx, Δy). The confocal volume was then calculated as $V_C=\Delta x \cdot \Delta y \cdot \Delta z$.

In point-scanning mode, the background intensity per pixel in DNA-free regions corresponds to fluorescence from a single confocal volume, representing $c \cdot V_C$ LEM2 molecules, where $c$ is the LEM2 concentration (Extended Data Fig. 4k). The average intensity per protein molecule was then estimated as $I_p=b^{LEM2}/(c \cdot V_C)$ where $b^{LEM2}$ is the average background photon count per pixel in bulk, calculated from a small ROI excluding DNA and beads. The total background-subtracted photon count in the ROI, which includes the DNA, was computed as $J = \sum_{j=1}^{N_2} \sum_{i=1}^{N_1} (I_{i,j}^{LEM2} - b^{LEM2})$ where $N_1$ and $N_2$ are the total number of rows and columns in the ROI. The number of bound LEM2 (Extended Data Fig. 4l) was then calculated as $J * (d/0.08)/(N_1 * I_p * f_{eff})$, where $d$ is the DNA end-to-end distance (in µm), 0.08 µm is the pixel size, and $f_{eff}$ (= 66%, Extended Data Fig. 4j, methods) is the fluorescence labeling efficiency, representing the fraction of labeled versus total LEM2 molecules.

#### N. Work done per LEM2 molecule :

To calculate the mechanical work performed per LEM2 molecule, we first estimated the total work associated with the shortening of the DNA end-to-end distance under constant tension, following the constant-tension experimental protocol. At a force of 40.1 ± 2.4 pN, we observed a shortening of 226 ± 43 nm (Extended Data Fig. 4e) in the presence of 100 nM BAF and 500 nM LEM2. Based on the estimated number of LEM2 modules bound to λ-DNA (8557 ± 2875, Extended Data Fig. 4l), this corresponds to an average mechanical work of 0.26 ± 0.10 $k_BT$ per BAF-LEM2 module.

#### Section 5: Cell biology, Imaging and Analysis

##### A. Cell lines and cell culture

All cell lines used in this study were routinely tested and confirmed negative for mycoplasma contamination. Their sources and authentication details are provided in the Supplementary Note 2. Cells were cultured in McCoy's 5A medium (Gibco, #16600082) with 5% (v/v) fetal bovine serum (FBS; Gibco, #A5670701) and, when overexpression constructs were used, supplemented with puromycin (0.5 µg/mL, Gibco, #A11138).

Cells were cultured in T25 or 60 mm plastic dishes or 35 mm glass-bottom dishes (Nunc, #169900, 150288 & 150680 respectively) or 8-well glass-bottom µ-slides (Ibidi, #80827). Cells were regularly inspected for growth and passage using a Nikon Eclipse TS2 cell culture microscope with 10X or 40X air objective. Cells were counted either manually using a Neubauer chamber or using Luna II brightfield cell counter. Cells were always maintained at 37 °C and 5% $CO_2$ while storing and imaging. LEM2, BAF, and Lamin A/C were visualized via stable expression of C-terminal mNeonGreen-tagged constructs. Live-cell imaging was performed in McCoy's 5A medium with 5% (v/v) FBS, omitting phenol red to minimize autofluorescence.

##### B. CRISPR/Cas9-mediated generation of HCT116 knock-in lines

Cell lines that stably express fluorescently labelled marker proteins, BAF-mNeonGreen, LEM2-mNeonGreen and Lamin A/C-mNeonGreen were produced by the Genome Engineering Facility at MPI of Molecular Cell Biology and Genetics, Dresden. Genome editing was performed using a CRISPR–Cas9 nickase strategy.

hBAF and hLEM2 were targeted for C-terminal tagging, hLamin A/C for N-terminal tagging with mNeonGreen in HCT116 cells (Public Health England #91091005). Sequences of target regions were analyzed for appropriate guide RNA binding sites using the Geneious (2022.1.1, http://www.geneious.com/) software and the CRISPOR design webtool (http://crispor.tefor.net/). The genomic target sequences in HCT116 cells were checked beforehand for a perfect match with sequences of guide RNAs and targeting constructs by Sanger sequencing. Guide RNAs (Supplementary Note 2) were ordered as crRNA from IDT (Integrated DNA Technologies). crRNAs were duplexed with tracrRNA (IDT #1072534) by mixing equimolar amounts. Targeting constructs containing codon-optimized mNeonGreen flanked by 500 bp homology arms were generated through Gene Synthesis by Genscript (sequences in Supplementary Note 2). HCT116 cells were transfected with RNP (44 pmol cr:tracr duplexes and 37 pmol Cas9, IDT #1081061) and 500 ng of targeting construct using the NEON transfection system (1530V, 20ms, 1 pulse). 72 h post-electroporation single-cell sorting was performed in a BD FACSAria Fusion flow cytometer (Beckton Dickinson). Genomic DNA from single-cell clones was extracted using the QuickExtract DNA extraction kit (Epicentre) following the manufacturer's instructions. Cell clones with tagged allele (BAF: monoallelic, Lamin A/C: biallelic and LEM2: biallelic) were identified by PCR using Phusion Flash High-Fidelity PCR Master

Mix (ThermoFisher) and verified by Sanger Sequencing (primer sequences in Supplementary Note 2).

**C. Plasmid and small interfering RNA transfections**

For both transient and stable expression of fluorescently labelled markers, we used pMGF196 vectors having siRNA resistant gene for protein or mutant of interest coupled with an antibiotic resistance gene for puromycin that allow the protein of interest and the resistance gene to be expressed from the same transcript. The plasmids encoding LEM2-mCherry and $LEM2_{M21}$-mCherry were a gift from the lab of Katie Ullman, University of Utah. The blotted plasmids were extracted into solution and midi-prepped for further use. The plasmid encoding mApple-53BP1trunc was purchased from Addgene (#69531).The plasmid encoding $LEM2_{MUT}$-mCherry was prepared by Gibson assembly of human codon-optimised synthesised $LEM2_{MUT}$ gene block and pMGF196 backbone coding for mCherry and puromycin.

For transient expression, plasmids were transfected using X-tremeGENE 9 DNA transfection reagent (Roche, #XTG9-RO) or Lipofectamine 2000 (Invitrogen, #11668019) following the manufacturer's instructions. Transfected DNA amounts were optimized for each assay and varied between 5 µg and 8 µg per 20 µl transfection reagent in 250 µl Opti-MEM (Gibco, #11058021). After adding the plasmid-lipid mix, the cells were incubated for 24 h before replenishing fresh medium (imaging medium for direct confocal imaging, or McCoy's 5A with 5% FBS supplemented with 5 µg/mL Puromycin for antibiotic selection, accordingly). For stable expression, plasmids were transfected using Lipofectamine 2000, and incubated for 24 h before antibiotic selection.

Small interfering RNAs (siRNAs) targeting BAF or LEM2 were obtained from previous studies[8,9]. siRNAs were delivered with lipofectamine RNAiMax (Invitrogen, #13778150) according to the manufacturer's instructions. BAF (also known as BANF1) was targeted using 10 nM s16808 (AACGGAUUGAAAGUGAGGGtt), IDT, including a 3′ overhanging tt dinucleotide for increased efficiency) and analysed after 48 h; LEM2 was targeted using 10 nM (UACAUAUGGAUAGCGCUCCtt, IDT, including a 3′ overhanging tt dinucleotide for increased efficiency) and analysed after 48 h. A scrambled siRNA sequence (UCUACGAUCGGAGAUUUGCtt, IDT, including a 3′ overhanging tt dinucleotide for increased efficiency) was used as a non-targeting siRNA control (siCtrl).

**D. Western blotting**

HCT116 Lamin A/C-mNeonGreen cells treated with siLEM2 or siCtrl were scratched from the culture dish and lysed in disruption buffer (NP40 Lysis buffer, #J60766.AP, 50mM Tris-HCl (pH 7.4), 150mM NaCl, 1% NP-40 and 5mM EDTA). The soluble protein fraction was separated by centrifugation (30 min, 30,000 xg), and the supernatant was collected. Total protein concentration was measured using a Bradford assay. Optimized amounts of lysate (10 or 25µg) from both control and knockdown samples were loaded on NuPage 4–12% gradient Bis–Tris gels (Invitrogen, #NP0321BOX) and transferred to a nitrocellulose membrane (Bio-Rad, #1620112) via wet blotting. LEM2 was detected with a polyclonal rabbit anti-LEM2 antibody (Sigma Aldrich, HPA017340, 1:500), and

GAPDH was used as a loading control, probed with a polyclonal mouse anti-GAPDH antibody (Invitrogen, MA5-15738, 1:5000). The membrane was imaged (Extended Data Fig. 6e) using the iBright Gel Documentation System (Thermo Fisher Scientific) and analyzed in FIJI.

#### E. Cell compression assays

Cells to be compressed were seeded on fibronectin (Gibco, #33016015) coated (2 h) coverslips (12 mm × 12 mm) at a density of 0.8 × $10^5$ cells/mL in 30 mm dishes and incubated at 37°C with 5% $CO_2$ for two days. On the third day, cells were washed three times with PBS (3 mL each) before aspirating excess liquid to ensure a dry surface. A 100 µL medium droplet containing spacer beads (3 µm or 6 µm, diluted 1:1000 from a 2.5% w/v stock, Polysciences Polybead® microspheres, Cat: 17134-15 (3 µm) and 07312-5 (6 µm)) was added to the coverslip to form a dome. After 5 min of bead settling, a clean coverslip was placed on top, gently touching one edge before release. The medium was blotted away using Whatman filter paper until the coverslips were nearly in contact, compressing the cells for 5 min. Fresh medium (3 mL) was then flushed into the dish to allow the top coverslip to float before removal. Cells were incubated for 30 min post-compression before proceeding with immunofluorescence staining. The compression of cells was verified by visualizing the disruption of microtubule organisation in immunostaining for tubulin (Synaptic Systems, #302 206G) in Extended Data Fig. 9b.

For cell compression with real-time imaging, the same protocol was followed, except that cells were seeded on 30 mm glass-bottom dishes, and circular coverslips were used to compress the cells from above.

#### F. Live-cell microscopy

Confocal microscopy to test stable expression of endogenously mNeonGreen tagged LEM2, BAF and Lamin A/C was performed on Andor Olympus IX81 inverted spinning disc confocal microscope equipped with Andor iXon 897 EMCCD camera, Yokogawa CSU-X1 (5000 rpm, pinhole diameter 50 µm, pinhole spacing 250 µm) and with temperature, $O_2$, and $CO_2$ control. The excitation wavelengths were 405 nm, 488 nm, 561 nm and 640 nm. The images were acquired using either an Olympus U Plan SApo 60X (1.35 NA) Oil-immersion or Olympus U Plan SApo 100X (1.4 NA) Oil-immersion objective. The microscope was controlled using Andor iQ 3.6 software with custom made protocols.

Cells with siRNA treatments (Extended Data Fig. 7a,c), transient or stable expression of fluorescently tagged proteins and BAF-mNeonGreen cells for measuring BAF distribution were imaged using an Andor Revolution spinning disk confocal system on an Olympus IX83 inverted microscope with a cage incubator for temperature, $CO_2$, and $O_2$ control. Excitation was provided by 405 nm, 445 nm, 488 nm, 514 nm, 561 nm, 633 nm, and 726 nm lasers from two Andor ILE units. A Yokogawa CSU-W1 (4000 rpm) spinning disk unit with 50 µm pinhole (500 µm spacing) was used. Images were acquired with an Andor iXon 888 Ultra EMCCD camera using either an Olympus UPlanSApo 100X (1.4 NA) oil-immersion or Olympus U ApoN 150X (1.45 NA) objective. The system was controlled with Andor iQ 3.6 software with custom made protocols.

Imaging for nuclear shape analysis across knockdowns and imaging of cells with live-cell compression were performed using an Olympus IXplore SpinSR spinning disk confocal system on an Olympus IX83 inverted motorized stand with hardware autofocus (ZDC2) and a stage-top Z-piezo (400 µm range). A cage incubator maintained temperature, $CO_2$, and $O_2$ levels. Excitation was provided by 405 nm, 445 nm, 488 nm, 515 nm, 561 nm, and 640 nm lasers. A Yokogawa CSU-W1 (4000 rpm) spinning disk unit with 50 µm and SORA disks enabled standard and super-resolution imaging. Images were acquired with two Hamamatsu ORCA-Fusion BT Digital CMOS cameras using Olympus UPlanSApo 10X (0.4 NA) air or Olympus U-ApoN 100X (1.49 NA) oil-immersion objectives. The system was controlled with Olympus cellSens 4.1 software with custom made protocols.

#### G. Immunofluorescence

Immunofluorescence (IF) was performed at room temperature unless stated otherwise. Cells were either compressed (as mentioned before) or left uncompressed and proceeded to wash. The medium was aspirated out and washed gently with PBS twice. The cells were then fixed in 2% paraformaldehyde (10 min, dark, Thermo Scientific Chemicals, 043368.9M, diluted using PBS) and permeabilized with 0.2% Triton-X 100 (5 min, dark, Thermo Scientific Chemicals, #J66624.AP). Blocking was done using PBS-Tween-BSA (30–40 min, dark, Thermo Scientific Chemicals, #003005). Primary antibodies were diluted in PBS-Tween-BSA (50 µL per coverslip) and incubated in a humidity chamber for 1 hour at room temperature or overnight at 4°C. Coverslips were washed twice with PBS before secondary antibody staining (1:1000 in PBS-Tween-BSA, 30 min–1 hour, dark). DAPI/Hoechst (1:10,000, Invitrogen, #62248 or H21492) was added in the final PBS wash (10 min, if required). Coverslips were mounted using Vectashield (5 µL per coverslip, Vectorlabs, #H-1000-10) and sealed with nail polish. Samples were imaged within one week to prevent signal decay.

Primary antibodies used for immunodetection included polyclonal rabbit anti-LEM2 (Sigma Aldrich, #HPA017340), chicken anti-tubulin (Synaptic Systems, #302 206G), rabbit anti-BANF1 (Abcam, #AB129184), mouse anti-53BP1 (Merck Millipore, #MAB3802) and mouse anti-cGAS (Cell Signalling, #15102T). Fluorescently labeled secondary antibodies (Goat anti-rabbit 488, anti-mouse 488, anti-rabbit 647, and anti-rabbit 568; Invitrogen) were used for detection. The stained samples were imaged by laser-scanning confocal microscopy using a Leica Stellaris 8 Falcon upright microscope with a white light laser that allows excitation of fluorochromes at any wavelength in a range of 440 - 790 nm and a 405 nm DMOD diode laser. The microscope is equipped with an 8 kHz tandem scanner. Detection was achieved using three Power HyD S, one Power HyD X, and one Power HyD R detectors, along with a Hamamatsu Flash 4.0 V3 camera. Images were acquired using an HC PL APO 63X (1.40 NA) oil-immersion objective or HC PL APO 10X (0.30 NA) air objective. The microscope was controlled using Leica LAS X software with custom made protocols.

#### H. DNA damage and nuclear envelope rupture analysis

Cells (control and knockdown conditions) were compressed (6 µm or 3 µm) as described before. Uncompressed cells were used as control for mechanical strain. DNA damage in interphase cells (with and without compression across knockdowns) was assessed using anti-mouse 647 against mouse anti-53BP1 and anti-chicken 568 against chicken anti-tubulin. Nuclear envelope rupture (with and without compression across knockdowns) was assessed using anti-mouse 647 against mouse anti-cGAS and anti-chicken 568 against chicken anti-tubulin. DNA was stained with DAPI in all the cases (1:10000). LEM2 was imaged using the endogenous mNeonGreen tag. The imaging datasets for DNA damage and cGAS were acquired separately. For each field of view, a complete z-stack covering the whole of nuclei was recorded for all the four channels imaged (405: DAPI, 488: LEM2-mNeonGreen, 561: Tubulin, 647:53BP1). Each condition was imaged in three biological replicates (Extended Data Fig. 9c,g,k).

DNA damage was assessed using a custom ImageJ macro. Maximum intensity z-projections of the DNA and 53BP1 channels were generated. Nuclei masks were created by applying an Otsu threshold to segment nuclei based on DNA intensity contrast. Nuclei touching the image boundaries or exhibiting mitotic or degraded morphology were excluded based on size, circularity, and coordinates. The masks were applied to the 53BP1 channel to restrict damage quantification to nuclear regions and reduce background interference. DNA damage foci, defined as 53BP1 clusters ranging from 5 to 20 pixels (pixel size = 0.065 µm), were counted in each nucleus. Given minimal intra-field variation, the number of foci per nucleus was calculated for each field (image positions were randomized). Data from all replicates were pooled and plotted as bar/scatter plots displaying mean ± s.d. with individual data points (Extended Data Fig. 9d,h).

Nuclear envelope rupture was evaluated using a similar approach, incorporating the measurement of the total intensity of cGAS at each focus after background subtraction. Data were pooled from three independent replicates for each knockdown and compression condition and analyzed to determine the percentage of cells with cGAS foci (100 × number of cGAS foci per nucleus) in each image. The results are plotted in Extended Data Fig. 9l as violin plots displaying the mean ± s.d. with individual data points shown.

#### I. DNA damage close to nuclear periphery

DNA damage foci near the nuclear periphery were quantified using a custom workflow in Imaris, based on the same immunofluorescence imaging dataset. Using the DNA channel in the 3D volume image, nuclei were reconstructed into 3D surfaces using the Surfaces function with automatic thresholding, and clumped nuclei were separated by applying a seed diameter of 8 µm. Incomplete nuclei or those touching image boundaries were excluded, and additional filtering removed abnormally large or small nuclei (e.g., mitotic or defective). Damage foci were segmented from the 53BP1 channel using the Spots tool with a typical diameter of 1 μm. Using the MATLAB XTension 'Spots close to surface', foci within 1 μm of the nuclear rim were classified as peripheral, while the rest were considered bulk foci (Extended Data Fig. 9i). Data from three biological replicates were pooled, and the fraction of peripheral foci was plotted for each knockdown condition as a bar plot with mean ± s.d. and scatter of individual data points (Extended Data Fig. 9j).

##### J. Nuclear invaginations analysis

Nuclear invaginations were analyzed using the same datasets acquired for DNA damage analysis. Image analysis was performed using a custom Python script. The DNA channel was first subjected to a maximum z-projection followed by smoothing using Gaussian blur ($\sigma = 1$) and thresholded using Otsu's method to generate a binary mask. The mask was refined by filling holes and applying morphological opening (disk size: 5-pixel diameter) to remove dark spots within bright objects. Nuclei were then segmented using a watershed algorithm, and those with an area outside 5000–20000 square pixels (pixel size = 100 nm)were excluded.

For each selected nucleus, a contour of the nuclear boundary was generated. Curvatures along the contour were measured using a moving window of 100 points and classified into three categories: -1 for curvature ≤ -0.01, 0 for -0.01 < curvature < 0.01, and 1 for curvature ≥ 0.01. To minimize noise, the curvature profile was smoothed using a 30-point window. Invaginations were identified as regions where the contour exhibited a transition from negative to positive curvature spanning at least 10 pixels (Extended Data Fig. 8m). The number of invaginations per nucleus was calculated for each nucleus in each field of view across replicates. Data were pooled for each knockdown and compression condition and analyzed to generate Extended Data Fig. 8n, where individual points represent replicates, and bars indicate mean ± s.d. of replicate means.

##### K. Micronucleation under live compression of cells

Cells (control and knockdown conditions) were grown in 30 mm dishes at a density of 8 x $10^5$ and proceeded to live cell compression as mentioned before. The cells were pre-incubated with SPYDNA650 (Spirochrome) to stain DNA. The dish containing cells were mounted on a spinning disc confocal microscope within a temperature, $CO_2$ and $O_2$ controlled cage as described before. The focus of the microscope was aligned to the midplane of the nuclei of the monolayer of cells by manual inspection. Only one z-plane was imaged given the fast time-scales of compression and subsequent events. 5 frames were recorded prior to compression at an interval of 20 sec and then cells were compressed *in situ*. Imaging was continued immediately after compression recording 91 frames at an interval of 20 sec. Excitation wavelengths were 488 nm (LaminA/C-mNeonGreen) and 640 nm (SPYDNA-650). Representative frames post-compression is shown in Extended Data Fig. 9e.

Micronuclei in the imaged z-plane were manually counted before and after compression with dataset blinding. Only nuclear fragments with diameters between 1 and 3 μm were classified as micronuclei. The number of new micronucleation events was determined by subtracting the pre-compression count from the post-compression count. Data from different replicates were pooled for each knockdown or compression condition and analyzed to generate Extended Data Fig. 9f and Extended Data Fig. 10j (comparing siBAF) where mean ± s.d. is displayed as a bar plot with individual data points.

To determine whether live micronuclei budding events are associated with DNA damage, HCT116 cells with endogenously tagged Lamin A/C-mNeonGreen were transfected with a construct expressing mApple-53BP1trunc. Live-cell compression and micronucleation tracking were performed as described, except that 53BP1-trunc was imaged instead of DNA. Due to the low transfection efficiency of the plasmid in these cells, the data were used qualitatively in Extended Data Fig. 10h, without quantitative analysis from live-cell imaging. Instead, fixed compressed samples subjected to immunofluorescence were used for quantification.

#### L. Counting micronuclei and damage foci inside micronuclei

Micronuclei and DNA damage foci within them were analyzed using a custom CellProfiler pipeline applied to the same immunofluorescence dataset used for DNA damage assessment. First, maximum intensity Z-projections of the DNA and 53BP1 channels were generated (Extended Data Fig. 10i). The DNA channel was then downsampled (0.5×, nearest neighbor method), followed by morphological opening (disk size = 35 px). The image was upscaled back (2×, nearest neighbor method) and subtracted from the original to have an image with higher contrast between background and foreground (Image #2). To refine segmentation, Image #2 underwent morphological closing (disk size = 13 px), Sobel edge enhancement, and subtraction from the original. Micronuclei were identified using Otsu thresholding (diameter: 10–50 px, form factor ≥ 0.4, area ≥ 50 $px^2$). The resulting mask was applied to the 53BP1 channel, and DNA damage foci were segmented via Otsu thresholding (diameter: 1–15 px, area ≥ 25 $px^2$). Micronuclei counts per nucleus and foci counts per micronucleus were extracted. Data were pooled for each knockdown condition (compressed) and analyzed to generate Extended Data Fig. 10k, where the percentage of nuclei with damaged micronuclei was plotted as mean ± s.d. with individual data points.

#### M. Imaging and analyzing BAF distribution

The relative abundance of BAF in the nucleus, NE, and cytoplasm was quantified in HCT116 cells endogenously expressing BAF-mNeonGreen. Cells were imaged to track BAF (excitation: 488 nm), DNA (NucBlue, Invitrogen #R37605, excitation: 405 nm), and the plasma membrane (CellMask DeepRed PM stain, excitation: 638 nm). Z-stacks spanning the entire cell volume were acquired for all three channels with a 250 nm step size (Extended Data Fig. 5g).

Image processing and 3D segmentation were performed in Arivis Pro using a custom pipeline. To reduce noise, Gaussian blur ($\sigma = 1$) was applied to the BAF and CellMask channels, while a median filter (3-pixel wide pixel size=80 nm) was used for the DNA channel. The nuclear envelope (NE) and cell membrane were detected using the membrane detection tool in Arivis, with thickness and maximum gap size parameters set to 3 and 4 pixels, respectively. Membranes were segmented based on detected contours and Otsu thresholding of BAF and CellMask intensities. A nuclear DNA volume mask was generated via Otsu thresholding of DNA intensity and eroded by 3 pixels to account for confocal resolution limits. The NE volume mask was obtained by subtracting the DNA volume from the total NE-enclosed volume, while the cytoplasmic volume mask was defined by subtracting the NE-enclosed volume from the total cell volume (Extended Data Fig. 5h).

Segmented volumes and corresponding BAF intensity data were exported from Arivis for further analysis. For each nucleus, summed BAF intensities within the cytoplasmic, NE, and nuclear volume masks were computed across five replicates. Using mean intensity ratios and previously reported total cellular BAF copy numbers from mass spectrometry (OpenCell[68]), the absolute number of BAF molecules per compartment was estimated (Extended Data Fig. 5i).

The mean spacing of BAF dimers on the chromatin surface was calculated using the number of BAF dimers in the NE (N) and the total nuclear surface area (A), following the relation:

$$Mean\ spacing\ (l)\ =\ \sqrt{\frac{A}{N}}$$

#### N. Nuclear envelope enrichment of BAF, LEM2 and DNA

A plasmid encoding LEM2-mCherry was transfected into HCT116 cells, in which BAF was endogenously tagged with mNeonGreen, as described before. After 48 h, the cells were transferred to an imaging medium and stained with SPYDNA-650 for 15 min to label DNA. Imaging was performed using a spinning disk confocal microscope with excitation wavelengths of 488 nm for BAF-mNeonGreen, 561 nm for LEM2-mCherry, and 638 nm for SPYDNA-650. Complete z-stacks spanning the entire nucleus were acquired for all three channels with a z-step size of 250 nm (Fig. 1c, Extended Data Fig. 1i). A custom Python script was used to analyze the images, generating conventional normalized, background-subtracted line intensity profiles across the nucleus for BAF, LEM2, and DNA. The results are presented as mean ± s.d of replicates in Extended Data Fig. 1i.

DNA enrichment at the nuclear rim across knockdowns was assessed using CellProfiler with a custom pipeline. A maximum intensity projection of the DNA channel was performed, followed by Gaussian blur ($\sigma = 1$) and segmentation via Otsu's method. The resulting mask was eroded by 5 pixels and subtracted from the original to generate a nuclear envelope (NE) mask. The mean DNA intensity within the NE mask was quantified per nucleus across knockdowns after background subtraction. The results are shown as violin plots in Extended Data Fig. 1j.

#### O. Estimating extent of knockdowns of LEM2 and BAF

HCT116 LEM2-mNeonGreen or BAF-mNeonGreen cells treated with control or LEM2 siRNA were transferred to imaging medium after 48 hours and stained with SPY650-DNA for 15 minutes to label DNA. Imaging was performed using a spinning disk confocal microscope, acquiring Z-stacks spanning the entire nucleus (excitation: 488 nm for mNeonGreen-tagged proteins, 638 nm for DNA, Extended Data Fig. 6a,c).

Knockdown efficiency was assessed using a custom Python script based on fluorescence signal loss. Background subtraction was applied to each image, followed by mean intensity projections of the DNA and protein channels. Total protein and DNA intensities were then measured per image. To account for cell number, protein intensity was normalized by dividing it by total DNA intensity. Replicates were pooled, and fluorescence signal loss was quantified as the ratio of mean normalized protein intensity in control versus knockdown conditions. This fraction was converted to a percentage and plotted as bars (mean ± s.d.) with individual data points representing each image (Extended Data Fig. 6b,d).

#### P. Nuclear surface and shape calculations

For surface reconstruction, siRNA-treated HCT116 LEM2-mNeonGreen cells were used. Following 48 hours of siRNA treatment, cells were transferred to imaging medium, stained with SPYDNA-650 for 15 minutes to label DNA, and imaged using a spinning disk confocal microscope. Excitation wavelengths of 488 nm and 638 nm were used to visualize the mNeonGreen-tagged LEM2 and DNA, respectively, with z-stacks acquired to cover the entire nucleus (step size=250 nm).

Image analysis was performed in Imaris using a custom pipeline. The DNA volume was reconstructed into 3D surfaces using the Surfaces function with automatic thresholding (Fig. 3e) ensuring that the surfaces are properly captured without gaps and intrusions through manual inspection. To separate clumped nuclei, a seed diameter of 8 μm was applied. Nuclei that were incomplete in the z-stack or touched image boundaries were excluded, and additional filtering was applied to remove abnormally large or small nuclei (e.g., mitotic or defective) using surface area\sphericity filters. From the reconstructed surfaces, nuclear shape features including total surface area, volume, sphericity, eccentricity, prolate ellipticity, oblate ellipticity, and ellipsoid axis lengths were extracted and exported.

Data from multiple replicates were pooled and analysed using a custom MATLAB script. To calculate nuclear asphericity, the surface area of a sphere with the same volume as the nucleus (Av) was subtracted from the actual nuclear surface area (A) and the difference was normalized by Av for each nuclei.

Histograms of asphericity, sphericity, nuclear volume and surface area across knockdown conditions were plotted (Fig. 3f bottom, Extended Data Fig. 6i,j, Extended Data Fig. 8j,k, Extended Data Fig. 8l bottom). The raw data were fitted to a lognormal distribution using a non-linear least squares method, and key statistical parameters, including mean, median, mode, and peak position, were computed.

The cross-sectional area and circularity of nuclei were quantified from maximum projections of the DNA signal, using a threshold of 137 (range 1–255) for segmentation in Fiji. Area and circularity of segmented particles were directly measured and pooled from three replicates and compared across knockdowns or compression conditions (Extended Data Fig. 6g,h).

##### Q. Electron tomography and reconstruction

We used cryo-ET data acquired from FIB milled mouse embryonic stem-cell samples by the Back lab at Max Planck Institute of Biophysics, Frankfurt[44]. Cryo-ET data acquisition and reconstruction were performed by the Beck lab as described[44]. We analysed a total of three tomograms. To enhance the contrast and improve the isotropy, the tomogram was deconvolved, denoised and missing-wedge corrected by the deep-learning based software IsoNet [71]. Segmentation was performed using Dragonfly software (Version 2022.2 for Linux, developed by Object Research Systems (ORS) Inc, Montreal, Canada, 2020; www.theobjects.com/dragonfly)[72]. A small, representative region was selected from the full tomogram, containing key features of interest. Within this area, all voxels were manually segmented and assigned to one of seven classes: membrane, microtubules, nuclear lamina, ribosomes, proteasomes, chromatin, or background. This segmentation was then used to train a neural network. The Deep Learning Tool in Dragonfly was employed to create a multi-slice (five slices) U-Net. The resulting AI segmentations were manually cleaned and refined, with the nuclear lamina requiring a complete manual re-segmentation. Segmentations were exported as binary TIFF images and converted to MRC format using IMOD [73]. Surface smoothing and visualization were done in UCSF ChimeraX 1.8 [67] (Extended Data Fig. 5b-d).

The thickness of the hydrogel layer was measured as the distance between inner nuclear membrane and the mass of nucleosomes. The measurements were performed manually in IMOD across different tomogram slices and plotted as a histogram in Extended Data Fig. 5e. The mean thickness of the amorphous layer was calculated from the mean of the fitted normal distribution.

#### Section 6: AFM on cells: Nuclear stiffness measurements

##### A. Atomic force microscopy

The nucleus stiffness measurements were performed with a JPK Nanowizard 4XP AFM (Bruker) mounted on an LSM 700 confocal microscope (Zeiss). Can-tilevers with a quadratic pyramidal tip, a half-angle to face of 17.5°, and a nominal spring constant of 0.03 N/m (MLCT-D, Bruker) were used in all experiments. The spring constant was calibrated before each measurement session using the thermal fluctuation method (JPK SPM software, Bruker).

Cells were seeded in glass-bottom petri dishes (Fluorodish, FD35-100) two days before the measurements. One hour before the start of the measurements, the medium was changed to $CO_2$-independent DMEM (Invitrogen, #12800-017) containing 4 mM NaHCO3, buffered with 20 mM HEPES/NaOH pH 7.2 and supplemented with 10% fetal bovine serum (Life Technologies, #10270106). When latrunculin A (Invitrogen, #L12370) treatment was used, the drug was added to the medium 15 minutes before the start of the measurements to a final concentration of 200 nM. During all the AFM experiments, a petri dish heater (Bruker) was used to maintain the cells at 37°C.

For each measured cell, the nucleus was identified by eye using brightfield or differential interference contrast (DIC) microscopy. A 1 × 1 $\mu m^2$ region at the nuclear center was selected using the acquisition software of the atomic force microscope (AFM; JPK SPM, Bruker), and a 3 × 3 grid of measurement points was overlaid. Force mapping was performed in contact mode with a setpoint force of 1 nN and an indentation speed of 1 µm/s, yielding nine indentation curves per cell. For experiments with latrunculin A, a reduced setpoint force of 0.5 nN was used due to increased cell softness. In the LEM2 knockdown condition, confocal microscopy images of mNeonGreen (excitation: 488 nm) were acquired before AFM measurements. Cells in which LEM2-mNeonGreen localized to the nuclear membrane were excluded from indentation, as this indicated incomplete knockdown.

##### B. Analysis of the AFM data

The indentation curves were first fitted using the AFM manufacturer's software JPK Data Processing (Bruker). In brief, two analysis steps were performed: (i) determining the contact point through Hertz model fitting, and (ii) estimating an apparent Young's modulus through performing a Hertz fit in a defined indentation range of 0.5 µm. A Poisson's ratio of 0.5 was assumed.

In detail, as part of the first analysis step, any tilt or offset in the curve was corrected using the curve's baseline as reference. At this point, curves with no clear baseline were discarded as they denote failed indentation experiments. The Hertz-Sneddon model was then fitted on the part of the indentation curve where the force was below 100 pN, to ensure a good fit. The obtained fit contact point was kept for the next analysis step, but the fitted Young's modulus was disregarded. In the second analysis step, the Hertz-Sneddon model was fitted again to corrected force-indentation curves, this time with a restriction to a 0.5 µm indentation depth, using the previously fitted contact point as a fixed parameter. The Young's modulus obtained in this second step was kept as the measured apparent Young's modulus of the corresponding indentation. Finally, the median Young's modulus across the 9 indentations was calculated for each cell. The values were pooled from three replicates and the comparison across knockdowns was plotted in Extended Data Fig. 10g.

The analysis of the distribution of rupture events in the indentation curves was carried out with a custom MATLAB code. First, the part of the indentation curve ranging from the fitted contact point to the point of maximum force was smoothed using a low-pass filter, and the segments of the smoothed curve with decreasing force were considered. Each segment where the force monotonically decreased by more than 50 pN was marked as a rupture event. Finally, the fraction of curves showing at least one rupture event was calculated for each cell (Extended data Fig. 10e,f).

The effective area modulus of nuclei across knockdowns was determined from individual AFM force-indentation curves in the presence of latrunculin A to remove cytoskeletal interference. The work done during indentation from 0 to 2 μm was calculated by integrating the area under the AFM force-indentation curve (Fig. 3b, Extended Data Fig. 8a) using trapezoidal method of numerical integration. The nuclear surface area change induced by indentation was quantified through 3D segmentation and surface fitting of volumetric images of nuclei acquired at 0 and 2 μm indentations, as previously described (Fig. 3a, methods: nuclear surface and shape calculations). The effective area modulus ($\lambda_{eff}$) was then calculated using the following equation, incorporating the work done ($W$), the initial surface area at 0 μm indentation ($A$), and the area change upon indentation ($\Delta A$):

$$\lambda_{eff} = \frac{2WA}{\Delta A^2}$$

The value of $\frac{A}{\Delta A^2}$ was calculated for six different cells as in Extended Data Fig. 10b and the mean of the values was used in the above equation.The histogram of $\lambda_{eff}$ followed a lognormal distribution. The lognormal fit provided estimates for the mean, median, and mode of the distribution (Extended Data Fig. 10c,d). The calculated work done and $\lambda_{eff}$ values were plotted as a scatter of individual points across conditions with mean ± s.d. (Fig. 3c,d Extended Data Fig. 8b,c).

Averaged force-indentation curves were obtained by aligning individual curves to the contact point using a custom Python script. Only curves that reached an indentation of 2 μm were used. The mean ± s.d. of force at each indentation depth was then computed from the aligned curves and plotted in Extended Data Fig. 10a.

**Statistical analysis and sample numbers**

All experiments were repeated several times and indicated experiment numbers always refer to biological replicates for cell experiments and technical replicates for in-vitro experiments. Data were tested for normality or lognormality using Kolmogorov–Smirnov test. The appropriate statistical test was chosen as follows: Unpaired not normal distributed data containing only two groups: Mann-Whitney test, unpaired normal distributed data containing two or more groups: One-Way ANOVA with Tukey's test to correct for multiple comparisons, unpaired lognormal data with two groups: Welch's t-test and unpaired not normal distributed data containing two or more groups: Kruskal Wallis test with multiple comparisons using Dunn's test. $\alpha = 0.05$ for all hypothesis testing. *$p < 0.05$, **$p < 0.005$, ***$p < 0.001$, ****$p < 0.0001$, ns: non-significant.

### Supplementary Note 1

#### I. SIMPLE MODEL FOR DNA ELASTICITY UPON PROTEIN BINDING

In this section, we describe a simple model for how BAF-LEM2 binding affects the mechanical properties of DNA.

##### A. Statistical mechanics of molecules binding to elastic elements

In the enthalpic regime, DNA behaves as a Hookean spring with rest length $X_0$, and its end-to-end distance is denoted by $X$. Each bound BAF-LEM2 module spans a segment of length $X/N$ on DNA, where $N$ is the total number of binding sites of BAF-LEM2 module. The energy of the DNA binding state with end-to-end distance $X$ and $n$ BAF-LEM2 modules bound is:

$$E_n(X) = \epsilon\, n + \frac{1}{2}K_n(X - X_0)^2 - f(X - X_0) \ , \tag{1}$$

where $\epsilon$ denotes the binding energy of BAF-LEM2 module, $K_n$ is the effective stiffness of this binding state, and $f$ represents the external tension applied to DNA. The effective stiffness $K_n$ arises from $N - n$ DNA segments, each spring of stiffness $k_0$, and $n$ segments of stiffness $k_0 + k_i$, formed by a DNA spring ($k_0$) in parallel with a BAF–LEM2 spring ($k_i$):

$$\frac{1}{K_n} = \frac{N - n}{k_0} + \frac{n}{k_0 + k_i} \ . \tag{2}$$

The chemical potential of the BAF–LEM2 module is given by

$$\mu = \mu_0 + \beta^{-1}\ln(c_L/c_0^L) + \beta^{-1}\ln(c_B/c_0^B), \tag{3}$$

where $\mu_0$ is a reference chemical potential, $\beta = 1/(k_BT)$, $c_L$ and $c_B$ are the concentrations of LEM2 monomers and BAF dimers, respectively, and $c_0^L$ and $c_0^B$ are reference concentrations. The expression for chemical potential holds under the assumption of ideal dilute behavior at equilibrium, in which BAF–LEM2 modules form from LEM2 molecules and BAF dimers according to

$$\text{BAF-LEM2 module} \ \rightleftharpoons\ \text{LEM2} + \text{BAF}_{\text{dimer}} \ . \tag{4}$$

For simplicity, we write

$$\mu = \beta^{-1}\ln(c_L/c_0) \ , \tag{5}$$

where $c_0 = c_0^L(c_0^B/c_B)e^{-\beta\mu_0}$ is the reference concentration that depends on BAF concentration. The grand-canonical partition function is:

$$Z(\mu, f, N) = \sum_{n=0}^{N}\int_{-\infty}^{\infty} dX \frac{N!}{n!(N-n)!}e^{-\beta(E_n(X)-\mu n)}, \tag{6}$$

where the combinatorial factor $\binom{N}{n} = N!/[n!(N-n)!]$ accounts for the number of ways to bind $n$ BAF-LEM2 modules. We define distances per binding site:

$$x_0 = \frac{X_0}{N}, \quad x = \frac{X}{N}, \tag{7}$$

and the occupation fraction $p$ according to:

$$p = \frac{n}{N}, \tag{8}$$

The partition function in the large $N$ limit, up to an overall prefactor and using the Stirling approximation, becomes

$$Z(\mu, f, N) = \int_0^1 dp \int_{-\infty}^{\infty} dx\, e^{-\beta N g(x,p)}, \tag{9}$$

where

$$g(x,p) = \beta^{-1}[p\ln p + (1-p)\ln(1-p)] + (\epsilon-\mu)p + \frac{1}{2}\frac{k_0}{1-\frac{k_i}{k_0+k_i}p}(x-x_0)^2 - f(x-x_0)\ . \tag{10}$$

In the thermodynamic limit of a long DNA molecule, number of binding sites $N$ is large. We can apply the saddle point approximation in Eq. (9). For large $N$, the integral is dominated by the saddle point $(\bar{x}, \bar{p})$, where $g(x,p)$ is extremized. The extremum conditions

$$\left.\frac{\partial g(x,p)}{\partial x}\right|_{\bar{x},\bar{p}} = 0\ , \qquad \left.\frac{\partial g(x,p)}{\partial p}\right|_{\bar{x},\bar{p}} = 0 \tag{11}$$

yield closed-form solutions for $\bar{x}$ and $\bar{p}$ as the most probable end-to-end distance and occupation fraction, respectively, at equilibrium:

$$\bar{x} = x_0 + \frac{1}{k_0}\Big(1 - \frac{1}{1+k_0/k_i}\bar{p}(f)\Big)f\ , \qquad \bar{p}(f) = \frac{1}{1+e^{\beta(\epsilon-\mu+f^2/[2k_0(1+k_0/k_i)])}}. \tag{12}$$

These are shown as a function of $f$ in Extended Data Fig. 4d. In the limit $N\to\infty$, these coincide exactly with the ensemble averages. For finite $N$, they remain accurate when fluctuations are small. The average stretch $\bar{x}-x_0$ as a function of $f$ has linear regimes at large $f \gg \sqrt{k_0/\beta}$:

$$\bar{x} - x_0 = k_0^{-1} f\left[1 + \mathcal{O}\left(e^{-\gamma(f/\sqrt{k_0/\beta})^2}\right)\right]\ , \qquad \bar{p}(f) = \mathcal{O}\left(e^{-\gamma(f/\sqrt{k_0/\beta})^2}\right) \tag{13}$$

and at small $f \ll \sqrt{k_0/\beta}$

$$\bar{x} - x_0 = k(\bar{p})^{-1} f + \mathcal{O}\left(\left[f/\sqrt{k_0/\beta}\right]^3\right)\ , \qquad \bar{p}(f) = \frac{1}{1+e^{\beta(\epsilon-\mu)}} + \mathcal{O}\left(\left[f/\sqrt{k_0/\beta}\right]^2\right), \tag{14}$$

where

$$k(\bar{p}) = k_0\Big(1 - \frac{1}{1+k_0/k_i}\bar{p}\Big)^{-1} \tag{15}$$

and $\gamma = [2(1+k_0/k_i)]^{-1}$.

##### B. Comparison to single molecule experiments

DNA is held at constant tension $f$ via a feedback loop that constantly adjusts optical trap positions and therefore DNA end-to-end distance $\bar{X}$. The end-to-end distance $\bar{X}$ depends on the BAF-LEM2 module occupation fraction $\bar{p}$ on DNA according to Eq. (14):

$$\bar{X}(\bar{p}) = X_0 + K(\bar{p})^{-1} f + \mathcal{O}\left(\left[f/\sqrt{k_0/\beta}\right]^3\right)\ , \tag{16}$$

where we used $\bar{X} = N\,\bar{x}$ and defined $K(\bar{p}) = k(\bar{p})/N$. Adding LEM2 results in a reduction of the end-to-end distance

$$\bar{X}(0) - \bar{X}(\bar{p}) = \Big(K(0)^{-1} - K(\bar{p})^{-1}\Big)f + \mathcal{O}\left(\left[f/\sqrt{k_0/\beta}\right]^3\right)\ . \tag{17}$$

Substituting $\mu$ from Eq. (5) into the differences of compliances $K(0)^{-1} - K(\bar{p})^{-1}$, we have

$$K(0)^{-1} - K(\bar{p})^{-1} = \frac{N/k_0}{1+k_0/k_i}\frac{1}{1+e^{\beta\epsilon}c_0/c_L}\ . \tag{18}$$

Equation (17) is fit to the shortening data as a function of tension at different LEM2 concentrations $c_L$ (Extended Data Fig. 4e,f), where $K(0)^{-1} - K(\bar{p})^{-1} = B_1/(1+B_2/c_L)$ is used and $B_1$ and $B_2$ are the fit parameters, with $B_1 = N(1+k_0/k_i)^{-1}k_0^{-1}$ and $B_2 = e^{\beta\epsilon}c_0$. The best fit values are given in Table I. We will use the effective springs $k(\bar{p})$ to build an elastic continuum model of a nuclear shell.

#### II. NUCLEAR SHAPE GOVERNED BY AN ELASTIC SURFACE HYDROGEL

Here we define the rest-length mismatched surface hydrogel model of the cell nucleus discussed in the main text.

##### A. Elastic energy of the nuclear shell

We describe the nucleus as a thin elastic shell enclosing a fixed volume $V$, following the microscopic framework introduced above. The total free energy combines (i) bending, (ii) isotropic surface compression (no shear), and (iii) elastic contribution from BAF-LEM2 modules governed by the effective stiffness $k(\bar{p})$:

$$F = \int dA \, \frac{\kappa}{2} \left(2H - C_0\right)^2 + \int dA_* \left( \frac{\lambda}{2} u^2 + \frac{k(\bar{p})}{2} \left( \frac{x(u) - x_0}{\ell_*} \right)^2 \right) , \tag{19}$$

where $A$ is the total area, $\kappa$ the bending rigidity, $H$ the mean curvature, $C_0$ the spontaneous curvature, $A_*$ the strain-free area, $\lambda$ the bare area elastic modulus of (lamin) gel in the absence of DNA-BAF-LEM2 modules, $u \equiv u_{kk}$ denotes the trace of the strain tensor $u_{ij} = 1/2(\partial_i u_j + \partial_j u_i)$. Here, summation over repeating indices is assumed. Parameter $\bar{p}$ is the fraction of BAF-LEM2 modules bound to DNA, $k(\bar{p})$ the effective stiffness of DNA-BAF-LEM2 module, $x$ the end-to-end distance of BAF-LEM2 module, $x_0$ DNA rest length per BAF-LEM2 module, $1/\ell_*^2$ the surface density BAF-LEM2 module binding sites in the strain-free state. Using Gauss theorem, we rewrite the area integral of the divergence $u_{kk}$ as the flux of the field $\mathbf{u}$ through the boundary of the area element:

$$\int dA_* \, u_{kk} = \oint d\mathbf{l} \cdot \mathbf{u} = A - A_* \ . \tag{20}$$

For simplicity, we consider isotropic strain $u$ to be uniform:

$$u \simeq \frac{A - A_*}{A_*} \ , \tag{21}$$

which relates the surface area $A$ to the strain-free area $A_*$:

$$A \simeq (1 + u) A_* \ . \tag{22}$$

The end-to-end distance of a BAF-LEM2 module under strain is thus given by:

$$x(u) \simeq \sqrt{1 + u} \, x_* \ , \tag{23}$$

where $x_* = x(0)$ is the end-to-end distance in the strain-free state $u = 0$. In the following, we neglect the contribution of the bending energy, because $\kappa/(\lambda A_*) \sim 10^{-7}$, see Section III. To determine the free energy as a function of $u$, we use Eqs. (19) and (23) to obtain:

$$F(u) = \int dA_* \left[ \frac{\lambda}{2} u^2 + \frac{k(\bar{p})}{2} \left( \frac{x_0}{l_*} \right)^2 \left( \frac{\sqrt{1 + u}}{1 + v_0} - 1 \right)^2 \right] , \tag{24}$$

where we introduced a dimensionless rest-length mismatch parameter $v_0 = x_0/x_* - 1$. We define the dimensionless free energy

$$F/F_0 = \frac{1}{2} u^2 + \frac{1}{2} \rho \Big( \frac{\sqrt{1 + u}}{1 + v_0} - 1 \Big)^2 , \tag{25}$$

where $F_0 = \lambda A_*$, and we introduce the dimensionless parameter $\rho$, which is proportional to BAF-LEM2 module binding sites surface density $1/\ell_*^2$ in the strain-free state:

$$\rho = \frac{k(\bar{p})}{\lambda} \left( \frac{x_0}{\ell_*} \right)^2 \ . \tag{26}$$

We define the asphericity

$$\Delta = A/A_V - 1 \ , \tag{27}$$

where $A$ is the surface area and $A_V = (6\sqrt{\pi}\, V)^{2/3}$, the area of a sphere with volume $V$. We can now express strain $u$ (Eq. (21)) as a function of $\Delta$,

$$u = u_0 + (1 + u_0)\Delta \ , \tag{28}$$

where

$$u_0 = A_V/A_* - 1 \tag{29}$$

is a reference strain, corresponding to the strain achieved for a spherical shape.

##### B. Free energy for parameterized shapes

We consider a simple axisymmetric parametrization of the shape close to a sphere (Extended Data Fig. 7a)

$$R(\theta) = \bar{r}(\alpha, V)\Big[1 + \alpha Y_{20}(\theta)\Big] \ , \tag{30}$$

where $\theta$ is the polar angle, $\alpha$ the shape parameter, and $Y_{20}(\theta) = \sqrt{5/(16\pi)}(3\cos^2\theta - 1)$ the normalized ($\ell = 2, m = 0$) spherical harmonic. The shape parameter $\alpha$ controls whether the shape is oblate ($\alpha < 0$, flattened at the poles), spherical ($\alpha = 0$), or prolate ($\alpha > 0$, elongated along the polar axis). We impose constant volume $V$ when $\alpha$ changes. This fixes the function $\bar{r}(\alpha, V)$, see Section III, Eq. (50).

In order to calculate the free energy $F(\alpha)$ as a function of the shape parameter $\alpha$, we first determine the asphericity $\Delta$ as a function of $\alpha$, see Extended Data Fig. 7b. This function is determined in Section III, Eq. (55). The normalized free energy $F/F_0$ defined in Eq. (25) is shown as a function of $\alpha$ in Extended Data Fig. 7c–f. For simplicity, we restrict our analysis to oblate shapes. We denote $\alpha(u_0, v_0, \rho) < 0$ to be the value of $\alpha$ that minimizes the free energy $F(\alpha)$ for given $u_0$, $v_0$, and $\rho$.

##### C. Phase diagrams of nuclear shapes

We construct two phase diagrams, one on the $u_0 - \rho$ plane and one on the $v_0 - \rho$ plane (Extended Data Fig. 7g,h). We determine the critical line that separates the region of spherical shapes (Extended Data Fig. 7g,h, no shading; $\alpha = 0$) from the region of oblate shapes (Extended Data Fig. 7g,h, color shading, $\alpha < 0$). We express $u$ (Eq. (28)) in powers of $\alpha$, using $\Delta(\alpha)$ specified in Section III, Eq. (60):

$$u = u_0 + (1 + u_0)\left(\frac{3}{2\pi}\,\alpha^2 + \frac{\sqrt{5}}{42\,\pi^{3/2}}\,\alpha^3 + O(\alpha^5)\right) \ . \tag{31}$$

We next use this expression to expand $F(\alpha)/F_0$ (Eq. (25)) in powers of $\alpha$:

$$\begin{aligned} F(\alpha)/F_0 = {} & \frac{1}{2}\left[u_0^2 + \rho\left(\frac{\sqrt{1+u_0}}{1+v_0} - 1\right)^2\right] \\ & + \frac{3}{4\pi}\left[2u_0(1+u_0) + \rho\frac{\sqrt{1+u_0}}{1+v_0}\left(\frac{\sqrt{1+u_0}}{1+v_0} - 1\right)\right]\alpha^2 \\ & - \frac{\sqrt{5}}{84\pi^{3/2}}\left[2u_0(1+u_0) + \rho\frac{\sqrt{1+u_0}}{1+v_0}\left(\frac{\sqrt{1+u_0}}{1+v_0} - 1\right)\right]\alpha^3 + \mathcal{O}(\alpha^4) \ . \end{aligned} \tag{32}$$

The system will evolve until $F(\alpha)/F_0$ is at a minimum. We next determine the critical density $\rho = \rho_c$ at which the system undergoes a continuous (second-order) shape bifurcation (supercritical pitchfork) between oblate and prolate shapes. By imposing inflection point conditions $\partial^2 F/\partial\alpha^2 = 0$ and $\partial^3 F/\partial\alpha^3 = 0$, we find

$$\rho_c = \frac{2u_0(1+v_0)^2\sqrt{1+u_0}}{1 + v_0 - \sqrt{1+u_0}} \ , \tag{33}$$

displayed as a solid black line in Extended Data Figs. 7g,h. Note that $\rho_c$ diverges at $u_0^c = 2v_0 + v_0^2$ (dashed lines, Extended Data Fig. 7g,h). For small $\rho$, one finds $v_0 \approx -1 + \frac{1}{2}\sqrt{-2\rho/u_0}$ (Extended Data Fig. 7h).

##### D. Effective elastic area modulus

Here we derive the effective area elastic modulus, defined as an elastic response of $F(\alpha)/F_0$ upon a change in area

$$\lambda_{\text{eff}} = \frac{1}{A_*} \frac{\partial^2 F}{\partial u^2}\bigg|_{u=\bar{u}} \ , \tag{34}$$

where $\bar{u}$ is the equilibrium strain, $\partial F/\partial u|_{u=\bar{u}} = 0$. Eq. (25) in dimensional form becomes

$$F = \frac{\lambda}{2} u^2 A_* + \frac{\rho}{2} \lambda \Big( \frac{\sqrt{1+u}}{1+v_0} - 1 \Big)^2 A_* \ , \tag{35}$$

resulting in an effective area modulus

$$\lambda_{\text{eff}} = \lambda \left( 1 + \frac{\rho}{4(1+\bar{u})^{3/2}(1+v_0)} \right) \ . \tag{36}$$

##### E. Comparison to measured shapes in siLEM2 perturbation

We measured nuclear shapes in two conditions, siCtrl (C), corresponding to the presence of BAF-LEM2 modules ($\bar{p}_C = 1$ is assumed), and siLEM2 (L), corresponding to a knockdown of LEM2 by $1 - \bar{p}_L = 0.74$ (Extended Data Fig. 6d). For the parameters $u_0 = -0.43$ and $v_0 = -0.25$ (Section II G) used in the main text, the normalized free energy for oblate shapes $\alpha < 0$ is shown in Fig. 3f as a function of asphericity $\Delta$. Two values of $\rho$ are considered, corresponding to the two experimental conditions siCtrl (C) with $\rho_C = 13.3$ and siLEM2 (L) with $\rho_L = 9.9$.

The asphericities $\Delta_C$ and $\Delta_L$ correspond to the peaks of the measured distributions in siCtrl and siLEM2 conditions, respectively (Fig. 3f, bottom). We numerically determine values $u_0$ and $v_0$ that satisfy

$$\begin{cases} \Delta\Big[\alpha\big(u_0, v_0, \rho_C\big)\Big] = \Delta_C \\ \Delta\Big[\alpha\big(u_0, v_0, \rho_L\big)\Big] = \Delta_L \end{cases} , \tag{37}$$

where we used Eq. (55) to calculate $\Delta$ as a function of $\alpha$. To this end we express the ratio of $\rho_C$ and $\rho_L$ using Eq. (26) as

$$\frac{\rho_C}{\rho_L} = \xi_L \, \lambda^L / \lambda^C \ , \tag{38}$$

where stiffness ratio $\xi_L = k(\bar{p}_C)/k(\bar{p}_L)$ of the two conditions with occupation fraction $\bar{p}_C$ and $\bar{p}_L$ is:

$$\xi_L = \frac{1 + k_0/k_i - \bar{p}_L}{1 + k_0/k_i - \bar{p}_C}. \tag{39}$$

Here, the stiffness ratio $\xi_L \simeq 1.34$ (95% CI 1.27–1.46), Table II. We used that $x_0$ is independent of BAF-LEM2 module occupation fraction in optical tweezer experiments (Fig. 2d). Extended Data Fig. 7i shows numerical solutions of Eq. (37) for $\lambda^L/\lambda^C = 1$.

##### F. Comparison to measured area moduli in siLEM2 perturbation

Using Eq. (36), we obtain a ratio of the effective area moduli

$$\frac{\lambda^C_{\text{eff}}}{\lambda^L_{\text{eff}}} = \frac{1 + \eta_C \rho_C}{1 + \eta_L \rho_L} \frac{\lambda^C}{\lambda^L} \ , \tag{40}$$

where $\eta_C = (4(1+\bar{u}_C)^{3/2}(1+v_0))^{-1}$ and $\eta_L = (4(1+\bar{u}_L)^{3/2}(1+v_0))^{-1}$, $\lambda^C$ and $\lambda^L$ denote bare area moduli in siCtrl and siLEM2 conditions, respectively, $\bar{u}_C$ and $\bar{u}_L$ are the corresponding equilibrium strains. Using

$$\rho_L = \frac{k(\bar{p}_L)}{\lambda^L} \left( \frac{x_0}{\ell_*} \right)^2 = \xi_L^{-1} \frac{\lambda^C}{\lambda^L} \, \rho_C \ , \tag{41}$$

where $\xi_L = k(\bar{p}_C)/k(\bar{p}_L)$, we find

$$\frac{\lambda^C_{\text{eff}}}{\lambda^L_{\text{eff}}} = \frac{1+\eta_C\rho_C}{\lambda^L/\lambda^C + \eta_L\xi_L^{-1}\rho_C} \ . \tag{42}$$

We further connect this relationship with the dimensional variable $\lambda^C$, shown in Extended Data Fig. 7l, where we used $\rho_C = \left(k(\bar{p}_C)/\lambda^C\right)(x_0/\ell_*)^2$, $\ell_* = \ell_C/\sqrt{1+\bar{u}_C}$, $\ell_C \simeq 79$ nm (Table 1 of the Main Text). We estimate $x_0$ using Eq. (7), where $X_0 \simeq 15.84\,\mu$m (Fig. 2d) and $N \simeq 8500$ from the saturating value of the titration curve on Extended Data Fig. 4l. Here we used $\lambda_L = \lambda_C$ which is supported by experimental evidence [2].

##### G. Incorporating siBAF data and selecting parameters

We conduct the same procedure as described for siLEM2 in Sections II E and F for siBAF (B) condition, see Table II and Extended Data Fig. 8d-g. Here, we assume that a decrease in the number of BAF dimers leads to a proportional decrease in BAF-LEM2 occupation fraction $\bar{p}_B$.

The intercept between the model CI band and the AFM uncertainty range (Extended Data Fig. 7l and 8g) yields the uncertainty regions in $u_0, v_0$ space for siLEM2 and siBAF respectively (Extended Data Fig. 8h). These regions overlap and provide a common choice of surface hydrogel parameters to account for both conditions. Specifically, the model agrees with siLEM2 data for $\lambda^C < 0.2$ pN/nm, corresponding to $\rho_C > 2$, (Extended Data Fig. 7l); and with siBAF data for $\lambda^C < 0.014$ pN/nm corresponding to $\rho_C > 13.3$ (Extended Data Fig. 8g). We therefore choose $\rho_C = 13.3$ that satisfies both conditions (vertical dashed line on Extended Data Fig. 7i-l and Extended Data Fig. 8d-g). This results in $u_0 = -0.43$ and $v_0 = -0.25$. We conclude that our surface hydrogel model accounts for the observed changes in both the area elastic modulus and nuclear shape in siLEM2 and siBAF.

We also note that depletion of BAF causes changes to the elastic properties of the lamin network due to the reduction in its crosslinks [10,55]. By incorporating this change, the model shows enlarged parameter range to account experimental observations (Extended Data Fig. 8g).

#### III. GEOMETRY OF DEFORMED SURFACES

In this section, we derive the geometric relations used in Section II.

##### A. Volume of axisymmetric shapes

We parametrize axisymmetic shapes as

$$R(\theta) = R_0(1+\alpha Y_{20}(\theta)) \ , \tag{43}$$

see Extended Data Fig. 7a. The volume can be expressed as

$$V = \int_0^\pi d\theta \int_0^{2\pi} d\phi \int_0^{R(\theta)} r^2 \sin\theta \, dr \ . \tag{44}$$

This reduces to a single integral over the polar angle $\theta$:

$$V = \frac{2\pi}{3}\int_0^\pi R(\theta)^3 \sin\theta \, d\theta \ . \tag{45}$$

Using the shape parametrization (Eq. (30)), we find

$$R^3(\theta) = R_0^3[1 + 3\alpha Y_{20}(\theta) + 3(\alpha Y_{20}(\theta))^2 + (\alpha Y_{20}(\theta))^3] \ . \tag{46}$$

We note that

$$\int_0^\pi Y_{20}(\theta)\,\sin\theta\,d\theta = 0 \ , \qquad \int_0^\pi Y_{20}^2(\theta)\,\sin\theta\,d\theta = 1 \tag{47}$$

and

$$\int_0^\pi \big[Y_{20}(\theta)\big]^3 \sin\theta\, d\theta = \Big(\sqrt{\frac{5}{16\pi}}\Big)^3 \int_{-1}^1 (3u^2-1)^3\, du = \frac{\sqrt{5}}{14\,\pi^{3/2}} \ , \tag{48}$$

where $u = \cos\theta$. With this, the volume integral (Eq. (45)) becomes:

$$V = \bar{R}_0^3\Big(\frac{4\pi}{3} + \alpha^2 + \frac{1}{21}\sqrt{\frac{5}{\pi}}\alpha^3\Big) \ . \tag{49}$$

We impose fixed volume $V$ when changing the shape with $\alpha$. This implies $R_0 = \bar{r}(\alpha, V)$, with

$$\bar{r}(\alpha, V) = \left(1 + \frac{3}{4\pi}\alpha^2 + \frac{1}{28}\frac{\sqrt{5}}{\pi^{3/2}}\alpha^3\right)^{-1/3} \sqrt{\frac{A_V}{4\pi}} \ , \tag{50}$$

where $A_V = (6\sqrt{\pi}\,V)^{2/3}$ is the area of a sphere with volume $V$.

##### B. Area of axisymmetric shapes

The surface area of an axisymmetric shape can be written as

$$A = 4\pi \int_0^\pi \frac{R(\theta)^2}{2}\sqrt{1 + \frac{1}{R(\theta)^2}\left(\frac{\partial R(\theta)}{\partial\theta}\right)^2}\,\sin\theta\, d\theta \ . \tag{51}$$

For the axisymmetric shape parametrized by Eq. (43), this integral becomes:

$$A = 4\pi R_0^2\, I(\alpha) \ , \tag{52}$$

where $I(\alpha)$ is:

$$I(\alpha) = \int_0^\pi \frac{(1+\alpha Y_{20}(\theta))^2}{2}\sqrt{1 + \frac{\alpha^2\, Y_{20}'(\theta)^2}{(1+\alpha Y_{20}(\theta))^2}}\,\sin\theta\, d\theta \ , \tag{53}$$

and $Y_{20}'(\theta)$ denotes the derivative of $Y_{20}(\theta)$ with respect to $\theta$. The surface area $A(\alpha)$, enclosing $V$ thus reads from Eq. (52):

$$A(\alpha, V) = A_V \left(1 + \frac{3}{4\pi}\alpha^2 + \frac{\sqrt{5}}{28\pi^{3/2}}\alpha^3\right)^{-2/3} I(\alpha) \ , \tag{54}$$

where we implied $R_0 = \bar{r}(\alpha, V)$. The asphericity defined in Eq. (27) for the shape parametrization in Eq. (30) is:

$$\Delta(\alpha) = I(\alpha)\left(1 + \frac{3}{4\pi}\alpha^2 + \frac{\sqrt{5}}{28\pi^{3/2}}\alpha^3\right)^{-2/3} - 1 \ , \tag{55}$$

which is shown as a function of $\alpha$ in Extended Data Fig. 7b.

##### C. Perturbative expansion in $\alpha$

We now determine the approximate surface area of the shapes defined in Eq. (30) by expanding $I(\alpha)$ in powers of $\alpha$:

$$\begin{aligned} I(\alpha) &= \int_0^\pi \Big[\frac{1}{2} + \alpha\, Y_{20}(\theta) + \alpha^2\Big(\frac{1}{2}Y_{20}(\theta)^2 + \frac{1}{4}Y_{20}'(\theta)^2\Big)\Big] \sin\theta\, d\theta \, + \mathcal{O}(\alpha^4) \\ &= 1 + \frac{\alpha^2}{\pi} + \mathcal{O}(\alpha^4) \ , \end{aligned} \tag{56}$$

where we have used Eq. (47) as well as following the integrals:

$$\int_0^\pi \left[Y_{20}'(\theta)\right]^2 \sin\theta \, d\theta = 36\Big(\sqrt{\frac{5}{16\pi}}\Big)^2 \int_{-1}^1 u^2(1-u^2)\, du = \frac{3}{\pi} \ . \tag{57}$$

Substituting this newly-obtained expression for $I(\alpha)$ into Eq. (52), we find:

$$\frac{A}{4\pi R_0^2} = 1 + \frac{\alpha^2}{\pi} + \mathcal{O}(\alpha^4) \ . \tag{58}$$

We obtain the surface area $A(\alpha)$ for shapes enclosing the volume $V$ by substituting $\bar{r}(\alpha, V)$ from Eq. (50) into Eq. (52):

$$\begin{aligned} \frac{A(\alpha)}{A_V} &= \left(1 + \frac{3}{4\pi}\alpha^2 + \frac{1}{28}\frac{\sqrt{5}}{\pi^{3/2}}\alpha^3\right)^{2/3} \left(1 + \frac{\alpha^2}{\pi} + \mathcal{O}(\alpha^4)\right) \\ &= 1 + \frac{3}{2\pi}\,\alpha^2 + \frac{\sqrt{5}}{42\,\pi^{3/2}}\,\alpha^3 + \mathcal{O}(\alpha^3) \ . \end{aligned} \tag{59}$$

This expression leads directly to an approximate form for $\Delta(\alpha)$:

$$\Delta(\alpha) = \frac{3}{2\pi}\,\alpha^2 + \frac{\sqrt{5}}{42\,\pi^{3/2}}\,\alpha^3 + \mathcal{O}(\alpha^3) \ , \tag{60}$$

which we employ in phase diagram calculations.

##### D. Bending energy

Here, we calculate the full free energy Eq. (19) with bending energy term. The bending energy

$$E_\kappa = \int \frac{\kappa}{2}(2H - C_0)^2 dA \ , \tag{61}$$

can be calculated for shape perturbations close to the sphere

$$R(\theta) = \bar{r}(\alpha, V)\Big[1 + \sum_{\ell=2}^{l_{\max}} \sum_{m=-\ell}^{\ell} \alpha_{\ell,m}\, Y_{\ell m}(\theta,\phi)\Big] \tag{62}$$

and reads

$$E_\kappa^{\ell,m} \simeq 8\pi\,\kappa + \frac{1}{2}\,\kappa\,|\alpha_{l,m}|^2(\ell+2)(\ell+1)\ell(\ell-1) \tag{63}$$

[1, 3], which becomes $E_\kappa \simeq 8\pi\,\kappa + 12\,\kappa\,\alpha^2$ for $\ell = 2, m = 0$.

The normalized free energy reads

$$F/F_0 = \int dA_* \Big[\frac{\tilde{\kappa}}{2}(2H - C_0)^2 + \frac{1}{2}\frac{u^2}{A_*} + \frac{1}{2}\frac{\rho}{A_*}\Big(1 - \frac{\sqrt{1+u}}{1+v_0}\Big)^2\Big] \ , \tag{64}$$

where $\tilde{\kappa} = \kappa/(\lambda A_*)$. Combining Eqs. (32) and (63), we expand the free energy up to second order in $\alpha$:

$$\begin{aligned} F(\alpha)/F_0 &= \frac{1}{2}\left[16\pi\tilde{\kappa} + u_0^2 + \rho\left(\frac{\sqrt{1+u_0}}{1+v_0} - 1\right)^2\right] \\ &+ \frac{3}{4\pi}\left[16\pi\tilde{\kappa} + 2u_0(1+u_0) + \rho\frac{\sqrt{1+u_0}}{1+v_0}\left(\frac{\sqrt{1+u_0}}{1+v_0} - 1\right)\right]\alpha^2 + \mathcal{O}(\alpha^3) \ . \end{aligned} \tag{65}$$

We now estimate the bending rigidity of the surface hydrogel. The bending rigidity of the thin shell with thickness $h$ can be estimated to be $\kappa \simeq \lambda h^2/12$. Thus, $\tilde{\kappa} \simeq \frac{1+u_0}{(6\sqrt{\pi})^{2/3}}\frac{h^2}{V^{2/3}} \simeq 10^{-7}$, where $h \simeq 14\,\text{nm}$ is the thickness of the hydrogel (Extended Data Fig. 5e) and $V \simeq 10^3 \mu m^3$ nuclear volume (Extended Data Fig. 6j).

| Parameter | Name | Equation | Value | 95% CI | Units |
|---|---|---|---|---|---|
| $B_1$ | Shortening amplitude | $N/k_0\,(1+k_0/k_i)^{-1}$ | 6.2 | (5.3, 7.6) | nm/pN |
| $B_2$ | Reference parameter | $c_0\,e^{\beta\epsilon}$ | 105 | (65, 190) | nM |

TABLE I: Parameters from the 2-parameter fit of shortening (Extended Data Fig. 4e,f). Summary of parameter estimates and 95% confidence intervals (CI), computed from $10^5$ bootstrap replicates.

| Parameter | Equation | Value | 95% CI |
|---|---|---|---|
| DNA-BAF-LEM2 module to DNA stiffness ratio | $(k_0+k_i)/k_0$ | 1.46 | (1.36, 1.62) |
| stiffness ratio $\xi_L = k(\bar{p}_C)/k(\bar{p}_L)$ | $\frac{1+k_0/k_i-\bar{p}_L}{1+k_0/k_i-\bar{p}_C}$ | 1.34 | (1.27, 1.46) |
| stiffness ratio $\xi_B = k(\bar{p}_C)/k(\bar{p}_B)$ | $\frac{1+k_0/k_i-\bar{p}_B}{1+k_0/k_i-\bar{p}_C}$ | 1.41 | (1.33, 1.56) |

TABLE II: Estimated parameters and propagated 95% CI, corresponding to Table 1. Parameters used: $k_0/N \simeq 50.2$ pN/$\mu$m (Fig. 2e), $\bar{p}_C = 1$, $\bar{p}_B \simeq 0.1$ (Extended Data Fig. 6b), $\bar{p}_L \simeq 0.26$ (Extended Data Fig. 6d).

| Parameter | Symbol | Value |
|---|---|---|
| Reference strain | $u_0$ | -0.44 |
| Rest length mismatch $v_0$ | $v_0$ | -0.25 |
| Dimensionless BAF-LEM2 surface density in siCtrl | $\rho_C$ | 13.3 |

TABLE III: Surface hydrogel parameters used in Fig. 3f and Extended Data Fig. 8i.

[1] W. Helfrich. Size distributions of vesicles : the role of the effective rigidity of membranes. *Journal de Physique*, 47(2):321–329, 1986.

[2] Jacob A Ross, Nathaly Arcos-Villacis, Edmund Battey, Cornelis Boogerd, Constanza Avalos Orellana, Emilie Marhuenda, Pamela Swiatlowska, Didier Hodzic, Fabrice Prin, Tim Mohun, Norman Catibog, Olga Tapia, Larry Gerace, Thomas Iskratsch, Ajay M Shah, and Matthew J Stroud. Lem2 is essential for cardiac development by maintaining nuclear integrity. *Cardiovascular Research*, 119(11):2074–2088, April 2023.

[3] Udo Seifert. Configurations of fluid membranes and vesicles. *Advances in Physics*, 46(1):13–137, 1997.

### Supplementary Note 2

##### CRISPR/Cas9-mediated generation of HCT116 knock-in lines

| Item ID | Name | Developer | Parent line |
|---|---|---|---|
| LR-20221209-11 | HCT116-BANF1_Cterm-mNG #B03 | Genome engineering facility, MPI-CBG | HCT116 |
| LR-20221209-4 | HCT116-LEMD2_Cterm-mNG #B03 | Genome engineering facility, MPI-CBG | HCT116 |
| LR-20221209-18 | HCT116_LMNA_Nterm-mNG #A06 | Genome engineering facility, MPI-CBG | HCT116 |

*Table. Cell lines used in the study*

| Gene | 5' guide RNA |
|---|---|
| hBAF | AGGCGTCGCACCACTCTCGA |
| hLEM2 | TCAGAGCGATAAGCCCCGGG |
| hLamin A/C | TGCAGCCGGGTGATGCGGGT |

*Table. Guide RNA sequences*

| Gene | primer forward (5'-) | primer reverse (5'-) |
|---|---|---|
| hBAF | GTTCCAGGTCTTCAGCCCTAA | CCACATCACTCGGGGATTGAG |
| hLEM2 | GCTTAGCTCATCCCATCCCC | GACCCTCCTCTCTGCTCTCA |
| hLamin A/C | AGAAGGTCTGAGGCAATGGGG | CTGATACCCCCACCATTCCT |

*Table. Primer sequences*

1. hBAF-mNeonGreen targeting construct (homololgy arms, **mNeonGreen CDS**)

CTGATCAAGATGACAACCTCCCAAAAGCACCGAGACTTCGTGGCAGAGCCCATGGGGGAGAAGC
CAGTGGGGAGCCTGGCTGGGATTGGTGAAGTCCTGGGCAAGAAGCTGGAGGAAAGGGGTTTTG
ACAAGGTGTGGGGTGGCTGCGTGTACCTAGTGCAAGCGGGGGGTGGAAGGGAAGTGATTCCAT
CTGCTGGGGGATGGACAGTAAGGTATAATCTGAAGAGGCTGCCAGAGCCTGGGCACCTGGTGG
AGAGGAGAGGGGGGCAAAACCCGCGCTGCTTCCTGGGCTTGTGTGCTCTGAATGGCACAGGAA
TGGCTGTCTTGCTCTTATCTCTCACTGAGCACTGAGCAGCACGCTCCTTCCTTTTCCCTGTTTTGC
AGGCCTATGTTGTCCTTGGCCAGTTTCTGGTGCTAAAGAAAGATGAAGACCTCTTCCGGGAATGG
CTGAAAGACACTTGTGGCGCCAACGCCAAGCAGTCCCGGGACTGCTTCGGATGTTTGAGGGAAT
GGTGTGATGCATTTCTGGGCGGAGGTGGCTCTGGCGGTGGCGGATCG**GTCTCTAAGGGCGAGG**
**AGGACAACATGGCTTCACTGCCTGCTACTCACGAGCTGCATATTTTCGGGTCAATTAACGGGGT**
**GGATTTTGACATGGTGGGCCAGGGCACCGGCAACCCAAATGATGGCTACGAGGAGCTGAATCT**
**GAAGTCTACAAAGGGCGACCTGCAGTTCAGCCCTTGGATCCTGGTGCCACACATCGGCTATGG**
**CTTTCACCAGTATCTGCCCTACCCTGACGGAATGAGCCCATTCCAGGCAGCCATGGTGGATGG**
**CTCCGGCTACCAGGTGCACCGGACCATGCAGTTTGAGGACGGCGCCTCTCTGACCGTGAACTA**
**CCGCTATACATACGAGGGCAGCCACATCAAGGGAGAGGCACAGGTGAAGGGAACCGGATTCC**
**CAGCAGATGGACCCGTGATGACCAACTCCCTGACAGCCGCCGACTGGTGCAGGTCTAAGAAG**
**ACATATCCCAATGATAAGACCATCATCAGCACCTTCAAGTGGTCCTATACCACAGGCAACGGC**
**AAGAGGTACAGATCCACCGCCAGGACCACATATACATTTGCCAAGCCCATGGCCGCCAACTAT**
**CTGAAGAATCAGCCTATGTACGTGTTCCGGAAGACCGAGCTGAAGCACTCTAAGACAGAGCTG**
**AACTTTAAGGAATGGCAGAAGGCTTTTACTGATGTGATGGGAATGGATGAACTGTATAAG**TGAT
GCTCTCTGGGAAGCTCTCAATCCCCAGCCCTCATCCAGAGTTTGCAGCCGAGTAGGGACTCCTC
CCCTGTCCTCTACGAAGGAAAAGATTGCTATTGTCGTACTCACCTCCGACGTACTCCGGGGTCTT
TTGGGAGTTTTCTCCCCTAACCATTTCAACTTTTTTTTGGATTCTCGCTCTTGCATGCCTCCCCCG
TCCTTTTTCCCTTGCCAGTTCCCTGGTGACAGTTACCAGCTTTCCTGAATGGATTCCCGGCCCCA
TCCCTCACCCCCACCCTCACTTTCAATCCGTTTGATACCATTTGGCTCCTTTTTTGGCAGAACAGT
CACTGTCCTTGTAAAGTTTTTTAGATCAATAAAGTCAGTGGCTTTCATGACTGGGCTTTGTGCACT
GAAAAGCAAGTGGGCTGGGAGTTGCCTCTCCTTCAGGGACTGACCATGTTTCCCTCACTCCTTAT
CAGTAGAGACCGAAACTGGCAGCTTCCCTTCCACTCCCA

2. hLEM2-mNeonGreen targeting construct (homololgy arms, **mNeonGreen CDS**)

GGCGGGGTACTGGAACTGGCTTGGCAAGGGCTGCGCAGGCAACAGGTCCCAGCAAGAGTCAGC
TAGCCTAGCACAGCCCTGCACACCTGGAGACCTGGGGGTGCTCCAGACACCTCGGCCCTTTAGC
TCCCTTTAATTGAATGTGTTTGGATCAGTGAAGGTTGAGGAATCATTTCTCTATGGCCCAAGACGT
TTTTCTCCTCTGCAGTTGTCATGTTAGTACCTGCCAGCTTTTCCTCTCTTACATAAATTCCATGCCA
GAGCCTGGAAATGTGTGCCCTTTGTAGGAGGGGCATCCACAGGCTGGCTCACCTCGGCAGTGC
CAGGCAGAGCCCCGTCCCTCTCATTGCAGGAGGCGCATGAAGCGTGTCTGGGACCGAGCTGTG
GAGTTCCTGGCCTCCAACGAATCCCGGATCCAGACGGAGTCCCACCGCGTTGCAGGAGAGGAC
ATGCTGGTGTGGAGATGGACTAAGCCCTCTTCCTTCTCTGACTCAGAGCGAGGCGGAGGTGGCT
CTGGCGGTGGCGGATCG**GTCTCTAAGGGCGAGGAGGACAACATGGCTTCACTGCCTGCTACTC**
**ACGAGCTGCATATTTTCGGGTCAATTAACGGGGTGGATTTTGACATGGTGGGCCAGGGCACCG**
**GCAACCCAAATGATGGCTACGAGGAGCTGAATCTGAAGTCTACAAAGGGCGACCTGCAGTTCA**
**GCCCTTGGATCCTGGTGCCACACATCGGCTATGGCTTTCACCAGTATCTGCCCTACCCTGACGG**
**AATGAGCCCATTCCAGGCAGCCATGGTGGATGGCTCCGGCTACCAGGTGCACCGGACCATGC**
**AGTTTGAGGACGGCGCCTCTCTGACCGTGAACTACCGCTATACATACGAGGGCAGCCACATCA**
**AGGGAGAGGCACAGGTGAAGGGAACCGGATTCCCAGCAGATGGACCCGTGATGACCAACTCC**
**CTGACAGCCGCCGACTGGTGCAGGTCTAAGAAGACATATCCCAATGATAAGACCATCATCAGC**
**ACCTTCAAGTGGTCCTATACCACAGGCAACGGCAAGAGGTACAGATCCACCGCCAGGACCAC**
**ATATACATTTGCCAAGCCCATGGCCGCCAACTATCTGAAGAATCAGCCTATGTACGTGTTCCGG**
**AAGACCGAGCTGAAGCACTCTAAGACAGAGCTGAACTTTAAGGAATGGCAGAAGGCTTTTACT**
**GATGTGATGGGAATGGATGAACTGTATAAG**TAAGCCCCGGGCGGGGACTTGTTCCCGGTCAGC
CTGTCCCAGGACAGCCCACTCCAGAGGCACAGGAGGGTCACCAGGCCTGCGGTGCTGAATTCA
CACTTGCCTTGACTTTCAGTCTCGTCCTGACACGATTCCAAAGGTCTGCCATACTCTTCAGAGGA
GTTCGGCTCGCAGGAAAATGTGGGCTTCCTTTAAAGGGAGGGGGAGAAGCCAGGGTGATAGCC
CAGACTCTGCCCTGTTGTGCTGGCAGAAACGGGGAGAAGGGTTGAGTTTTCATGAATAGAAAACA
AGAGCAGCAGCTGTCAGGCCCAGCTGACAAGCGTGGGGCTGGTGGGCTGGGGCATGGCCACG
CCAGGGAGTCTGTCACCTGGGAGGTGGGGCAGGGATGGTGTTTCGGGTGGCGGGATGGGGAG
CAGCTTGGTCCATGTGCCTTCAGGGGGTTGTTACTTGAGAGCTCCAGGGTGAAGGGAGAAAGCC
TGTGCTCTGGCTGGCTGGGCCTC

3. mNeonGreen-hLamin A/C targeting construct (homololgy arms, **mNeonGreen CDS**)

AGCCTCACGCAGTTAGGGGTGCGCTGGAGAGGGTGGGGCCCGACTCCGCCACACCCCAACGGT
CCTTCCCCCTCCTCACCACTCCCGCCCCCACCCCCAATGGATCTGGGACTGCCCCTTTAAGAGT
AGTGGCCCCTCCTCCCTTCAGAGGAGGACCTATTAGAGCCTTTGCCCCGGCGTCGGTGACTCAG
TGTTCGCGGGAGCGCCGCACCTACACCAGCCAACCCAGATCCCGAGGTCCGACAGCGCCCGGC
CCAGATCCCCACGCCTGCCAGGAGCAAGCCGAGAGCCAGCCGGCCGGCGCACTCCGACTCCGA
GCAGTCTCTGTCCTTCGACCCGAGCCCCGCGCCCTTTCCGGGACCCCTGCCCCGCGGGCAGCG
CTGCCAACCTGCCGGCCATG**GTCTCTAAGGGCGAGGAGGACAACATGGCTTCACTGCCTGCTA**
**CTCACGAGCTGCATATTTTCGGGTCAATTAACGGGGTGGATTTTGACATGGTGGGCCAGGGCA**
**CCGGCAACCCAAATGATGGCTACGAGGAGCTGAATCTGAAGTCTACAAAGGGCGACCTGCAG**
**TTCAGCCCTTGGATCCTGGTGCCACACATCGGCTATGGCTTTCACCAGTATCTGCCCTACCCTG**
**ACGGAATGAGCCCATTCCAGGCAGCCATGGTGGATGGCTCCGGCTACCAGGTGCACCGGACC**
**ATGCAGTTTGAGGACGGCGCCTCTCTGACCGTGAACTACCGCTATACATACGAGGGCAGCCAC**
**ATCAAGGGAGAGGCACAGGTGAAGGGAACCGGATTCCCAGCAGATGGACCCGTGATGACCAA**
**CTCCCTGACAGCCGCCGACTGGTGCAGGTCTAAGAAGACATATCCCAATGATAAGACCATCAT**
**CAGCACCTTCAAGTGGTCCTATACCACAGGCAACGGCAAGAGGTACAGATCCACCGCCAGGA**
**CCACATATACATTTGCCAAGCCCATGGCCGCCAACTATCTGAAGAATCAGCCTATGTACGTGTT**
**CCGGAAGACCGAGCTGAAGCACTCTAAGACAGAGCTGAACTTTAAGGAATGGCAGAAGGCTTT**
**TACTGATGTGATGGGAATGGATGAACTGTATAAG**GGCGGAGGTGGCTCTGGCGGTGGCGGATC
GGAAACTCCATCACAACGCCGGGCTACTCGGAGTGGAGCCCAAGCTAGTTCAACACCTCTCTCA

CCAACTCGGATTACCCGGCTGCAGGAGAAGGAGGACCTGCAGGAGCTCAATGATCGCTTGGCG
GTCTACATCGACCGTGTGCGCTCGCTGGAAACGGAGAACGCAGGGCTGCGCCTTCGCATCACC
GAGTCTGAAGAGGTGGTCAGCCGCGAGGTGTCCGGCATCAAGGCCGCCTACGAGGCCGAGCTC
GGGGATGCCCGCAAGACCCTTGACTCAGTAGCCAAGGAGCGCGCCCGCCTGCAGCTGGAGCTG
AGCAAAGTGCGTGAGGAGTTTAAGGAGCTGAAAGCGCGGTGAGTTCGCCCAGGTGGCTGCGTG
CCTGGCGGGGAGTGGAGAGGGCGGCGGGCCGGCGCCCCTGGCCGGCCGCAGGAAGGGAGTG
AGAGGGCCTGGAGGCCGATAACTTTGCCATAGTCTCCTCCCTCCCCGGAACTGCCCCCAGCGG
GTGACTGGCAGTGTCAAGGGGAATTGTCAAGACAGGACAGAGAGGGAAGTGGTGGTCTCTGGG
AGAGGGTCGG

**siRNA mediated knockdown of proteins**

| Item ID | Name | Target | Antisense (RNA) sequence 5' to 3' | Modifications (ie 3' dt dt) | Target sequence |
|---|---|---|---|---|---|
| LR-20231221-5 | siCtrl | Scrambled siRNA | UCUACGAUCGGAGAUUUGC | 3' dt dt | NA |
| LR-20231221-13 | siLEM2 | LEM2 | UACAUAUGGAUAGCGCUCC | 3' dt dt | ggagcgctatccatatgta |
| LR-20231221-12 | siBAF | Barrier to autointegration factor | AACGGAUUGAAAGUGAGGG | 3' dt dt | UTR of BAF |

*Table. siRNA sequences used*

### Extended Data Table 1

| Protein | Uniprot ID | Nucleoplasmic region | Copy number | LEM motif aa (SD) | IDR aa | Other structured domains aa SD) | Exclusively INM? | Number of residues belong to IDR | Number of residues belong to SD |
|---|---|---|---|---|---|---|---|---|---|
| LBR | Q14739 | 1-211 | 4.40E+05 | na | 61-211 | 1-60 | no | 66,440,000 | 26,400,000 |
| LEM2 | Q8NC56 | 1-212;398-503 | 8.20E+04 | 1-41 | 42-200 | 201-212 | yes | 13,038,000 | 13,038,000 |
| Emerin | P50402 | 1-224 | 1.30E+06 | 1-48 | 49-224 | na | mostly | 228,800,000 | 62,400,000 |
| Man1 | Q9Y2U8 | 1-470;651-912 | 8.70E+04 | 14-57 | 1-13;58-465;888-912 | | yes | 38,802,000 | 24,882,000 |
| LaminA | P02545 | 1-664 | 3.60E+05 | | 388-426;548-664 | | | 56,160,000 | 182,880,000 |
| LaminC | P02545 | 1-573 | 3.60E+05 | | 388-426;548-573 | | | 23,400,000 | 182,880,000 |
| LaminB1 | P20700 | 1-586 | 2.60E+06 | | 391-429;550-586 | | | 197,600,000 | 1,326,000,000 |
| LaminB2 | Q03252 | 1-620 | 8.30E+05 | | 405-460;582-620 | | | 78,850,000 | 435,750,000 |
| Sun1 | O94901 | 1-288 | 5.20E+04 | | 1-235 | | | 12,220,000 | 2,756,000 |
| Sun2 | Q9UH99 | 1-212 | 5.70E+04 | | 1-155 | | | 8,835,000 | 3,249,000 |
| Lap1 | Q5JTV8 | 1-333 | 3.70E+05 | | 1-333 | | | 123,210,000 | 0 |
| Lap2beta | P42167 | 1-409 | 7.50E+05 | 1-49 | 50-110;154-409 | | | 237,750,000 | 69,000,000 |
| Banf1 | O75531 | 1-89 | 2.80E+06 | | - | | no | 0 | 249,200,000 |

| Excluded | Protein | Remarks |
|---|---|---|
| | LEMD1 | meiosis specific |
| | Lap2alpha | Nucleoplasmic |
| | Lull1 | ER |
| | Ankle2 | ER |
| | Ankle1 | Nucleoplasmic |

| Group | Number of residues |
|---|---|
| IDR of LEM-proteins | 518390000 |
| IDR of other proteins | 566715000 |
| SD of BAF and LEM proteins | 249200192 |
| SD of Lamins | 21275.1 |
| SD of other proteins | 95213000 |

#### Supplementary Information

**Extended Data Table 1.** Top, copy numbers and domain distributions of INM proteins. Bottom, categorized protein domain distributions in the INM.

**Supplementary Video 1.** Binding of BAF to DNA in optical tweezers. BAF, labeled with DyLight 488 NHS Ester (appearing cyan in the movie), binds to λ-phage DNA under ~40 pN of tension. The DNA was initially tethered between two beads in a buffer channel, then transferred to a protein channel containing 5% labeled BAF(100nM) in constant trap position mode. In the movie, the two beads appear as dark spots, while the stretched DNA is seen as a bright cyan horizontal line.

**Supplementary Video 2. Binding of LEM2 to DNA in presence of BAF in optical tweezers.** LEM2, labeled with DyLight 650 NHS Ester (appearing magenta in the movie), binds to λ-phage DNA under ~40 pN of tension in the presence of BAF (unlabelled). The DNA was initially tethered between two beads in a buffer channel, then transferred to a protein channel containing unlabeled BAF(100nM) and 1% labeled LEM2(500nM). In the movie, the two beads appear as bright blobs (due to unspecific binding of LEM2 to the bead surface), while the stretched DNA is seen as a bright magenta horizontal line.

**Supplementary Video 3. LEM2 alone does not bind to DNA.** LEM2, labeled with DyLight 650 NHS Ester (appearing magenta in the movie), does not bind to λ-phage DNA on its own. The DNA was initially tethered between two beads in a buffer channel, then transferred to a protein channel containing 1% DyLight-labeled LEM2 (500 nM). In the movie, only two bright fluorescent blobs are visible, corresponding to nonspecific LEM2 binding on the bead surfaces, with no signal along the DNA. To confirm the DNA tether, the left bead (held in a steerable optical trap) is brought close to the right bead and then retracted, allowing the DNA to relax and re-stretch, as shown by the tension-distance profile.

**Supplementary Video 4. Formation of DNA-BAF co-condensates in optical tweezers.** DNA initially held at 40 pN tension in a buffer-only channel was transferred to a channel containing BAF (100 nM, 5% fluorescently labeled, appearing cyan in the movie) and 25 nM Sytox Orange (appearing yellow in the movie). The steerable trap (left trap) was then moved to introduce slack into the DNA, resulting in the formation of DNA-BAF co-condensates. These appear as three bright puncta in the movie. The two beads appear as bright yellowish-cyan blobs (due to unspecific binding of BAF/Sytox Orange to the bead surface).

**Supplementary Video 5. Formation of DNA-BAF-LEM2 co-condensates in optical tweezers.** DNA, initially held at 40 pN tension in a buffer-only channel was transferred to a channel containing BAF (100 nM, unlabeled), LEM2 (500 nM, 1% labeled, appearing magenta in the movie), and 25 nM Sytox Orange (appearing yellow in the movie). The steerable trap (left trap) was then moved to introduce slack into the DNA, resulting in the formation of DNA–BAF–LEM2 co-condensates, which appear as bright white puncta in the movie. The two trapped beads appear as bright blobs due to nonspecific binding of LEM2/BAF/Sytox Orange to the bead surfaces, predominantly from LEM2.

**Supplementary Video 6. Melting of ds-DNA outside co-condensate.** DNA with preformed DNA–BAF–LEM2 co-condensates was continuously stretched to disassemble the co-condensates in the presence of FUS (200 nM, GFP-tagged, appearing green in the movie). Magenta fluorescence corresponds to LEM2 (500 nM, 1% labeled), while BAF is unlabeled. The green FUS signal is observed only outside the magenta puncta representing the DNA–BAF–LEM2 co-condensates. The two large magenta blobs correspond to the trapped beads, due to nonspecific binding of LEM2 to the bead surfaces.
